# How learning to read visual Braille co-opts the reading brain

**DOI:** 10.1101/2024.05.08.593104

**Authors:** Filippo Cerpelloni, Alice Van Audenhaege, Jacek Matuszewski, Remi Gau, Ceren Battal, Federica Falagiarda, Hans Op de Beeck, Olivier Collignon

## Abstract

Learning to read assigns linguistic value to an abstract visual code. Whether regions of the reading network tune to visual properties common to most scripts or code for more abstracted units of language remains debated. Here we investigate this question using visual Braille, a script developed for touch that does not share the typical explicit shape information of other alphabets, yet maps onto the same phonology and lexicon as other more regular scripts. First, we compared univariate responses in visual Braille readers and a naïve control group and found that individually localized Visual Word Form Area (VWFA) was selectively activated for visual Braille when compared to scrambled Braille only in expert Braille readers. Multivariate analyses showed that linguistic properties can be decoded from Latin script in both groups and from Braille script in expert readers in an extended network of brain regions including the early visual cortex (V1), the lateral occipital region (LO), the VWFA and the left Posterior Temporal area (l-PosTemp). These results suggest that the tuning of an extended reading network to orthography relies more on the linguistic content of the script rather than their specific visual features (e.g. line junctions). Nevertheless, cross-scripts generalization was significantly lower than within-script decoding and failed to reveal common representations across Latin and Braille in experts in all regions except the l-PosTemp. These results suggest that V1, LO and VWFA encode orthographic representations in a script-specific manner, whereas l-PosTemp encodes abstracted linguistic information.

## Introduction

In vision, object recognition is modelled as a hierarchical process going from the detection of simple features to their progressive integration into more complex shapes (Riesenhuber & Poggio, 1999; Biederman, 1987; Krizhevsky et al., 2017). Shape, an arrangement formed by lines joined together in a particular way or by lines around the outer edge of an object, plays therefore a prominent role in human object recognition (Grill-Spector & Malach, 2004; Landau et al., 1988; Tarr & Bulthoff, 1998). The flexibility of shape processing in the human brain has allowed the development of written scripts, highly complex visual stimuli with symbolic meaning. Most written systems developed throughout history share basic shape features with natural objects (Changizi et al., 2006). Core features of visual object recognition may have been co-opted for written word recognition (Dehaene et al., 2005), exerting pressure for scripts to appear the way they do. Supporting this idea, basic sensitivity to Latin script letters seems to be present in untrained monkeys (Rajalingham et al., 2020), further illustrating how culturally invented scripts may have been optimized from the natural scaffolding of shape processing in the brains of primates.

In the human brain, visual scripts activate an extended network of brain regions when compared to non-linguistic visual stimuli (Fedorenko et al., 2011). It has been suggested that a predisposed sensitivity to shape features (e.g. line junctions) represents a proto-organization on which orthographic selectivity develops (Arcaro & Livingstone, 2017; Dehaene et al., 2005). The so called Visual Word Form Area (VWFA; Cohen et al., 2000, 2002), the major focus of word form selectivity in visual cortex, is a paradigmatic example of this debate. In terms of visual features, the presence of pre-existing shape selectivity in VWFA is one of the most often mentioned causes of the location of this region (Dehaene & Cohen, 2011; Dehaene-Lambertz et al., 2018). The importance of line-junctions for fluent reading (Bola et al., 2017), the partial overlap between VWFA and ventral visual areas responding selectively to line junctions (Szwed et al., 2011), equal responses to line drawings of objects and false fonts (Ben-Shachar et al., 2007), and comparable levels of activation for known and foreign characters (Vogel et al., 2012; Xue & Poldrack, 2007) all lead to the suggestion that VWFA, along with other regions located in the fusiform gyrus, might be involved in specific shape processing more generally (Roberts et al., 2013). Even without reading experience, the pre-existing shape selectivity in occipitotemporal cortex might be very well suited for learning to read, as demonstrated by the presence of basic sensitivity to Latin script letters in untrained monkeys (Rajalingham et al., 2020).

However, an alternative account of VWFA downplays the role of visual features and emphasizes the domain- and task-specific role of this region. It regards the integration of visual regions selective to orthography with downstream language regions involved in phonological and semantic processing as the key organisational feature driving selectivity for written language (Dȩbska et al., 2023; Price & Devlin, 2003, 2004, 2011). Structural and functional connectivity between VWFA and the traditional language network, even before infants learn to read may back up an interactive account of VWFA (Saygin et al., 2016; Li et al., 2020; Stevens et al., 2017). In apparent support of this view, studies have shown activations of VWFA for speech processing (Pattamadilok et al., 2019; Planton et al., 2019), but this has often been interpreted as an automatic activation of orthographic representations (Cohen et al., 2000, 2002). Further, some studies have shown that learning to read Korean Hangul or a script made of pictures of houses or faces (L. Martin et al., 2019; Moore et al., 2014; Xue & Poldrack, 2007) activates the reading network. However, those “atypical” scripts still rely on the processing of line-junctions and shape, potentially triggering their recruitment of the typical reading network. Studies in in sighted and blind people with tactile Braille reading further support the idea that VWFA encodes linguistic information more generally, rather than being selective to specific visual features (Büchel et al., 1998; Reich et al., 2011; Siuda-Krzywicka et al., 2016).

To the extent that brain preference for visual features or linguistic properties each have a separate role to play in the emergence of orthographic selectivity due to experience, the question emerges on how these factors might work together, and what impact each property might have. The findings with house- and face-based scripts and tactile Braille are consistent with the suggestion that the domain-specific task-related connectivity between visual and linguistic regions might determine in which precise region in the visual system a preference for the experienced stimuli will emerge (Saygin et al., 2016). Yet, even if true, this does not exclude a role for visual properties. For example, visual properties might determine which neural populations are implicated within the activated region. So, whereas a regular shape-based script, a fabricated face-based script, and even tactile braille might all activate the reading network, the way these scripts are represented in these brain regions might be different due to their visual properties. Such investigation requires an in-depth comparison of multiple scripts with methods that have the sensitivity to pick up fine-scale differences in activity patterns. A relevant study was published recently by (Zhan et al., 2023), who showed with high-resolution 7T imaging that the English and Chinese script both activate nearby and sometimes even overlapping brain regions in English/Chinese bilinguals, yet some small patches seem uniquely activated by only one of the two scripts. However, English and Chinese are both line-based scripts, and also differ in their mapping onto phonology and lexical entries - leaving open the possibility that nonvisual linguistic factors might explain the partial separation of the two scripts.

Overall, the literature shows that both visual and domain-specific nonvisual factors influence the activity of VWFA. An assessment of how they might work together in determining how orthographic selectivity emerges due to experience is missing, and visual Braille represents a unique opportunity to assess the contribution of visual and linguistic aspects. It is a visual script that does not share the most basic features of other scripts like lines, junctions, and clear outer shapes thought to be necessary for efficient reading (Bola et al., 2017) and for the location of the reading network. The letters are simple configurations of 1 to 6 dots, and arguably simpler than any other stimulus that is typically used to investigate cortical vision. Yet visual Braille maps onto the same phonology and lexicon as other scripts in the same language community, specifically the Latin script in a western country. Therefore, visual Braille provides a strong difference in visual properties but an optimal matching of nonvisual linguistic properties.

In the present study, we used fMRI to individually localize VWFA and to precisely evaluate whether the word form system activated for a Latin script is co-activated by reading visual Braille (Figure 1A). We then performed multivariate pattern analysis (MVPA) on VWFA, on several other key areas involved in visual processing, early visual cortex (V1) and lateral Occipital (LO), and on left Posterior Temporal area in the language network. Through MVPA we tested whether the same coding principles (using Representational Similarity Analyses, RSA) and neural populations (using cross-script decoding) underly the processing of both scripts (Figure 1B). We observed that most regions tested present overwhelming similarities in the coding principles used to processing Latin and Braille scripts, showing the flexibility of the visual system and the reading network for scripts made of atypical visual features like Braille. Importantly however, in most of the regions including VWFA, these coding principles were implemented in a script-specific manner showing intermingled but separate representations of both scripts. Together, these findings reveal how the word processing system implements the same function of reading applied on visual input with fundamentally different properties.

**Figure 1.**
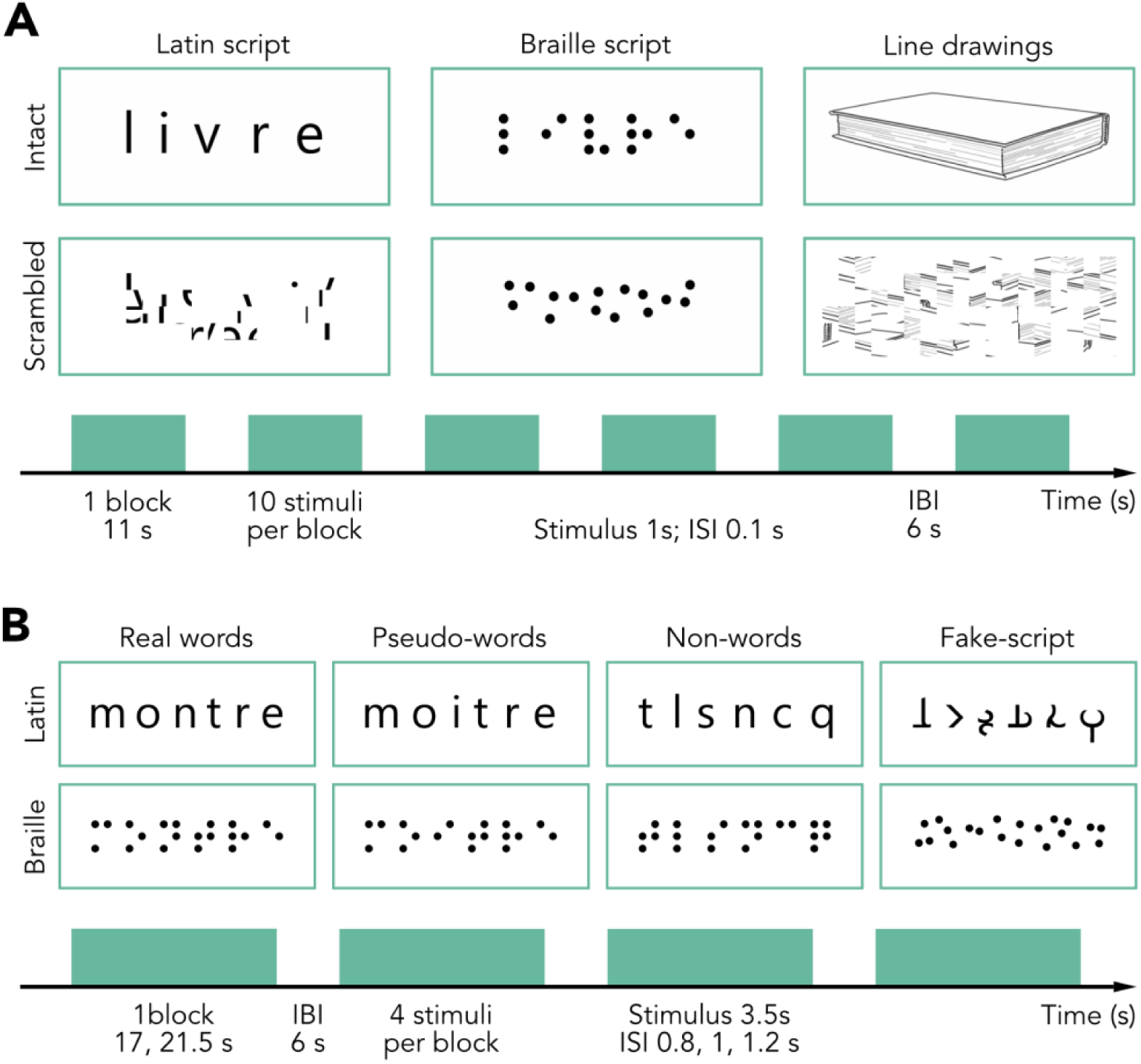
Experimental design and examples of stimuli for experiments 1 and 2. **(A)** Experiment 1: functional localizer. We presented six categories of stimuli (intact and scrambled Latin-script words, Braille words and line drawings) to participants in an fMRI block design. Each participant underwent two runs in which six blocks for each category were presented in pseudo-randomized order. Blocks lasted for 11 seconds and were separated by a fixed inter-block interval (IBI) of 6 seconds. Each block consisted of ten stimuli from one category, each for a duration of 1 second and with an inter-stimulus interval (ISI) of 0.1s. During the session, participants performed a 1-back task to detect the repetition of a stimulus. The target was present in 10% of the blocks. **(B)** Experiment 2: Coding of linguistic properties. Four different linguistic conditions (Real Words, Pseudo-Words, Non-Words, Fake Script) for each script (Latin, Braille) were used. We presented these categories of stimuli (intact and scrambled Latin-script words, Braille words, line drawings) to participants in a fMRI block design. Each participant underwent twelve sessions in total, six for each script (counterbalanced across subjects). In each run, three blocks for each category of one script were presented in a pseudo-randomized order. Each block lasted for 17 or 21.5 seconds, depending on the presence of a target, with a 6-second IBI separating the two blocks. Four stimuli from the same category were presented within a block, each for 3.5 seconds, with a jittered ISI of 0.8, 1, or 1.2 seconds. During the session, participants performed a 1-back task to detect the repetition of a stimulus. The target was present in 10% of the blocks. The position of the target, as well as jittered ISI, block order, and stimuli order, were counterbalanced across runs.

## Materials and Methods

### Participants

A total of twenty-three participants took part in the data acquisition. Experts in visual Braille reading are rare, so we knew a priori that the expert group could be small with number of participants mostly determined by availability. To overcome problems due to statistical power, we decided to recruit twice as many control participants. Furthermore, we performed extensive piloting in other participants to select two paradigms (localizer and multivariate imaging) that provide interpretable results in the main region of interest (visual word form area) in individual participants.

Data from five subjects were discarded due to incomplete acquisitions (three control subjects) or inability to functionally localize VWFA for the Latin-based script (two subjects: one control, one expert), resulting in eighteen subjects in the final sample. Six of them (6 females, mean age = 38.2 ± 11.4) were experts in reading Braille visually and twelve (10 females, mean age = 39.3 ± 10.4) were controls naïve to Braille alphabet. To determine visual Braille expertise, we asked expert participants to report their use of visual Braille as a declarative number of years spent working with / using Braille (mean expertise = 5.25 ± 5.43 years). Furthermore, we assessed their reading speed through a behavioural reading test. All expert readers were presented with the same list of 30 words and instructed to read as many words as possible out loud in the given time of 1 minute (mean number of words read = 12.2 ± 6.11 words; range = 6-23).

### Experimental design

Each participant took part in one experimental session, lasting maximum two hours in total, of which maximum 90 minutes inside the MRI scanner. Four participants did not complete the whole experiment in one session due to time constraints, and were invited to a second, shorter, session to conclude data acquisition. MRI acquisition followed the same procedure for all the participants. In the scanner, we acquired a structural image followed by two functional scans of the localizer experiment and twelve functional scans of the MVPA-oriented experiment, alternating between one run with Latin script stimuli and one with Braille script stimuli. The stimuli and the counterbalancing procedures used in each experiment are detailed below. All stimuli were presented in white on a black background, on a screen with 1920x1200 pixels of resolution, placed at 170 cm and viewed by the participant through a mirror placed on the headset.

### Functional localization experiment

We designed a functional localizer to individually define VWFA, selective for words, and bilateral Lateral Occipital area (LO), selective for object shape. We included six conditions of stimuli (Figure 1A): concrete French words expressed in the Latin alphabet (FW), the same French words presented in the Braille alphabet (BW), line drawings of the objects represented by the words (LD), and scrambled conditions for each of the ‘intact’ categories (scrambled Latin words, SFW; scrambled Braille words, SBW; scrambled line drawings, SLD). We selected 20 items for each condition. Latin words were selected from the Lexique 3.83 (New et al., 2005) and comprised of a length between 4 and 8 letters, and a frequency between 1 and 20 appearances per million words. Braille script words were the literal transcription of the Latin-based script words (*Code Braille Français Uniformisé Pour La Transcription Des Textes Imprimés (CBFU)*, 2008). Line drawings came from two images datasets (Brodeur et al., 2010; Rossion & Pourtois, 2004). If necessary, we manipulated the colour information by obtaining grayscale stimuli that presented only the essential lines of the drawings. Scrambled conditions were created through two different custom algorithms, applied to each ‘intact’ stimulus. Scrambling of Latin words and line drawings were the result of the re-arrangement of different parts of the corresponding ‘intact’ stimulus image in a grid-like manner. The scrambled Braille condition was created by randomizing the position of dots of each ‘intact’ Braille word. We used different scrambling methods to ensure the alteration of each word’s visual properties, line junctions in the case of Latin words and organization of dots in the case of Braille words. The localizer experiment included two block-design runs. In each run, participants were presented with 6 blocks of 10 stimuli presented for 1 second in white on a black background (Inter Stimulus Interval, ISI = 0.1s; Inter Block Interval, IBI = 6s). The order of the blocks was randomized, while ensuring that one condition was not presented in two consecutive blocks and that the first three blocks of a run were also presented at the end of the same run, in reverse order. Participants performed a 1-back task with a button press and were asked to detect an immediate repetition of the same stimulus (10% of the trials, so once per block). All stimuli subtended a frame of 520 x 208 pixels, corresponding to 6.1 and 2.5 degrees of visual angle in the MRI scanner, respectively.

### MVPA-oriented experiment

Inspired by Vinckier and colleagues (Vinckier et al., 2007), we designed four different conditions per script in a linguistic gradient fashion (Figure 1B). (1) Real Words (RW) possessed semantic, phonological, and orthographical properties. This set was composed of French words (New et al., 2005) six letters length, with high frequency (>1 per million), and high identifiability (definition of lemma > 90%). Additionally, we controlled bigram and trigram frequency (top 45^th^ percentile; Gimenes et al., 2020). (2) Pseudo-Words (PW) were based on Real Words by identifying stimuli from the French Lexicon Project (Ferrand et al., 2010). We ensured bi- and tri-grams composing Pseudo-Words to have the same frequency of the real word set. Additionally, we measured Levensthein distance (the number of additions, deletions, transformations necessary to transform one string into another) and only included elements with a distance of 2, to avoid excessive similarity and possible confusion with the subset of Real Words. The final set of Pseudo-Word stimuli possesses both orthographic and phonological regularities similar to Real Words, but with limited semantic meaning, and without a lexical entry. (3) For Non-Words (NW), we selected strings of consonants based on infrequent bigrams and trigrams (Gimenes et al., 2020), to create phonological distance between sets. We created the Braille conditions for all these categories by transliteration of the Latin conditions. (4) Fake Script (FS). For the Latin Fake Script, we transliterated the Non-Words set to a custom-made alphabet that maintained the same features of the original consonants, but in re-arranged line junctions. For Braille Fake Script, given the lack of evident shapes to re-arrange, we scrambled the dots arrangement as for the functional localizer.

For each condition, we selected twelve six-characters long different stimuli. This composition of stimuli, comparable with the one made by Vinckier and colleagues (Vinckier et al., 2007), created a linguistic gradient of words with meaning, phonology, and orthography, progressively stripped of one of those properties. The experiment was divided into twelve runs of blocked design, alternating between runs with Latin and with Braille stimuli (Figure 1B). Participants performed a 1-back task, reacting to stimuli repetitions (targets). The presentation of stimuli within a block and run was pseudo-randomized to balance the appearance of targets across runs, categories, and position of the stimulus repeated. A total of six targets in each run were presented (12.5% of all stimuli). We controlled for the order of blocks and, within a block, for the order of categories and single stimuli. All parameters were balanced, resulting in a fixed experiment design which participants performed almost in the same order, aside from the counter-balanced alternation of the starting script, either Latin or Braille. During each block, four stimuli were presented in white on a black background for 3.5s (ISI jittered = 0.8s, 1s, 1.2s; IBI = 6s), each block lasting ∼ 17-21.5s including target. The long stimulus presentation time is based on results from a previous study on experts’ habituation to Braille words (Coquillart et al., 2022). All stimuli subtended a frame of 500 x 200 pixels, corresponding to 5.9 and 2.4 degrees of visual angle in the MRI scanner, respectively.

### fMRI acquisition and pre-processing

This method section was automatically generated using bidspm (Gau et al., 2023). We tested all the participants using a 3T GE (Signa™ Premier, General Electric Company, USA, serial number: 000000210036MR03, software: 28\LX\MR, Software release: RX28.0_R04_2020.b) MRI scanner with a 48-channel head coil at the University Hospital of St Luc, Belgium. We acquired a whole-brain T1-weighted anatomical scan (3D-MPRAGE; 1.0 x 1.0 mm in-plane resolution; slice thickness 1mm; no gap; inversion time = 900 ms; repetition time (TR) = 2189,12 ms; echo time (TE) = 2.96 ms; flip angle (FA) = 8°; 156 slices; field of view (FOV) = 256 x 256 mm2; matrix size= 256 X 256). All functional runs were T2*-weighted scans acquired with Echo Planar and gradient recalled (EP/GR) imaging (2.6 x 2.6 mm in-plane resolution; slice thickness = 2.6mm; no gap; Multiband factor = 2; TR = 1750ms, TE = 30ms, 58 interleaved ascending slices; FA = 75°; FOV = 220 x 220 mm2; matrix size = 84 X 84). We extracted NIfTI files using MRIcroGL (https://github.com/rordenlab/MRIcroGL) and converted the content in BIDS format (Gorgolewski et al., 2016). We pre-processed the (f)MRI acquisitions with bidspm (Gau et al., 2023) (v3.1.0dev) using statistical parametric mapping SPM12 (version 7771) using MATLAB (R2021b) on a macOS computer (13.0.1). The pre-processing consisted of slice timing correction, realignment and unwarping, segmentation and skull-stripping, normalization to the Montreal Neuroimaging Institute (MNI) standard space, smoothing. Slice timing correction used the 28th slice as a reference (interpolation: sinc interpolation). Functional scans were realigned and unwarped using the mean image as a reference (SPM single pass with 6 degrees of freedom). The anatomical image was corrected for bias and segmented using a unified segmentation. The mean functional image from realignment was co-registered to the bias-corrected anatomical image (degrees of freedom: 6). The transformation matrix from this co-registration was applied to all the functional images. The deformation field was applied to all the functional images (target space: IXI549Space; interpolation: 4th-degree b-spline). Finally, we applied spatial smoothing using a 3D Gaussian kernel (functional localizer, FWHM = 6 mm; main experiment, FWHM = 2mm). These smoothed, normalised BOLD images served as inputs to statistical analyses.

### General Linear Models (GLMs)

For both experiments, we analysed fMRI acquisitions with bidspm (Gau et al., 2023) (v3.1.0dev) and SPM12 (version 7771) using the same MATLAB (R2021b) on a macOS computer (13.0.1). For the localizer experiment, at the subject level, we performed a univariate analysis with a linear regression at each voxel of the brain, using generalized least squares with a global FAST model and a high-pass filter (278 seconds cut-off). Image intensity scaling was done run-wide before statistical modelling such that the mean image would have a mean intracerebral intensity of 100. Block timings were convolved with SPM canonical hemodynamic response function (HRF) for the 6 stimuli categories (intact and scrambled Latin words, Braille words, Line drawings), in addition to ‘response’ and ‘target’ events of no interest and 6 head motion parameters (three translations, on the X, Y, Z axes, and three rotations) to account for motion artefacts. We computed four main contrasts: intact over scrambled Latin words (FW > SFW), Braille words over scrambled dots (BW > SBW), intact over scrambled line drawings (LD > SLD), intact and scrambled Latin words over rest (FW+SFW > rest). We used the [FW > SFW] and [BW > SBW] contrasts to independently localize VWFA in all the subjects, for Latin and for Braille. We tested the statistical significance of clusters identified in the [BW > SBW] contrast in or near the Latin-based VWFA, by applying Small Volume Correction (SVC) within a 10mm radius from the peak coordinates extracted from [FW > SFW]. We localized bilaterally the Lateral Occipital area (LO) by the [LD > SLD] contrast. Lastly, we localized V1 by the [FW+SFW > rest] contrast (see *Definition of the Regions of Interest* below). In the MVPA-oriented experiment GLM, we included 8 regressors of interest for stimuli conditions (Real Words, Pseudo-Words, Non-Words, Fake Script for both Latin and Braille scripts), and 8 regressors of no-interest (1 for targets; 1 for responses; 6 for head motion parameters). Contrast images were spatially smoothed using a 3D Gaussian kernel (FWHM = 2). The results of the main GLM were concatenated into 4D maps, to be included in the multivariate pattern analyses.

### Eye-movements control analysis

To exclude potential confounds of eye movements, we computed eye displacement and integrated it in a General Linear Model (GLM) as a regressor of no interest. We used bidsMReye (Gau & Cabee, 2023), a BIDS app relying on deepMReye (Frey et al., 2021) to decode eye motion from fMRI time series data, in particular from the MR signal of the eyeballs. Each run, the following values were computed: variance for the X gaze position, variance for the Y gaze position, framewise gaze displacement, number of X gaze position outliers, number of Y gaze position outliers, and number of gaze displacement outliers. Outliers were robustly estimated using an implementation of the median rule (Carling, 2000). We then used the newly assessed parameters to perform two additional subject-level GLM analyses. We added the eye displacement for each run to the regressors of no interest, and computed the same contrasts of the localizer GLM analysis. This analysis showed very similar findings as the main analysis without these regressors (Figure 2-3). Therefore, the definition of the regions of interest described in the next section refers to the GLM described at the beginning of the section “General Linear Model”.

### Definition of the Regions of Interest (ROIs)

To investigate the gradient of linguistic representations across low-level sensory and higher-order associative brain regions we defined multiple ROIs, namely: V1, LO, VWFA and left Posterior Temporal cortex (PosTemp), using intersections between neuroanatomy, existing literature and brain activation from our functional localizer. For VWFA, we started from the canonical coordinates (Cohen et al., 2000) of VWFA ([-45, -56, -16]) and we identified the peak activation in each subject corresponding to the [FW > SFW] contrast within a 10mm sphere from the canonical coordinates. We then expanded a progressively larger sphere from the individual peak coordinates and intersected this sphere with the activation map of the same [FW > SFW] contrast until the ROI comprised 100 voxels. Lastly, we constrained the expanded ROI to activation reported in the literature to ensure the precise localization of the area by overlapping the expanded mask with the functional activations downloaded from the neurosynth database (Yarkoni et al., 2011) (term: “visual words”). This procedure permitted us to control for the individual localization of VWFA and to avoid including voxels from adjacent but unrelated areas. To ensure the robustness of our results, we compared it to a classical definition of ROIs from neurosynth.org alone (entry keyword: “visual words”). We extracted the highest-responding 100 voxels from the activation map and performed the same analysis pipeline reported below in relation to our custom model. As reported in Figure 3-1 in the Appendix, we do not observe differences between the two methods. We continued the analyses with our individual definition of ROIs as we can better ensure the fine spatial localization of the areas (Fedorenko et al., 2010). We used the same approach to localize bilateral LO from the [LD > SLD] contrast. More specifically, we started from canonical coordinates (Left LO: [-47, -70, -5], right LO: [47, -70, -5]) and applied a 10mm sphere around each set of coordinates to identify individual peaks in the localized [LD > SLD] activation. We then expanded a sphere from the individual peaks until a minimum threshold of 100 voxels was reached. Lastly, we overlapped the expansion with the neurosynth mask for the term “object”. To identify early visual cortex (V1), we overlapped the bilateral brain mask from the Anatomy toolbox (Eickhoff et al., 2005) for bilateral V1 with a [FW + SFW > rest] contrast. For subsequent analyses on areas of the language network, we relied on parcels from Fedorenko and colleagues (Fedorenko et al., 2010). For each subject, we overlapped activations for the [FW > SFW] contrast to the different language parcels to identify brain regions that consistently responded to linguistic stimuli across subjects. Although in different subjects we could identify many areas of both the left and right hemispheres, we focused our analyses solely on the left Posterior Temporal region (l-PosTemp), as it was the only area to be present reliably in individual subjects, being identified in 15 out of the 18 total participants analysed, corresponding to more than 80% consensus in the activations.

### Multivariate Pattern Analyses

To test the gradient of linguistic representations we performed multivariate pattern analyses using the CoSMoMVPA toolbox (Oosterhof et al., 2016) combined with custom MATLAB code (version 2021b). We applied this method to all the regions described in the section “ROI definition”. To obtain decoding accuracies, we performed feature selection across individuals by restricting the number of features to correspond to the number of voxels in the smallest ROI extracted through each method. This resulted in the feature selection of 43 voxels for LO and VWFA, extracted through the functional localizers, 62 for l-PosTemp, and 81 for V1. We then trained a support vector machine (‘libsvm’) in a leave-one-run-out fashion. To identify an organization of linguistic content, we performed pair-wise decoding, obtaining six comparisons per script (RW-PW, RW-NW, RW-FS, PW-NW, PW-FS, NW-FS). To compare the representational patterns emerging from the pairwise decoding accuracies, we performed Representational Similarity Analysis (Kriegeskorte et al., 2008) (RSA). We created a Representational Dissimilarity Matrix (RDM) for each group and script, where the value in each cell is given by the binary decoding accuracy of the classifier for that comparison. Resulting RDMs represent the linguistic (dis)similarity between stimulus conditions. Additionally, we correlated each neural RDM to a model of linguistic distance between categories. We assigned a value to each category of stimuli based on the number of linguistic features possessed. Concretely, Real Words possess orthographic, phonological, and semantic properties (3 properties), Pseudo Words might possess some low semantic meaning but are void of a lexical entry (2 properties), Non Words have orthographic but low frequency (non-pronounceable) phonological elements (1 property), and Fake Script has none. The resulting distances were computed by subtracting the number of features shared and normalized by dividing for the number of properties of the Real Words. We performed the correlations between neural matrices by applying non-parametrical statistical tests (Stelzer et al., 2013; Mattioni et al., 2020). When comparing two conditions, we first computed the correlation for each individual of the first condition to the group average of the second condition. For conditions from the same group (e.g. experts-Latin and experts-Braille), we averaged the group RDM of the second condition by excluding the subject being compared. We then averaged the individual correlations. Next, we performed the same correlation in the other direction, correlating each individual RDM from the second condition to the first condition averaged (with the same precaution), and then calculated the mean of these correlations. We then averaged the correlations obtained from the two directions, obtaining the correlation between the two matrices. Lastly, to assess the generalizability of the decoding from one script to the other, we performed cross-script decoding. We limited our analyses to visual Braille experts, the only group showing significant decoding for both scripts in our ROIs. For each script, we trained our multi-voxel classifier on pair-wise linguistic differences within a script and tested the classification on the other script (e.g., train on the difference between Braille Real Words and Pseudo-Words and test on the same difference within the Latin script). We performed this classification reversely for both alphabets and averaged across the two directions. To control for possible collinearity between the linguistic and the visual properties of the two scripts, we computed additional models based on the difference in pixels between two stimuli of the same script, comparing each stimulus to every other based on the number of overlapping white pixels between the images. We then averaged the pixel differences within one condition and one script to obtain two 4-by-4 matrices representing the visual differences across categories. We then used the same procedure described above to compute partial correlations between one neural RDM and one model (linguistic or visual), controlling for the effects of the other model (visual or linguistic). We limited this analysis to VWFA, as it was the main focus of our study. The results are presented in Figure 3-2. We observed that both visual and linguistic factors seems to contribute to the brain representations of scripts in VWFA, and that these factors are modulated by experience, as we do not observe correlations between the brain representations of Braille in the non-experts for neither visual nor linguistic models.

### Statistical analyses

Given the inter-subject variability in the location of functional regions, all our analyses were carried out in individually localized ROIs (see ROI definition). To test between groups differences in VWFA activation for the [BW > SBW] contrast, we applied a Chi-square test on the number of participants who showed activation for such contrast. To test statistical significance of overall decoding accuracies we performed one-tail t-tests against chance level (mu = 0.5). To test group-script differences, we performed two-tailed t-tests with other script-group pairs to test for significant differences in the decoding accuracy. If the pairs referred to the same group (e.g. Latin, experts and Braille, experts), we performed paired t-tests. In all the t-tests, we additionally calculated Cohen’s D effect size measure. To test whether cross-script decoding was significantly different from within-script decoding, we averaged the decoding across all pairs of linguistic conditions and also across both scripts for the within-script decoding, and then used a t-test across participants to compare the cross-script and the within-script decoding. We applied repeated measures ANOVA (6 stimuli conditions * 2 groups) to test for pairwise decoding accuracy differences between groups, within each script and each ROI. Effect size was estimated through partial eta-squared (η²_p_) measures. Effect sizes for bilateral LO, V1, and pSTS, can be found in Figures 4-1, 4-3, 5-1, 6-1. Additionally, we applied post-hoc t-tests to assess which conditions drove the effect of conditions (Figures 4-2, 4-4, 5-2, 6-2). Correlations between RDMs were tested through non-parametric statistics. For each correlation between neural matrices, we constructed a null distribution by creating 10000 matrices with shuffled labels and calculating the corresponding correlations. We shuffled both correlation directions and then averaged the distributions for each direction separately and then between directions. The result was a null distribution of 10000 values representing correlations between conditions with shuffled labels. We then compared the correlations obtained from the matrices with intact labels to said distribution and extract a measure of the significance. In the case of correlations between neural-RDMs and our linguistic distance model, we only shuffled labels regarding each subject of one group and the model itself. The final result was again a null distribution of 10000 values representing correlations between conditions with shuffled labels. We then applied the same steps to obtain p-values relative to each observed correlation. The p values in the rmANOVA post-hoc t-test were corrected using Bonferroni correction. All p values in the MVPA analyses were corrected for multiple comparisons with false discovery rate (FDR) corrections.

## Results

### The Visual Word Form Area of expert readers is selective to visual Braille

The first major question in our study was whether the VWFA as activated by the Latin script is also activated by visual Braille in experts. In all eighteen subjects that took part in the experiment and showed a VWFA for the Latin script (six expert Braille readers, twelve control participants), we performed a General Linear Model (GLM) on the whole-brain level and contrasted univariate activations for intact over scrambled Latin script words (FW > SFW). We then contrasted, in the ROI identified, activity for intact over scrambled Braille words (BW > SBW). In the expert group, individual clusters appeared in the [BW > SBW] contrast, in similar sites to the response to Latin-based stimuli (Figure 2). We performed Small Volume Correction at an individual level by drawing a 10mm radius sphere in the [BW > SBW] contrast around the peak coordinates in VWFA separately and individually identified through the [FW > SFW] contrast. A significant selective response for Braille (BW > SBW) in 5 of the 6 experts emerged in VWFA defined from Latin-based French (Figure 2-1). In the control group, only 1 out of 12 subjects showed a small significant cluster (Figure 3-1). The proportion of activations in participants was significantly different between the two groups (Chi-square_(1)_ = 7.03, p = 0.008), showing that people who can read Braille through the visual modality activate VWFA, regardless of the lack of visual features present in canonical scripts (Changizi et al., 2006). Next, we investigated whether a significant preference for intact over scrambled Braille was also present in our expert group as a whole (Figure 2-2), through paired t-tests on the mean activation for intact or scrambled Braille stimuli. We found a significant difference between intact and scrambled Braille in VWFA defined through the Latin-based script (t_(5)_ = 4.45, p = 0.006). This finding is in line with the aforementioned observation of significant clusters of Braille selectivity in/near Latin-based VWFA. Our results were not influenced by including regressors controlling for eye movements (Figure 2-3) through a decoding of eye motion from fMRI time series using bidsMReye (Gau & Cabee, 2023) and deepMReye (Frey et al., 2021).

**Figure 2.**
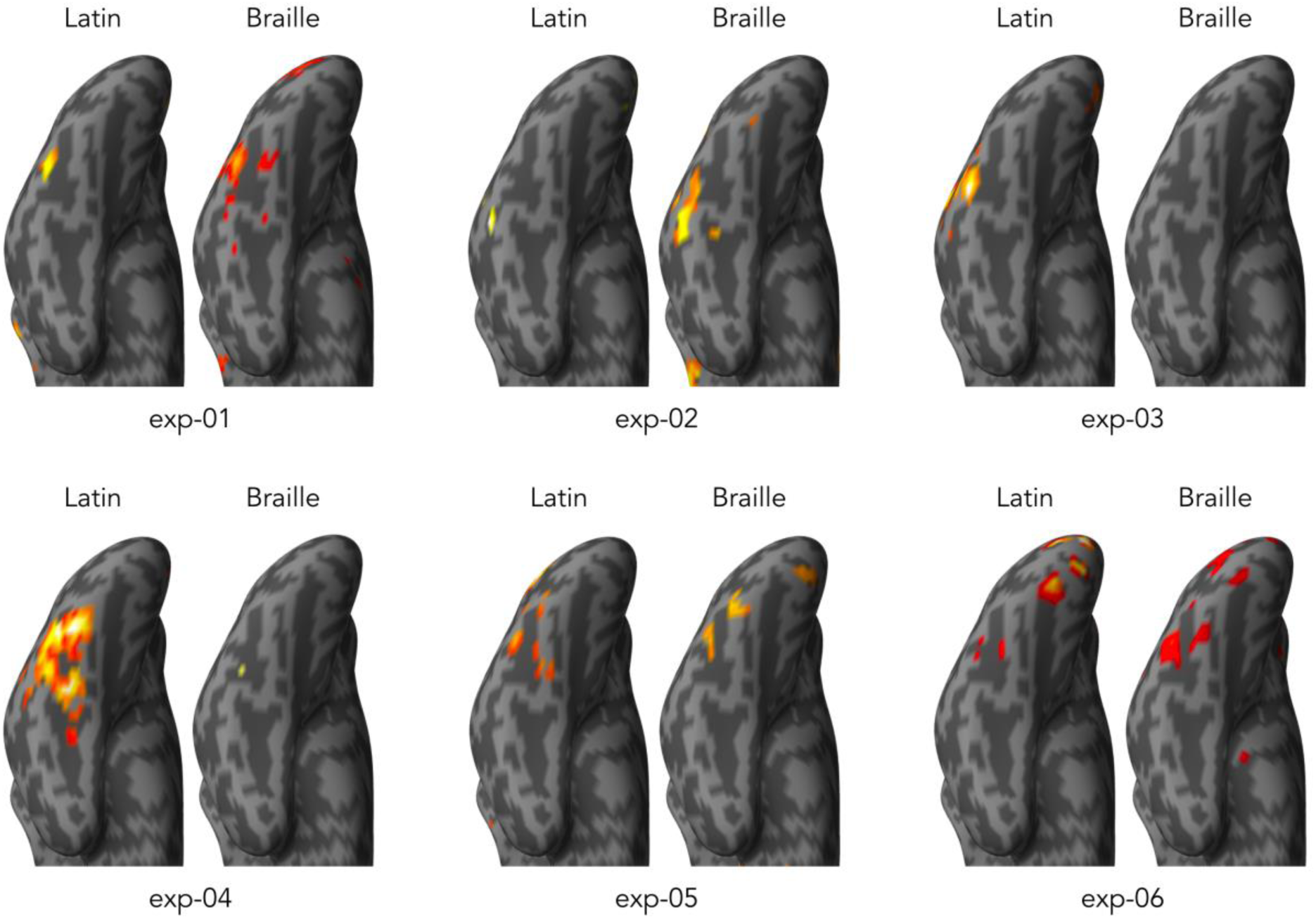
Visual Word Form Area’s univariate response to visual Braille in experts. Results of whole-brain analysis in each expert participant for [FW > SFW] and [BW > SBW] contrasts. For each expert visual Braille reader, contrasts for intact over scrambled Latin stimuli (left side of each participant) and intact over scrambled Braille stimuli (right side of each participant). All results are p_uncorr_ = 0.001 for visualization purposes. Through Small Volume Correction, we identified significant activation for intact over scrambled Braille in sites that localize intact over scrambled Latin stimuli in 5 experts (all except exp-03). Details of the areas localized, with peaks of activity, are available in Figure 2-1. Such selectivity is not present in other areas (Figure 2-2) and is not to be attributed to eye movements (Figure 2-3).

### Linguistic properties can be decoded in VWFA when a script is known, regardless of its visual features

To investigate whether the multivariate profiles of VWFA organization was based on linguistic content, we trained a support vector machine (SVM) to classify pairs of linguistic conditions. These conditions (Real Words, Pseudo-Words, Non-Words, Fake Script) differed in the number of linguistic properties presented (see “Methods” section). We decoded pairwise comparisons of linguistic conditions within each script, on an individual level in regions defined by selectivity for intact over scrambled Latin words (Figure 3A). In all the script-group pairs where the stimuli come from a known script (experts seeing Latin-based stimuli, experts seeing Braille stimuli, controls seeing Latin-based stimuli), pairwise decoding accuracy (DA) was on average strongly above chance level (Latin script, expert group: DA = 0.72, t_(5)_ = 5.850, p_FDR_ = 0.002, D = 2.3; Braille script, expert group: DA = 0.67, t_(5)_ = 8.386, p_FDR_ < 0.001, D = 3.4; Latin script, control group: DA = 0.72, t(11) = 5.347, pFDR < 0.001, D = 1.6). In striking contrast, visual Braille decoding in VWFA was not significant in the control group decoding (DA = 0.51, t_(11)_ = 0.61, p_FDR_ = 0.36). Additionally, we performed two-tailed t-tests on the decoding accuracies (in case of comparisons within a group, we performed paired t-tests) and found significant differences between known and unknown scripts (Latin, experts – Braille, controls: t_(8.6)_ = 4.73, p_FDR_ = 0.002, D = 2.5; Braille, experts – Braille, controls: t_(12.1)_ = 5.95, p_FDR_ < 0.001, D = 2.7; Latin, controls – Braille, controls: t_(11)_ = 4.88, p_FDR_ = 0.0012, D = 1.7, Figure 3A). In contrast, there were no significant differences between conditions across known scripts (p_FDR_ > 0.36). These results suggest that the VWFA of experts encodes the differences between linguistic properties in visual Braille as they do, and as controls do, in their native Latin script, while no such representation is detected for visual Braille in control participants.

**Figure 3.**
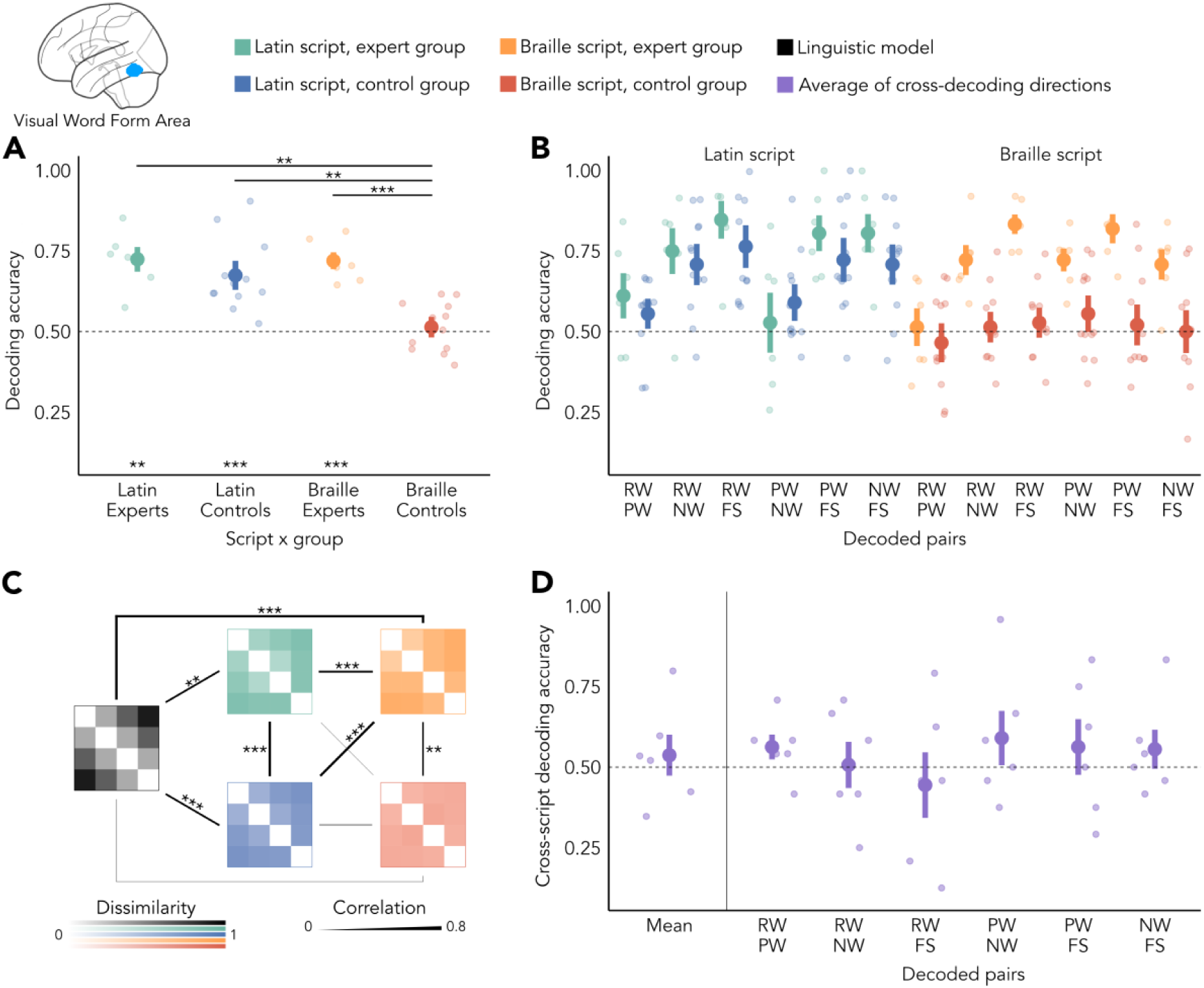
Multivariate representation of linguistic properties in the Visual Word Form Area (VWFA). **(A)** Average of pairwise decoding of conditions that differ in linguistic properties. In all the known scripts (Latin and Braille in experts, Latin in controls), we found significantly above-chance decoding and significant differences with the unknown and non-decodable condition of Braille presented to a control group. **(B)** Pairwise decoding within scripts. While experts and controls showed no group difference for their common native script, they showed a group difference for Braille, with experts showing a pattern similar to the one of Latin script. Details of post-hoc analyses can be found in Figure 3-2. **(C)** Representational similarity analysis (RSA) between neural representations (green, orange, blue, red) and with a linguistic model (black). Strength of the correlation is represented by the thickness of the bar connecting two patterns. Organizational patterns of known scripts correlated strongly with each other and with a theoretical model of linguistic distance. A script unknown to the reader (Braille for the control group) does not correlate with the model nor with patterns for Latin script. A significant correlation is present between the neural patterns of Braille. **(D)** Cross-script decoding. The mean of cross-decoding across all pairs of linguistic conditions did not show significant above-chance decoding, and neither did the individual pairs. RW = Real Words; PW = Pseudowords; NW = Nonwords; FS = Fake Script. In all the figures, stars represent significance (* p ≤ 0.05, ** p ≤ 0.01, *** p ≤ 0.001). Error bars represent standard error. Region of interest comes from an example subject. Each ROI was individually localized from peak activations at given contrasts (Table 3-1).

### Linguistic regularities in VWFA are organized similarly across scripts

Next, we investigated gradients of script-specific representations. We looked in detail at the same pairwise comparisons described in the section above to explore the different relationships between linguistic conditions. We performed within script repeated measures ANOVAs to identify group effects (differences between expert and naïve subjects), effects of the differences in linguistic properties (differences between decoding pairs), and interactions between those two variables (influence of the expertise on the task). In the case of the Braille script, we found an effect of group (F_(1)_ = 30.825, p < 0.001, η²_p_ = 0.66), as well as an effect of linguistic properties (F_(5)_ = 6.093, p < 0.001, η²_p_ = 0.25) and an interaction between the two factors (F_(5)_ = 3.195, p = 0.011, η²_p_ = 0.14), as shown in Figure 3B. These results indicate that the two groups process Braille differently. While VWFA can identify differences between conditions in the case of expert readers, this is not possible in the control group. In contrast, the analysis of the Latin script only showed an effect of the linguistic properties (F_(5)_ = 11.240, p < 0.001, η²_p_ = 0.4), but no difference between groups (F_(1)_ = 0.888, p = 0.36) and no interaction (F_(5)_ = 0.863, p = 0.51). Post-hoc t-tests on the conditions showed that the classification of Real Words against Pseudo-Words (RW-PW) is different (less accurate) from all the other pairs, except the pseudo- and - non-words pair (RW-NW: p_Bonf_ = 1, all other p_Bonf_ < 0.024). A full table of the post-hoc results is available in Figure 3-2. Differences between conditions show a sensitivity to the linguistic properties of a script such as enhanced linguistic distances triggers enhanced decoding accuracy for all known scripts (but not for Braille in non-experts). The lack of group differences for the Latin script shows that this sensitivity is induced by differences in expertise in a given script. When groups are compared on their native script (levelling the effect of training), no differences are found. To compare the organizational patterns between scripts and groups, we performed Representational Similarity Analysis (RSA). We built dissimilarity matrices using the pairwise decoding accuracies of each script-group pair (Figure 3C), as well as for a model of linguistic distance that captures the number of linguistic properties that are different between two conditions (with Real Words and Fake Script being most different). We computed the Pearson’s correlations between the values in these dissimilarity matrices, and determined their significance through non-parametric permutation analyses (Mattioni et al., 2020; Stelzer et al., 2013) (see “Methods”). We found high correlations for known scripts, both between patterns of neural activity (Latin, experts – Latin, controls: r = 0.58, p_FDR_ < 0.001; Braille, experts – Latin, controls: r = 0.58 p_FDR_ < 0.001) and with the linguistic model (Latin, experts – Model: r = 0.43, p_FDR_ = 0.005; Braille, experts – Model: r = 0.59, p_FDR_ < 0.0001; Latin, controls – Model: r = 0.45, p_FDR_ < 0.0001). Correlations with the dissimilarity matrix for Braille in the control group were mostly absent (all p_FDR_ > 0.20), with the exception of a correlation with the matrix for Braille stimuli in expert readers (r = 0.33, p_FDR_ = 0.002). Overall, these correlations further support the view that Braille script in experts is represented in the VWFA in a similar way as Latin-based stimuli and that the type of representation is related to a theoretical model of overlapping linguistic properties inspired by previous findings (Vinckier et al., 2007).

### Linguistic representations in VWFA are not generalized across scripts

Since linguistic conditions in Braille readers could be decoded in both scripts (Braille, Latin) we looked at possible generalizations between scripts. We performed cross-script decoding to see if the SVM classifier trained to distinguish linguistic conditions in one script would be able to decode similar conditions in the other script (Figure 3D). A significant result would indicate that the neural patterns associated with the different conditions partially abstract from the visual features intrinsic to each script and would represent linguistic properties independently of the script. We limited this analysis to the expert group since it was the only group showing significant decoding in each script separately. The average cross-script decoding accuracy was not significantly higher than chance (DA = 0.54, t_(5)_ = 0.58, p_FDR_ = 0.40). This result was maintained in the comparisons of all pairs of linguistic conditions, where no pair of stimuli showed significant cross-script decoding (highest DA value: RW – PW, DA = 0.56, t_(5)_ = 1.62, p_FDR_ = 0.41). Moreover, we compared the average decoding accuracy within scripts to the cross-decoding accuracy. We found a significant difference (t_(5)_ = 3.15, p = 0.025), providing positive evidence that the within-script decoding relies upon neural representations that are separated to such a degree that cross-script decoding is affected strongly.

### Shape-sensitive Lateral Occipital (LO) does not preferentially process Braille, but represents the differences in linguistic properties

We contrasted fMRI activation for intact and scrambled line drawings to identify individual bilateral Lateral Occipital (LO) sites as two regions of interest (ROIs), one for each hemisphere. To probe for extended plasticity effects of visual Braille in other areas commonly associated with shape, we then performed paired t-tests on the mean activation for intact or scrambled Braille stimuli in those ROIs and found no overall difference in LO (left-LO: t_(5)_ = 1.50, p = 0.19; right-LO: t_(5)_ = 1.86, p = 0.12). This result was not dependent on eye movements (Figure 2-3). We then performed the same Multivariate Pattern Analyses applied to VWFA (see “Methods”). In summary, both hemispheres showed similar results as to VWFA. We found significant decoding of linguistic properties (Figs. 4A and 4B) only in scripts known by the groups of participants (left-LO: all p_FDR_ < 0.006; right-LO: all p_FDR_ < 0.011) and not for Braille processed by the control group (all p_FDR_ > 0.10). We also observed statistical differences between known scripts and Braille for controls (all p_FDR_ < 0.031). In left-LO, we additionally observed differences between Braille in experts and the native script (both p_FDR_ < 0.033). Looking within each script, we found effects in line with VWFA. In Braille, we found group differences (both p ≤ 0.002), but no effects of linguistic properties (both p ≤ 0.11) nor interactions (both p > 0.23). Within Latin, both hemispheres presented a main effect of linguistic properties (both p < 0.001), but no group differences (both p > 0.36). In left-LO, we found an interaction between groups and linguistic properties (left-LO: F_(5)_ = 2.959, p = 0.018; right-LO F_(5)_ = 0.895, p = 0.49). Detailed statistical tests and results are available in Figure 4-1 and 4-3. These results indicate that experts’ LO represents Braille stimuli, although differently from the way Latin-based stimuli are processed. Representational Similarity Analysis (RSA) on the patterns of pairwise decoding accuracies (Figure 4C) showed strong correlations for known scripts, both between the neural RDMs (left-LO: all r > 0.55, all p_FDR_ < 0.001. Right-LO: all r > 0.64, all p_FDR_ < 0.001) and with the linguistic model (all p_FDR_ ≤ 0.002). In the left-LO of experts, the correlation between Braille-experts RDM with the model was close to significance (r = 0.34, p_FDR_ = 0.052). Braille representations in controls did not correlate with other neural RDMs or with the model (all p_FDR_ > 0.07). No significant cross-script decoding was observed (left-LO: DA = 0.59, t_(5)_ = 1.56, p_FDR_ = 0.16; right-LO DA = 0.58, t_(5)_ = 2.35, p_FDR_ = 0.10). Investigating the differences between within scripts and cross-script decoding, we found a significant difference (t_(5)_ = 3.45, p = 0.018) in the case of right-LO, and an almost-significant trend in the case of left-LO (t_(5)_ = 2.54, p = 0.051). Finer comparisons were mostly consistent with a lack of cross-decoding and with what we found in VWFA: except for Real Words contrasted to Pseudo-Words in the left LO (DA = 0.56, t_(5)_ = 4.39, p_FDR_ = 0.025), we observed no significant cross-decoding being above chance level in each pair-wise instances tested.

**Figure 4.**
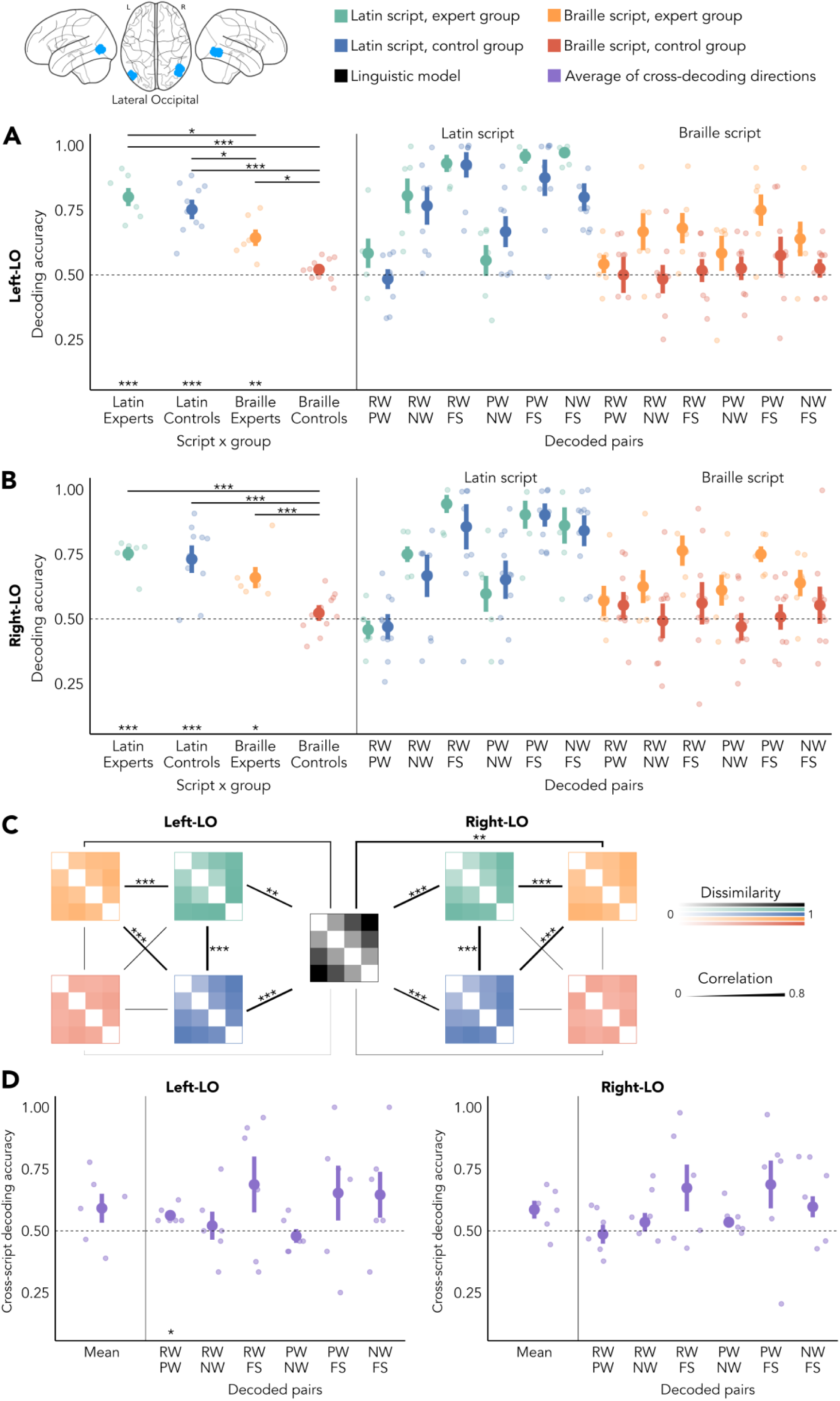
Multivariate representation of linguistic properties in left and right Lateral Occipital area (LO). **(A)** Pairwise decoding of conditions that differ in linguistic properties in left-LO, average (left) and individual pairs within script (right). Scripts known by the participants showed above-chance decoding accuracy and significant differences with the Braille-experts pair. Significant differences also emerged between the Braille-experts pair and Latin script in both groups. Individual pairs highlighted similar trends within Latin script but different ones within Braille, with experts showing a pattern similar to the one for Latin. **(B)** Pairwise decoding of conditions that differ in linguistic properties in right-LO, average (left) and individual pairs within script (right). Known scripts by the participants showed above-chance decoding accuracy and significant differences with the Braille-experts pair. Individual pairs highlight similar trends within Latin script but different ones within Braille, with experts showing a pattern similar to the one for Latin. Details of post-hoc analyses can be found in Figures 4-2 and 4-4. **(C)** Representational similarity analysis (RSA) between neural representations (green, orange, blue, red) and with a linguistic model (black). Strength of the correlation is represented by the thickness of the bar connecting two patterns. In left-LO, patterns of known scripts correlate strongly with each other. While Latin script patterns also correlate with the model, Braille representations in experts show an almost-significant trend (p = 0.052). In right-LO, only neural RDMs relative to known scripts show correlations between themselves and with the linguistic model. **(D)** Cross-script decoding. The mean of cross-decoding across all pairs of linguistic conditions did not show significant above-chance decoding. In right- LO (right plot), no individual comparison, nor the mean, can be decoded across scripts. RW = Real Words; PW = Pseudowords; NW = Nonwords; FS = Fake Script. In all the figures, stars represent significance (* p ≤ 0.05, ** p ≤ 0.01, *** p ≤ 0.001). Error bars represent standard error. Region of interest comes from an example subject. The full list of statistical test for left-LO and right-LO can be found respectively in Figures 4-1 and 4-3.

### Multivariate coding of linguistic properties of visual Braille extends to early visual cortex

We further extended our analysis of the organizational pattern found in VWFA and LO, to see if widespread visual Braille processing involves earlier stages of visual perception. To identify the region of interest, we relied on the bilateral early visual cortex (V1) mask from Anatomy toolbox (Eickhoff et al., 2005) and overlapped it with an independent contrast to determine overall visual activity as extracted from the General Linear Model (GLM) for each participant (intact and scrambled Latin words over rest, see “Methods”). Paired t-tests on the mean activation for intact or scrambled Braille stimuli showed no overall difference (t_(5)_ = 3.67, p = 0.07). This result was confirmed also when we controlled for eye movements (Figure S2). On the multivariate level (Figure 5A), we found above chance decoding for all scripts (all p_FDR_ <= 0.02). Moreover, decoding accuracies of known scripts differed from participants seeing an unknown script (all p_FDR_ < 0.042). Then, we performed repeated measures ANOVAs for individual scripts. Braille script (Figure 5B) showed a significant main effect of group (F_(1)_ = 10.611, p = 0.005), and of linguistic differences (F_(5)_ = 5.818, p < 0.001), as well as an interaction etween the two (F_(5)_ = 3.011, p = 0.015). Only experts showed distinctions in the neural patterns associated with different linguistic properties of the stimuli. Expectedly, Latin script only showed a main effect of the linguistic properties compared (F(5) = 7.475, p < 0.001), with no differences between groups (F_(1)_ = 0.007, p = 0.94). Post-hoc tests showed that the difference between Real and Pseudo-Words, the classification of stimuli that share the most linguistic features, is significantly different from (lower than) all the other comparisons of conditions (all p_Bonf_ < 0.004). These results indicate that, while Latin script is processed similarly in the two groups, a group difference is present in Braille, surely due to the difference of Braille expertise between our participants. Neural pattern of linguistic distances (Figure 5C) showed significant correlations between the neural RDMs for known scripts (all p_FDR_ < 0.001) and between scripts in the control group (Latin, Controls – Braille, Controls, r = 0.22, p_FDR_ = 0.021), suggesting that the processing of Braille in the expert group is similarly organized to the processing of the Latin script in both groups. The linguistic model correlated significantly with representations of Latin script stimuli (all p_FDR_ < 0.026) but, in contrast to the findings in all other ROIs, not with representations for Braille stimuli (all p_FDR_ > 0.09). Lastly, we did not find any significant cross-script decoding in V1 (Figure 5D), neither in the average of the comparison pairs (DA: 0.49, t_(5)_ = -0.36, p_FDR_ = 0.86) nor in the single pairs (most significant value: RW – FS, DA = 0.53, t_(5)_ = 0.72, p_FDR_ = 0.71), hinting that the distinct low-level visual features of the two scripts result in separate representations. Detailed statistical tests and results are available in Figure 5-1. Finally, there was a significant difference between within-script decoding and cross-script decoding (t_(5)_ = 3.28, p = 0.022).

**Figure 5.**
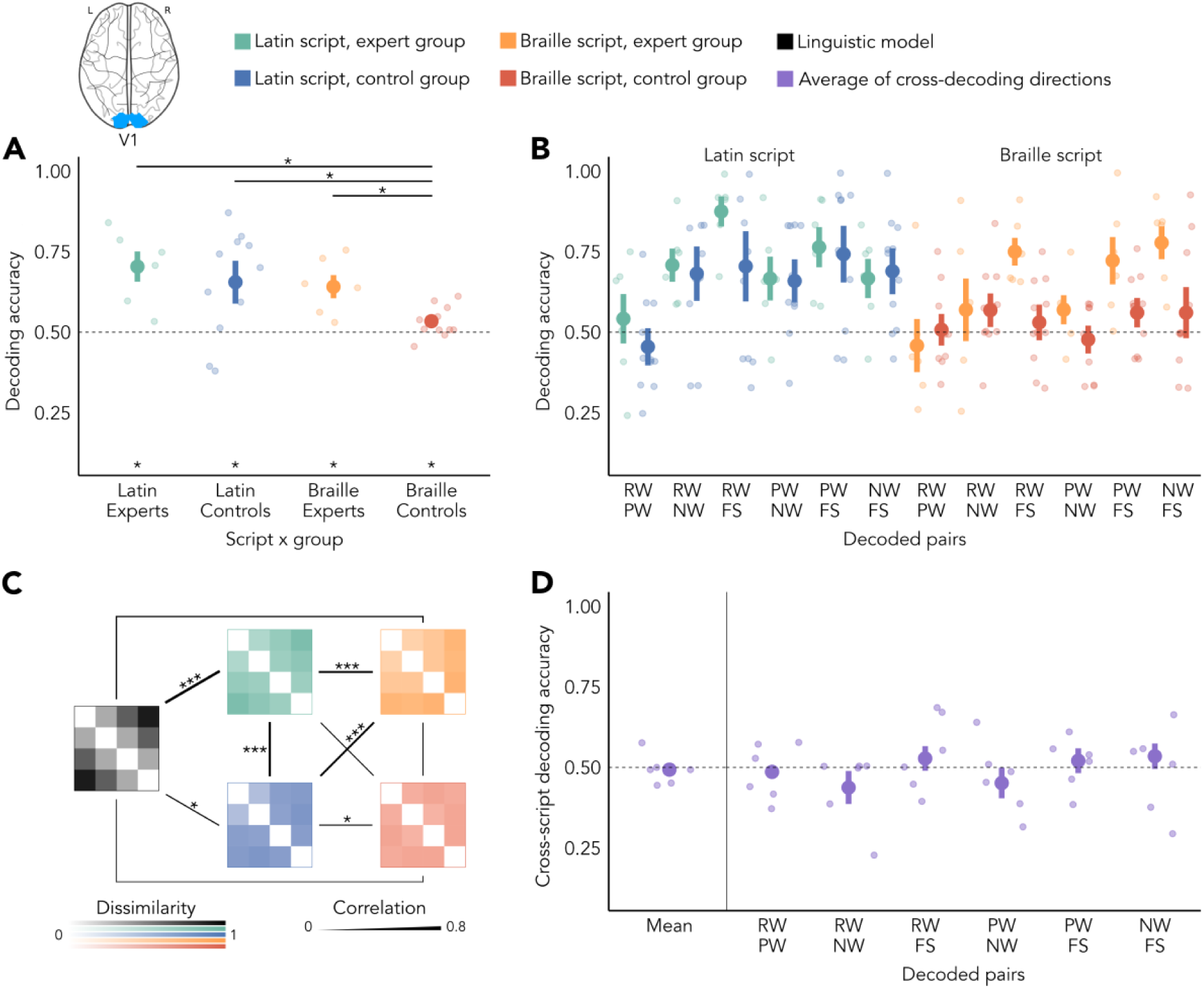
Multivariate representation of linguistic properties in V1. **(A)** Average of pairwise decoding of conditions that differ in linguistic properties. Stars above the labels on the x-axis represent the statistical significance of the decoding accuracy, while stars above the horizontal bars on top of the graph represent the significance of the difference between conditions (* p ≤ 0.05, ** p ≤ 0.01, *** p ≤ 0.001). Error bars represent standard error. In all the script-group pairs, we found significantly above-chance decoding and significant differences with the unknown and non-decodable condition of Braille presented to a control group. **(B)** Pairwise decoding within scripts. While experts and controls showed no group difference for their common native script, they showed a group difference for Braille, together with a main effect of condition and interaction between the variables, with experts showing a pattern similar to the one of Latin script. Details of post-hoc analyses can be found in Figure 5-2. **(C)** Representational similarity analysis (RSA) between neural representations (green, orange, blue, red) and with a linguistic model (black). Strength of the correlation is represented by the thickness of the bar connecting two patterns. In V1, patterns of known scripts correlate strongly with each other. Patterns relative to control participants showed a significant correlation across scripts. Only the neural RDMs relative to the organization of the Latin script show a correlation with the linguistic distance model, contrary to what happens in the case of Braille. **(D)** Cross-script decoding. The mean of cross-decoding across all pairs of linguistic conditions did not show significant above-chance decoding, and neither did the individual pairs. RW = Real Words; PW = Pseudowords; NW = Nonwords; FS = Fake Script. Region of interest comes from an example subject. The full list of statistical test can be found in Figure 5-1.

### Script-independent representation of linguistic properties in the Left Posterior Temporal Region

Lastly, we observed reliable activations outside of our ROIs when we contrasted intact Latin words over scrambled ones. By masking these activations to regions of the language network (Fedorenko et al., 2010), we identified the left-posterior temporal region, as a higher-order linguistic area. We limited our analyses to those subjects who show univariate activation for Latin words (fifteen participants, five experts). Paired t-test on the mean activation for intact Braille and scrambled dots did not highlight overall preferential activations by intact Braille in the expert group as a whole (t(4) = 2.05, p = 0.11), nor in the control group, as expected (t_(10)_ = -1.24, p = 0.24). These results did not change when eye movements were considered in the GLM. Multivariate analyses within each script revealed an overall profile very similar to the one previously seen in other ROIs. Pairwise comparisons (Figure 6A) could be significantly decoded if the script is known by the participants (all p_FDR_ < 0.001), while the script that is unknown to the group had chance-level accuracy (DA = 0.52, t_(9)_ = 0.925, p_FDR_ = 0.24), and significant differences emerged between scripts known and not known by the group of participants (all p_FDR_ < 0.001). More in-depth, pairs of comparison (Figure 6B) showed large differences within Braille (group effect: F_(1)_ = 58.416, p < 0.001; condition effect: F_(5)_ = 4.229, p = 0.002), while in the Latin script we observed only an effect of conditions (F_(5)_ = 4.991, p = 0.001) and no group differences (F_(1)_ = 0.833, p = 0.38). Post-hoc analyses on the differences between the conditions show that shared linguistic properties are harder to decode (RW-PW and PW-NW, p_Bonf_ = 0.38) than more linguistically distant comparisons (p_Bonf_ < 0.017). The similarity structure of pairwise comparisons (Figure 6C) showed significant correlations between known scripts (all p_FDR_ < 0.001) and no correlation with unknown scripts. Intriguingly, no neural RDM correlated with our linguistic model (all p_FDR_ > 0.06). The lack of correlation to the theoretical model suggests different coding principles compared to the other ROIs. Even more striking, and in contrast to all other ROIs, we found the average pairwise cross-decoding accuracy (Figure 6D) to be significantly higher than chance (DA = 0.66, t_(5)_ = 4.08, p_FDR_ = 0.02, D = 1.8). More in detail, several of the cross-scripts binary decoding were significant (RW-NW, : DA = 0.78, t_(5)_ = 4.49, p_FDR_ = 0.02; PW-NW: DA = 0.68, t_(5)_ = 3.64, p_FDR_ = 0.025; PW-FS: DA = 0.68, t_(5)_ = 2.69, p_FDR_ = 0.038; NW-FS: DA = 0.75, t_(5)_ = 3.35, p_FDR_ = 0.025). Real Words compared to the Fake Script did not seem to generalize (DA = 0.52, t_(5)_ = 0.293, p_FDR_ = 0.39). A borderline-significant trend emerged instead in the generalization of the comparison between Real Words and Pseudo-Words (DA = 0.58, t_(5)_ = 2.36, p_FDR_ = 0.052, D = 1). Together with all this evidence for cross-script decoding, we also observed significant differences with the average within-script decoding accuracy (t_(5)_ = 3.23, p = 0.031). These findings reveal, overall, a partially script-independent representation of linguistic content. Detailed statistical tests and results are available in Figure 6-1.

**Figure 6.**
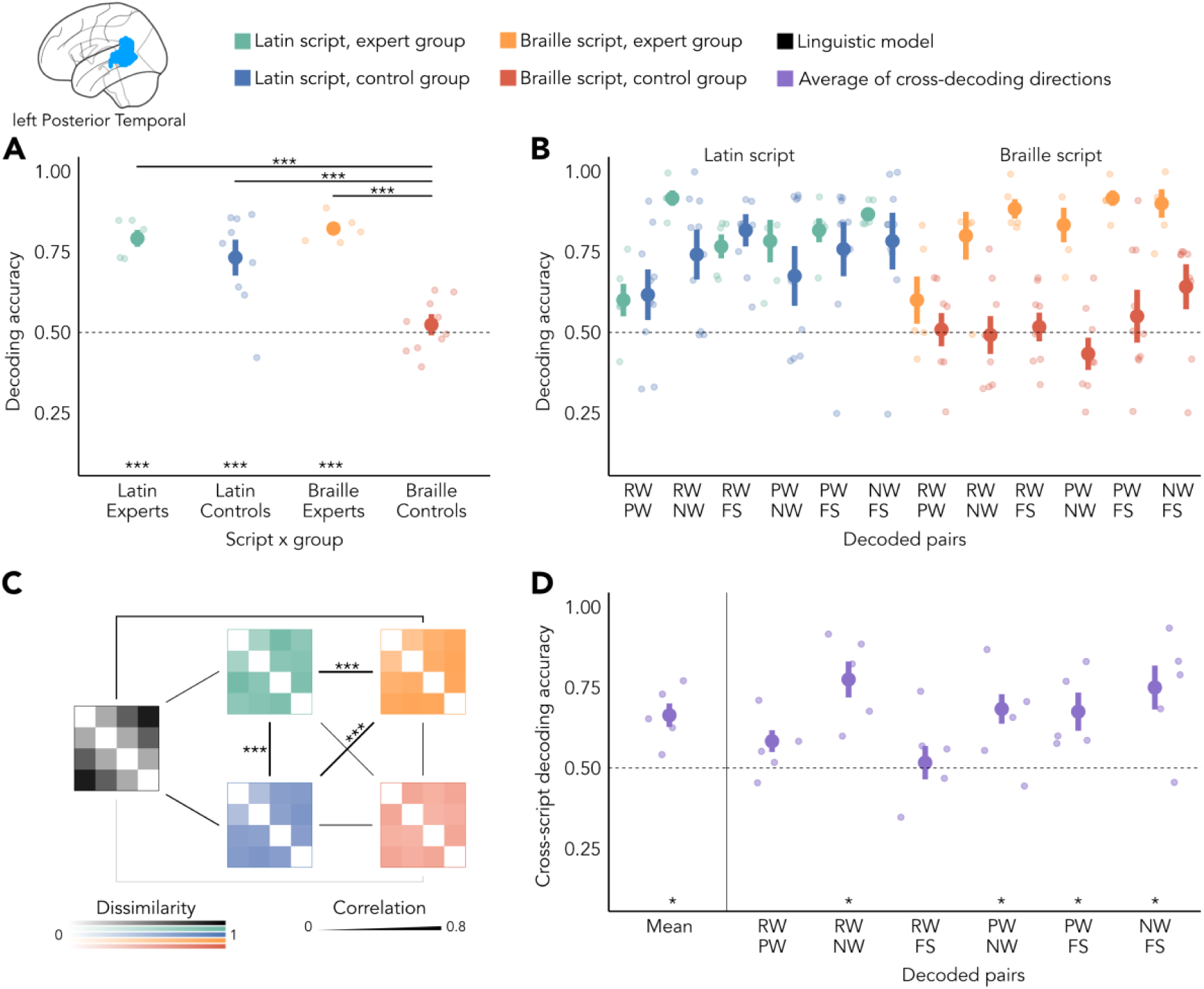
Multivariate representation of linguistic properties in l-PosTemp. **(A)** Average of pairwise decoding of conditions that differ in linguistic properties. In all the known scripts (Latin and Braille in experts, Latin in controls), we found significantly above-chance decoding and strong significant differences with the unknown and non-decodable condition of Braille presented to a control group. **(B)** Pairwise decoding within scripts. While experts and controls showed no group difference for their common native script, they showed a group difference for Braille, with experts showing a pattern similar to the one of Latin script. Details of post-hoc analyses can be found in Figure 6-2. **(C)** Representational similarity analysis (RSA) between neural representations (green, orange, blue, red) and with a linguistic model (black). Strength of correlation is represented by the thickness of the bar connecting two patterns. Known scripts correlated highly with each other and showed no correlation with the Braille-control pair. No correlation between neural RDMs and the model emerges. **(D)** Cross-script decoding. The mean of cross-decoding across all pairs of linguistic conditions showed significant above-chance decoding, hinting at generalization effects. Different individual pairs can be decoded from one script to another, while RW-PW shows an almost-significant trend (p = 0.052). In all the figures, stars represent statistical significance (* p ≤ 0.05, ** p ≤ 0.01, *** p ≤ 0.001). Error bars represent standard error. Region of interest comes from an example subject. The full list of statistical test can be found in Figure 6-1.

## Discussion

We investigated the neural bases of processing a peculiar script, visual Braille, that differs from most common scripts for its lack of shape features (Changizi et al., 2006). We found in expert readers a preference for words written in Braille over scrambled arrangements of dots in the visual word form area, as well as multivariate neural representations of linguistic content similar to the one of the participants’ native script. Despite these similarities between scripts in the nature of the representations, we did not find generalization of the neural representation between scripts. We also found similar multivariate processing of written text beyond VWFA, in the early stages of visual processing (V1), in shape-sensitive areas (LO), and in higher-order linguistic regions (l-PosTemp). l-PosTemp is the only region showing successful generalization of the two scripts and thus a partially script-invariant representation of linguistic properties. In the visual areas, conversely, At the univariate level, we found that in expert visual Braille readers, the same site in the ventral occipito-temporal cortex that selectively responds to words presented in the participant’s native Latin script also exhibits a preference for Braille words over scrambled dot arrangements. This responsiveness was absent in the control group, suggesting that it can be attributed to neural adaptations in the experts’ brains resulting from their training and experience in reading visual Braille. Previous studies showed increased selectivity in VWFA emerging with the acquisition of reading abilities (Dehaene et al., 2015; Dehaene-Lambertz et al., 2018). This sensitivity was investigated in the acquisition of specific scripts (e.g. Hebrew; Baker et al., 2007) and scripts made out of houses and faces (L. Martin et al., 2019; Moore et al., 2014), suggesting that VWFA could globally link symbols with their represented linguistic properties acquired through learning. Other studies emphasized the importance of shape features for the location and activation of VWFA (Dehaene et al., 2005). However, all investigations were aimed at scripts that are composed of canonical line-junctions, and this property of the stimuli may explain why they are co-opted by the reading network. Therefore, none of these studies elucidated whether shape (global form extracted from joined lines) is a mandatory feature to activate VWFA (Ben-Shachar et al., 2007; Roberts et al., 2013; Vogel et al., 2012; Xue & Poldrack, 2007). Other studies investigated Braille, as a unique orthographic system not composed of such explicit shape cues. Previous neuroimaging studies tested tactile Braille, in blind (Büchel et al., 1998; Reich et al., 2011) or sighted (Siuda-Krzywicka et al., 2016) individuals. In blind people, it cannot be excluded that the recruitment of typical orthographic regions can be due to neuroplasticity that occurs in the absence of vision (Xu et al., 2023). In the sighted, activations found for visual Braille could be attributed to mental imagery of classical orthographic letters mapped from tactile Braille (Zhang et al., 2004), a phenomenon arguably less likely when a visual stimulus is presented to the participant. Nevertheless, as a whole these studies already suggested that the domain-specific function of stimuli to read, be it regular scripts, specially trained houses/faces, or even nonvisual scripts, results in a selective response in VWFA for these stimuli. Our findings with visual Braille are consistent with this role of domain-specific task-related factors, and extend them in the direction of a very simple script that lacks the explicit shape cues that are often invoked to explain the location of the VWFA in the visual system.

The similarities between scripts go beyond univariate selectivity and also include the coding principles used to represent the scripts as investigated through multivariate analyses. Differences between patterns of different classes of linguistic stimuli (here Words, Pseudo-Words, Non Words and Fake Scripts) emerged based on the expertise of a script rather than its visual features. In other words, when the visual Braille is learned, it is represented similarly to the native, line-based alphabet. When we consider the patterns of brain activity elicited by our various linguistic conditions, a clear and graded multivoxel selectivity appears evident for visual Braille for conditions with a different lexical status and orthographic properties: Real Words, Pseudo-Words, Non-Words (consonant strings), and scrambled dot patterns. This selectivity was virtually absent in naive participants reading Braille. Furthermore, the coding principles for Braille appeared strikingly similar and strongly correlated to those relative to the Latin-based script.

However, the emerging selectivity for visual Braille is not all about domain specificity and linguistic properties. Our study provided a unique opportunity to investigate how this role of domain specificity interacts with the coding of visual features. Previous studies that showed activity in VWFA for special scripts did not investigate the multivariate organization of these scripts: whether they share common neural representations (suggesting amodal or cross-modal representations) or they co-exist in segregated spaces within the same regions. In our study, we were able to directly compare, in the same participant, orthographic systems with different visual characteristics but common linguistic content. Whereas similar coding principles characterize visual Braille and Latin alphabet processing, the significant drop in cross-script decoding relative to within-script decoding combined with the absence of cross-scripts generalization (present instead in l-PosTemp), demonstrates that the representations of linguistic properties in VWFA are interdigitated across both scripts. Different neural populations implement the two scripts. These findings resonate with the results of a recent 7T imaging study on French/Chinese bilinguals. In bilinguals, high similarity was present in activated neural populations between French and English, but less so between French and Chinese (Zhan et al., 2023). However, bilinguals might activate different sounds with the two scripts, and the dissociation between French and Chinese could be due to both orthography and phonology. Our study disentangles visual form from phonology and other linguistic factors. While the Latin-based script and visual Braille are visually highly dissimilar, they both map onto the same French words and the same sounds. The total absence of cross-decoding and its significant drop relative to within-script decoding provide clear evidence for the importance of visual form in determining which neural populations in VWFA are selectively activated. The absence of cross-decoding also constraints the interpretation of findings within each script, more than any of the previous studies investigating similar matters (most notably, Vinckier et al., 2007). Within each script there is coding of properties such as lexical access (Words vs other conditions) and statistical regularities (e.g., Pseudo-Words vs Non Words), but in a script-specific coding rather than an abstract linguistic code.

Given that expertise in visual Braille is rare and thus we anticipated a relatively low number of experts, we performed pilot experiments to select two powerful experimental designs that provide reliable data despite these lower numbers (e.g. very consistent pattern across conditions in the two groups for Latin script in Figs. 3B, 4A, 4B, and 5B, comparisons that can be considered internal replications), and that are able to provide clear data even at an individual level, including univariate activity (see Fig. 2) and high multivariate decoding accuracies (e.g., very clear split between experts and controls on Braille in Fig. 3A). As a result, the expertise-related effects are striking. However, the null result of finding no cross-decoding could depend on the number of participants. It is possible that a study with a much larger group of experts would in the end find a cross-decoding that is still small but significant. Here it is important that the conclusions about independent, script-specific representations are also fully supported by the significant drop in cross-decoding relative to within-script decoding. This is a clear effect and not a null result. Furthermore, the findings in l-PosTemp reveal that it is possible to find significant cross-decoding with our design and analyses methods. Together, our findings indicate that script specificity is a major property of the expertise-related representations in VWFA.

Similarly, the choice of an implicit task reveals the strength of the results. To be able to accurately compare the responses of the two groups, we opted for a task (1-back, see Methods) that could be performed by both groups given with the same instruction. It is likely that naive Braille readers engaged in a purely visual task when facing Braille stimuli while the expert participants would additionally employ a linguistic strategy. The presentation of Latin and Braille alphabets in different run, reduces the contextual effects between them.

Other visual areas showed no univariate differences between Braille and Scrambled Braille bilaterally in V1 and shape-selective LO in expert Braille readers. However, in expert readers, these regions showed similar multivariate profiles to the one identified in VWFA. This similarity in the patterns of linguistic distance between both known scripts reveals distributed expertise-induced plasticity in the visual system. Studies of visual word forms focused mainly on VWFA, but more widespread linguistic selectivity have already been observed for shape-based scripts across the occipito-temporal cortices (Szwed et al., 2014; Siuda-Krzywicka et al., 2016). Although V1 has already been associated with plasticity effects beyond VWFA and preference for known scripts (e.g. Latin-based and Chinese; Szwed et al., 2014), it is remarkable that such adaptation extends to a script that lacks explicit shape cues. Similarly, it is striking that LO shows significant coding of word-like stimuli, and that such coding is found in both hemispheres. Previous studies on the gradient of involvement in word processing (Vinckier et al., 2007) did not perform such multivariate analyses, so possibly the literature underestimates the involvement of LO in word processing. Indeed the multivariate profile in LO is not a simple consequence of visual differences across classes of stimuli, as this representation was not observed in control participants for Braille; nor is it restricted to typical script shape processing, as we also observed this coding in visual Braille for experts. This is particularly striking because LO is often seen as the major region supporting shape perception (Grill-Spector et al., 2001; Kanwisher et al., 1996; Malach et al., 1995), and visual Braille letters contain much less shape cues compared to other visual scripts. We note however that the absence of explicit shape cues does not exclude the possibility that the visual system would still engage in forms of pattern recognition, like Gestalt principles, such as proximity and good continuation, that would include the perception of subjective line and shape elements that would connect the dots with lines and more complex geometric patterns. However these principles might also emerge in the non-expert group, therefore not explaining our results. Also, previous studies have suggested that the Braille script is more difficult to learn than line or shape-based scripts, also suggesting the idiosyncrasy of Braille compared to line and shaped based scripts (Bola et al., 2017).

Beyond visual regions, we reliably identified activated voxels in the left posterior temporal area (l-PosTemp) with patterns of multivariate activity that resemble the ones found in visual areas. Although the neural patterns for known scripts presented strong correlations with each other, we surprisingly found no correlation with the linguistic distance model. The absence of correlations may be attributed to the model, inspired by results shown in vOTC (Vinckier et al., 2007), a region highly sensible to visual input. It is possible that the role of posterior temporal regions in phonological processing (Bhaya-Grossman & Chang, 2022; Chang et al., 2010) and reading (A. Martin et al., 2015) determines non-linear relationships between linguistic properties. The preferential phonological tuning of this region could lead to increased similarity between Words and Pseudo-Words, and increased dissimilarity to low-frequency phonological categories (Non-Words and Fake Script). Finally, the cross-script generalization indicates that the representations of linguistic conditions are invariant to the visual differences. Such common, script-invariant representation further demonstrates that at this stage of information processing in l-PosTemp, linguistic information is abstracted from the visual properties of the stimuli processed in the ventral occipito-temporal cortex (A. Martin et al., 2015). Note though that this script invariance was only partial, because the cross-script decoding was significantly smaller than the within-script decoding.

Overall, we observe that several regions of the reading network are involved in the processing of the orthographic material in a script without explicit line junctions. Looking beyond the domain of reading, our work adds to the literature showing effects of learning in visual processing, including learning the statistical properties of the stimuli such as manipulated in the comparison of Pseudo-Words and Non Words. Such learning has been found in other domains, namely the recognition of chessboards in professional chess players (Bilalic et al., 2011) or the processing of Greeble stimuli (Gauthier et al., 1998; Gauthier & Tarr, 1997) in the fusiform face area (FFA), proving the ability of VOTC to adapt to an extended set of stimuli. We show that a coding of the statistical regularities that compose the stimuli extends beyond the canonical line-based script and also apply to visual Braille, a script that is void of those explicit shape cues. Such impact does not come as a surprise, as statistical learning is part of the language acquisition process, but it is remarkable that it extends to this broad variety of visual features.

In conclusion, despite marked differences in the visual properties of Braille and Latin-based scripts, the coding principles of occipital regions showing a preference for a Latin-based script are strikingly similar to responses to visual Braille in experts readers. In occipital regions the neuronal populations that implement these principles do not overlap, suggesting interdigitated representation across scripts co-existing in the same region (Zhan et al., 2023); while in posterior temporal regions scripts are represented in a more script-invariant manner.

## Data and code availability

All the code used to perform the experiments (https://github.com/fcerpe/VBE_experiment) and analyse the results (https://github.com/fcerpe/VBE_data) is publicly available on GitHub. Raw defaced MRI scans are available on GIN in BIDS (Gorgolewski et al., 2016) format (https://gin.g-node.org/cpp_brewery/2022_StLuc_VisBraExpertise_FC_raw).

## Author contributions

**Figure 7.**
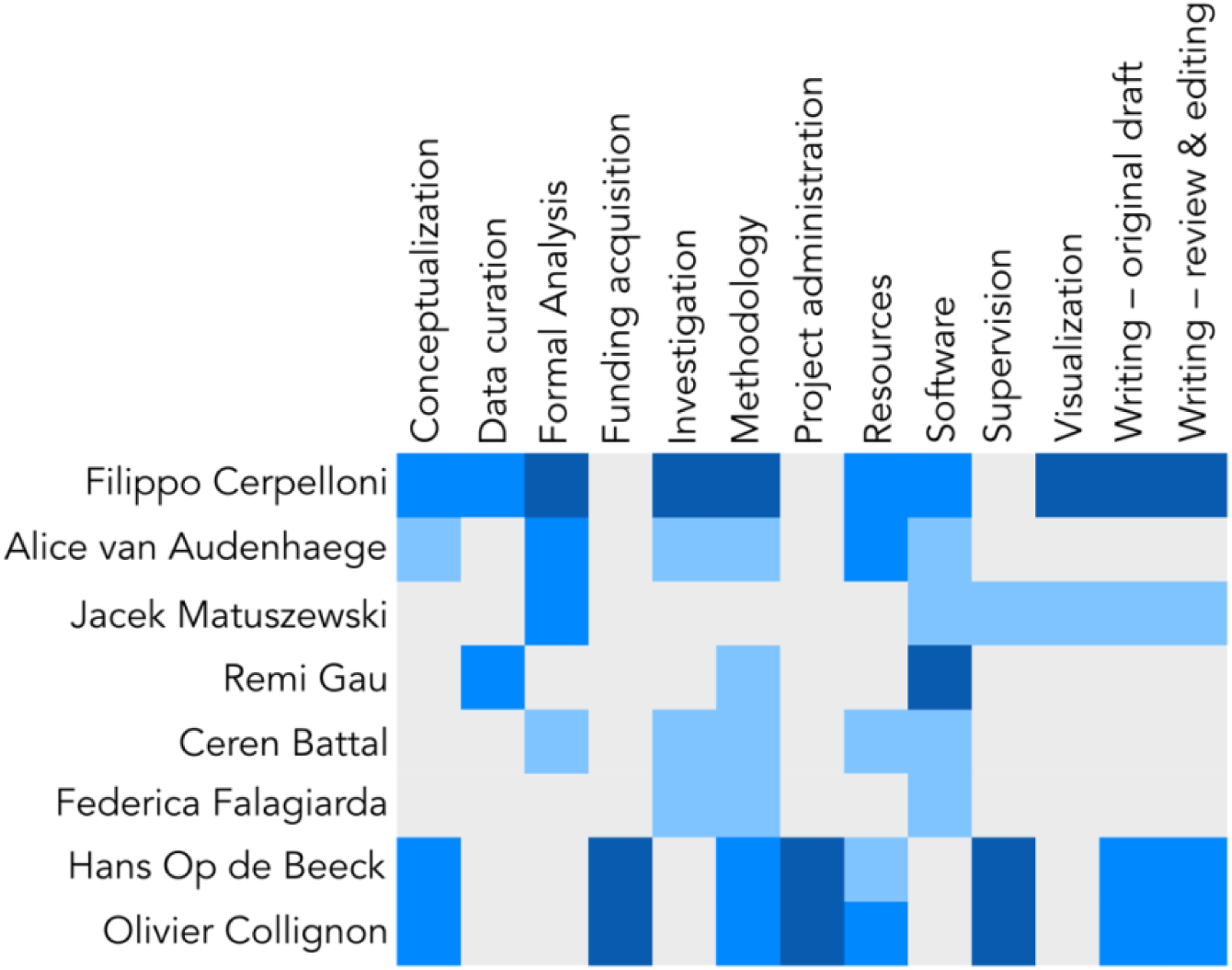
Author contributions. Based on the CRedIT (https://credit.niso.org/) taxonomy. Each contribution was assessed as ‘lead’ (dark blue), ‘equal’ (mid blue), ‘support’ (light blue).

## Acknowledgements

The authors would like to thank the Institut Royal pour Sourds et Aveugles (IRSA) and the association EQLA for their help in recruiting the Braille experts. The project was funded in parts by a Mandat d’Impulsion Scientifique awarded to OC, the Belgian Excellence of Science (EOS) program (Project No. 30991544) awarded to HO and OC, a Flagship ERA-NET grant SoundSight (FRS-FNRS PINT-MULTI R.8008.19) awarded to OC, and research project G0D3322N of the Fund for Scientific Research (FWO) Flanders and Methusalem project METH/24/003 awarded to HO. AV is a research fellow, CB and JM are postdoctoral researchers, and OC is a senior research associate at the Fond National de la Recherche Scientifique de Belgique (FRS-FNRS). FC received funding from a KU Leuven-UCLouvain cooperation grant.

## Appendix

**Table 2-1.** Functional activation of Visual Word Form Area (VWFA) in expert visual Braille readers. For each expert, we contrasted intact Latin words (FW) to scrambled ones (SFW) and obtained activation peaks. To determine the cluster size of VWFA, we performed Small Volume Correction in a 10mm radius sphere around the individual peak coordinates of VWFA identified through a Latin-script contrast (FW > SFW). Next, we performed an additional Small Volume Correction around the same peaks to determine significantly activated voxels for the contrast of Braille words (BW) over scrambled dots (SBW). MNI coordinates refer to the peak of each contrast.

| Contrast | Subject | p-value (FWE corrected) | z-score | Cluster size | MNI coordinates |
| --- | --- | --- | --- | --- | --- |
| FW > SFW | exp-01 | 0.000 | 5.04 | 24 | -47 -60 -13 |
|  | exp-02 | 0.000 | 4.78 | 9 | -52 -47 -18 |
|  | exp-03 | 0.000 | 6.57 | 83 | -52 -55 -18 |
|  | exp-04 | 0.000 | 5.90 | 159 | -44 -63 -13 |
|  | exp-05 | 0.002 | 4.31 | 29 | -44 -60 -15 |
|  | exp-06 | 0.000 | 5.27 | 27 | -44 -60 -13 |
| BW > SBW | exp-01 | 0.000 | 5.09 | 113 | -49 -63 -13 |
|  | exp-02 | 0.000 | 5.23 | 92 | -55 -47 -21 |
|  | exp-03 | NA | NA | NA | NA |
|  | exp-04 | 0.034 | 3.50 | 7 | -39 -60 -5 |
|  | exp-05 | 0.003 | 4.23 | 37 | -42 -57 -8 |
|  | exp-06 | 0.000 | 5.39 | 79 | -47 -63 -15 |

**Figure 2-2.**
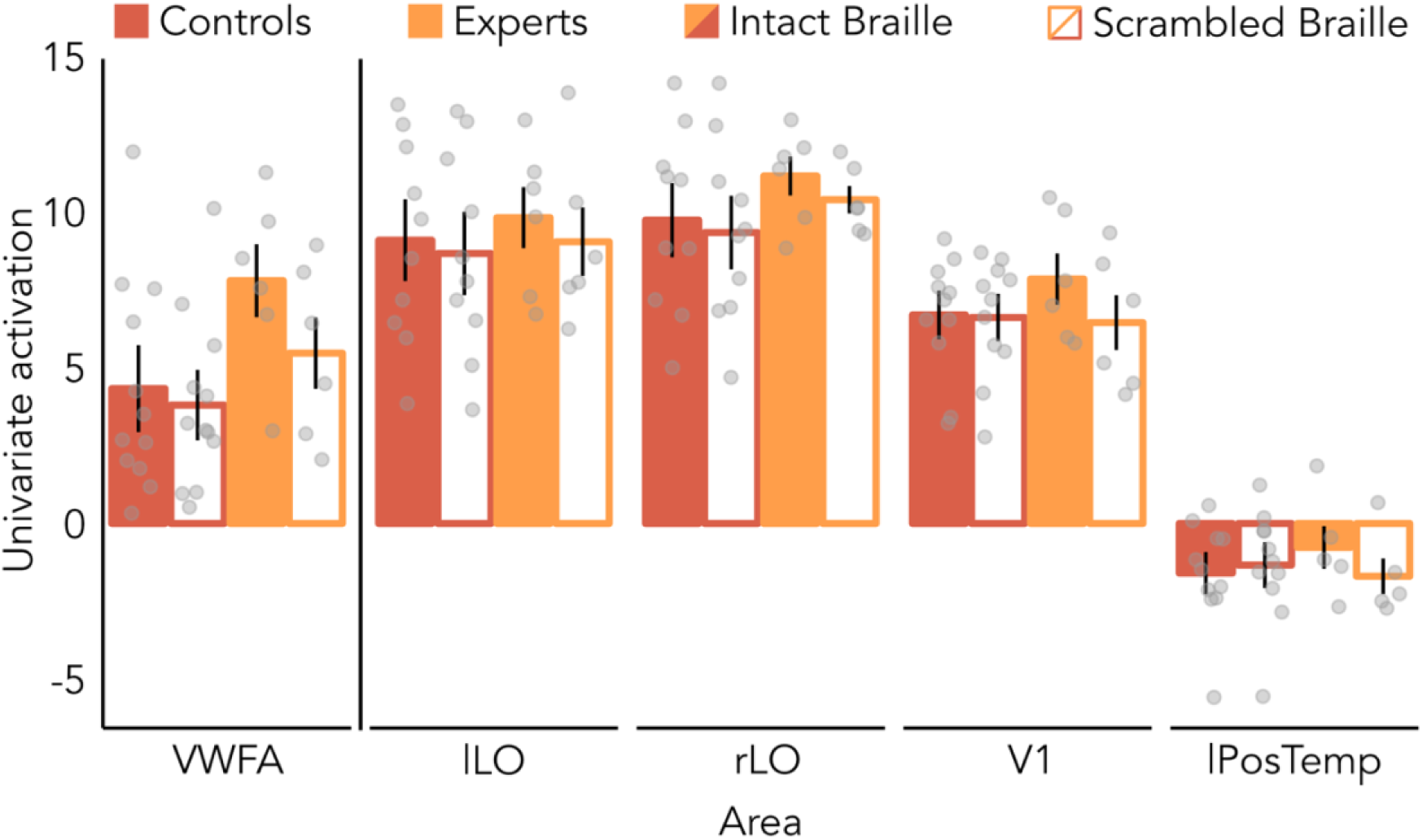
Univariate selectivity for intact over scrambled Braille across ROIs. For each ROI, we extracted the mean activation individually for intact Braille (full bars) and scrambled Braille (empty bars) stimuli. Black bars represent standard error (SE), grey dots represent individual subjects.

**Figure 2-3.**
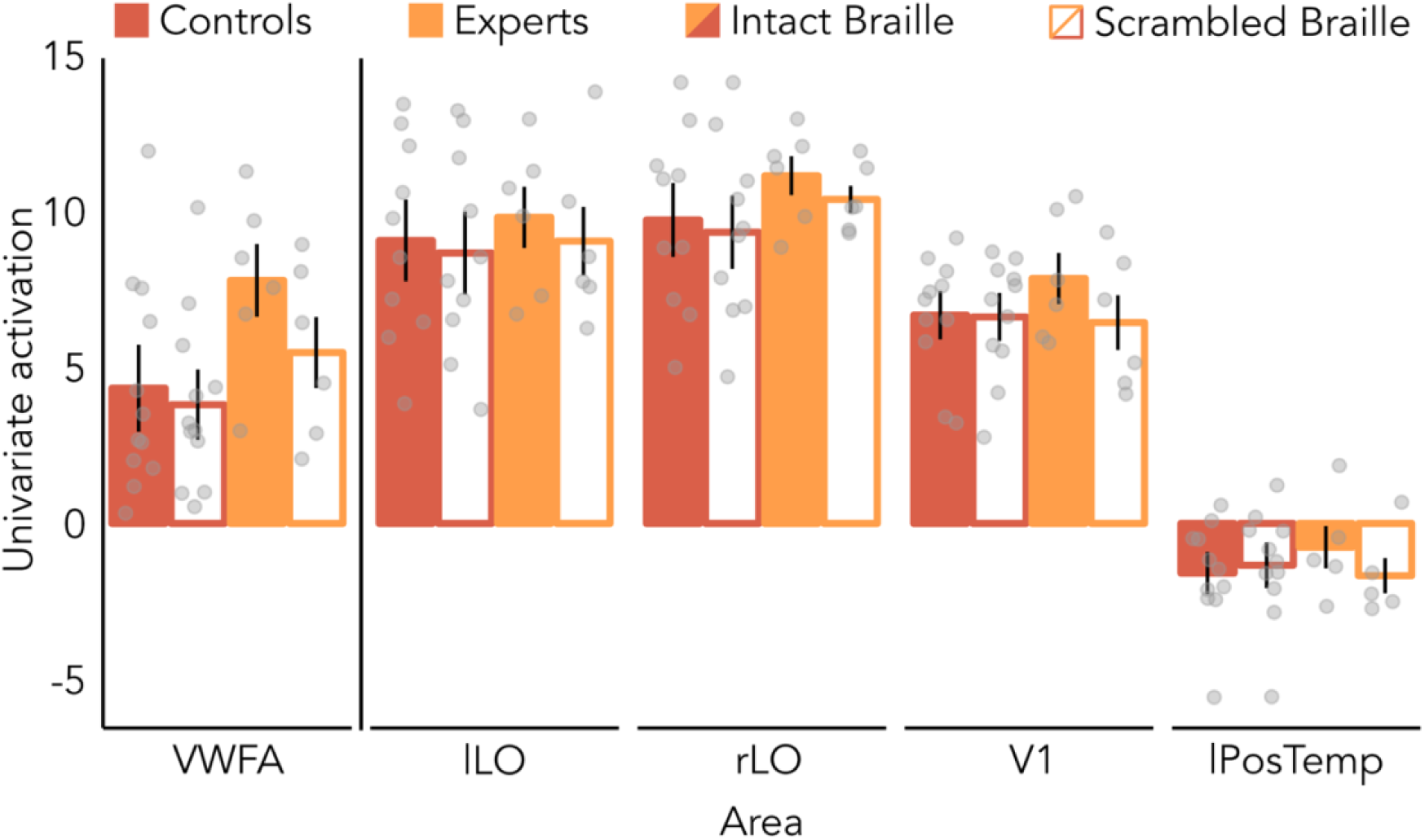
Univariate selectivity for intact over scrambled Braille across ROIs with eye movements. For each ROI, we extracted the mean activation individually for intact Braille (full bars) and scrambled Braille (empty bars) stimuli, using the GLM analysis where eye movements (computed through bidsMReye toolbox). Black bars represent standard error (SE), grey dots represent individual subjects.

**Figure 3-1.**
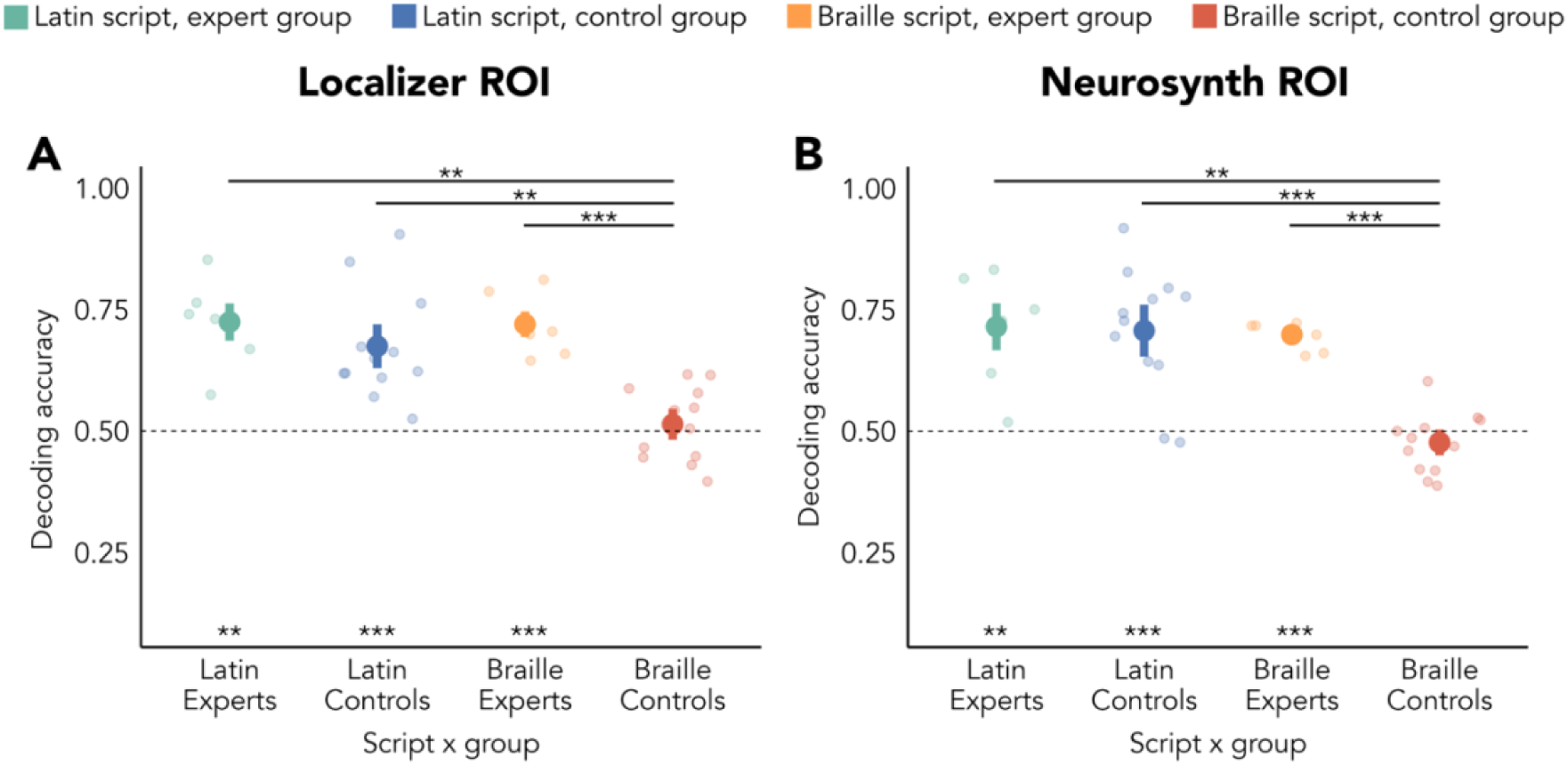
Comparison of ROI definitions and their multivariate representation of linguistic properties in the Visual Word Form Area (VWFA). (A) Region of Interest defined through the methods reported in “Definition of the Regions of Interest (ROIs)”. (B) Region of Interest defined through Neurosynth (keyword: “visual words”). We selected the highest-responding 100 voxels and applied the mask to each subject, using the same analysis pipeline used for our custom definition.

**Figure 3-2.**
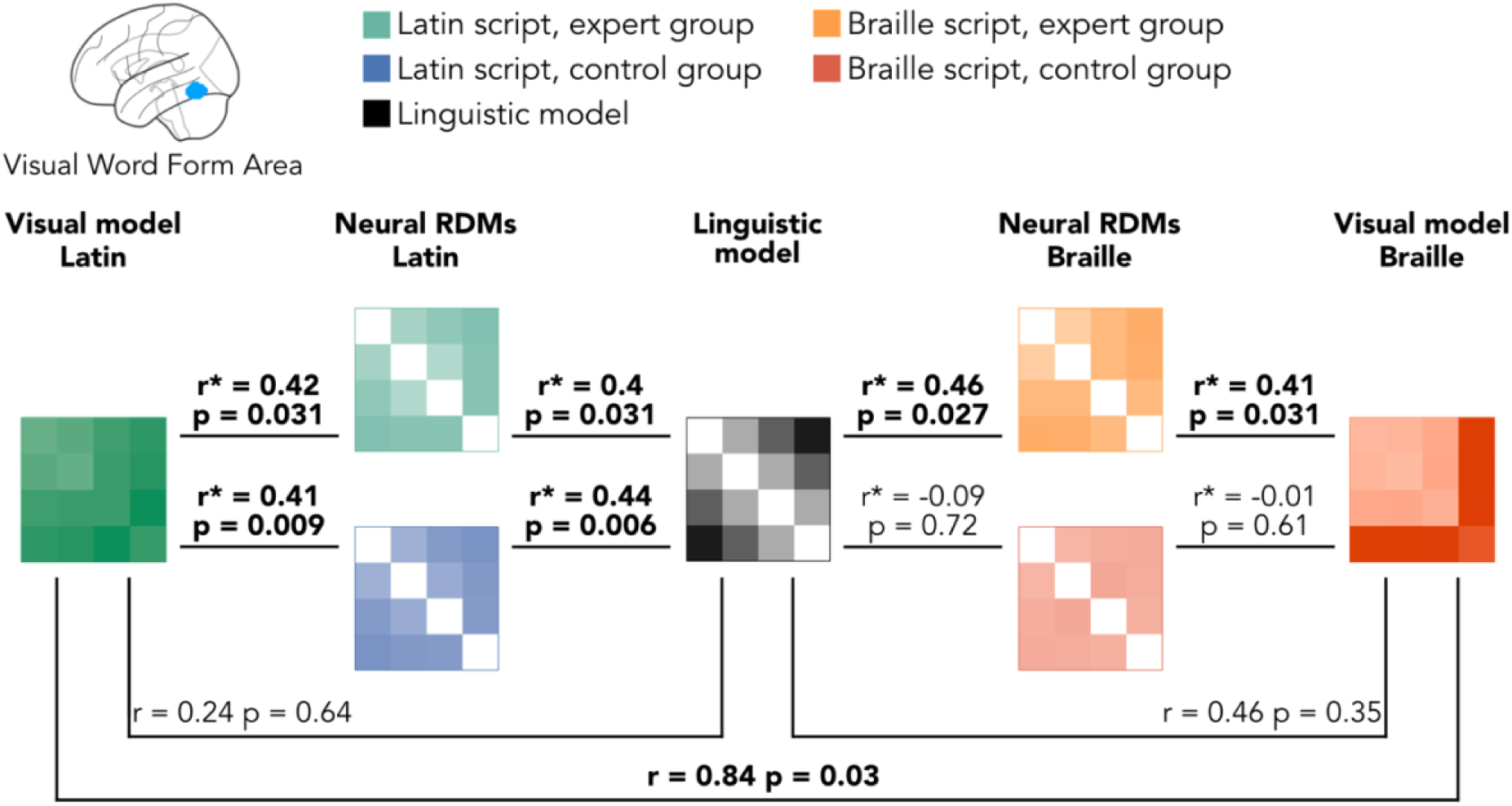
Correlations between neural and model RDMs in the Visual Word Form Ara (VWFA). Each correlation between the neural matrices and a model was made as a partial correlation, controlling for the other model. As an example, in the correlations between the neural RDM of the Latin script in one group and the linguistic model, we partialed out the correlation between the neural RDM and visual model for the Latin script. In the case of neural RDM and visual model correlations, we partialed out the correlation between the neural RDM and the linguistic model. Correlations between models, not partial, show a limited correlation between linguistic and visual explanations of the neural organization. Partial correlations are represented by r*. Significant correlations are represented in bold.

**Table 3-1.** Activation peaks for all the ROIs extracted from the functional localizer experiment. We localized VWFA individually in each expert, by contrasting the activation for intact Latin words over scrambled Latin words. Additionally, we localized activity for Braille by contrasting activation for intact Braille stimuli over scrambled ones. Bilateral Lateral Occipital (LO) was localized through the contrast of Line Drawing over scrambled Line Drawings. Details of the methods are available in the *Definition of the Regions of Interest (ROIs)* section

| Region | Group | Hemisphere | Contrast | Subject | Z-score | Cluster size | MNI coordinates |
| --- | --- | --- | --- | --- | --- | --- | --- |
| VWFA | Experts | Left | FW > SFW | Exp-01 | 5.04 | 24 | -47 -60 -13 |
|  |  |  |  | Exp-02 | 4.78 | 9 | -52 -47 -18 |
|  |  |  |  | Exp-03 | 6.57 | 83 | -52 -55 -18 |
|  |  |  |  | Exp-04 | 5.90 | 159 | -44 -63 -13 |
|  |  |  |  | Exp-05 | 4.31 | 29 | -44 -60 -15 |
|  |  |  |  | Exp-06 | 5.27 | 27 | -44 -60 -13 |
|  |  |  | BW > SBW | Exp-01 | 5.09 | 113 | -49 -63 -13 |
|  |  |  |  | Exp-02 | 5.23 | 92 | -55 -47 -21 |
|  |  |  |  | Exp-03 | NA | NA | NA |
|  |  |  |  | Exp-04 | 3.50 | 7 | -39 -60 -5 |
|  |  |  |  | Exp-05 | 4.23 | 37 | -42 -57 -8 |
|  |  |  |  | Exp-06 | 5.39 | 79 | -47 -63 -15 |
|  | Controls | Left | FW > SFW | Ctr-01 | Inf | 536 | -44 -66 -13 |
|  |  |  |  | Ctr-02 | 4.38 | 24 | -49 -60 -10 |
|  |  |  |  | Ctr-03 | 6.06 | 203 | -47 -55 -8 |
|  |  |  |  | Ctr-04 | Inf | 212 | -47 -47 -20 |
|  |  |  |  | Ctr-05 | 6.80 | 413 | -44 -57 -20 |
|  |  |  |  | Ctr-06 | 6.16 | 357 | -47 -50 -23 |
|  |  |  |  | Ctr-07 | 3.71 | 4 | -48 -65 -18 |
|  |  |  |  | Ctr-08 | 6.82 | 94 | -47 -50 -23 |
|  |  |  |  | Ctr-09 | 3.51 | 24 | -42 -52 -10 |
|  |  |  |  | Ctr-10 | Inf | 849 | -52 -57 -15 |
|  |  |  |  | Ctr-11 | Inf | 510 | -55 -57 -15 |
|  |  |  |  | Ctr-12 | 5.33 | 226 | -55 -50 -13 |
|  | Controls | Left | BW > SBW | Ctr-01 | NA | NA | NA |
|  |  |  |  | Ctr-02 | NA | NA | NA |
|  |  |  |  | Ctr-03 | NA | NA | NA |
|  |  |  |  | Ctr-04 | NA | NA | NA |
|  |  |  |  | Ctr-05 | 3.33 | 5 | -47 -52 -21 |
|  |  |  |  | Ctr-06 | NA | NA | NA |
|  |  |  |  | Ctr-07 | NA | NA | NA |
|  |  |  |  | Ctr-08 | NA | NA | NA |
|  |  |  |  | Ctr-09 | NA | NA | NA |
|  |  |  |  | Ctr-10 | NA | NA | NA |
|  |  |  |  | Ctr-11 | NA | NA | NA |
|  |  |  |  | Ctr-12 | NA | NA | NA |
| LOC | Experts | Left | LD > SLD | Exp-01 | 5.92 | 45 | -39 -76 -15 |
|  |  |  |  | Exp-02 | Inf | 864 | -44 -65 -13 |
|  |  |  |  | Exp-03 | 4.85 | 406 | -42 -78 -2 |
|  |  |  |  | Exp-04 | 6.44 | 911 | -49 -78 -3 |
|  |  |  |  | Exp-05 | 7.59 | 461 | -49 -76 -10 |
|  |  |  |  | Exp-06 | Inf | 1355 | -49 -78 -3 |
|  |  | Right | LD > SLD | Exp-01 | 5.68 | 100 | 39 -68 -10 |
|  |  |  |  | Exp-02 | Inf | 284 | 47 -78 0 |
|  |  |  |  | Exp-03 | 7.86 | 395 | 44 -78 -8 |
|  |  |  |  | Exp-04 | 6.26 | 335 | 37 -70 -13 |
|  |  |  |  | Exp-05 | 7.37 | 227 | 49 -73 -10 |
|  |  |  |  | Exp-06 | Inf | 669 | 49 -68 -13 |
|  | Controls | Left | LD > SLD | Ctr-01 | 7.67 | 943 | -49 -76 0 |
|  |  |  |  | Ctr-02 | Inf | 2176 | -39 -73 -10 |
|  |  |  |  | Ctr-03 | Inf | 857 | -47 -73 -5 |
|  |  |  |  | Ctr-04 | Inf | 717 | -41 -78 -5 |
|  |  |  |  | Ctr-05 | Inf | 1147 | -49 -78 -5 |
|  |  |  |  | Ctr-06 | Inf | 909 | -47 -65 -13 |
|  |  |  |  | Ctr-07 | Inf | 446 | -42 -75 -10 |
|  |  |  |  | Ctr-08 | NA | NA | NA |
|  |  |  |  | Ctr-09 | NA | NA | NA |
|  |  |  |  | Ctr-10 | Inf | 1043 | -49 -73 -10 |
|  |  |  |  | Ctr-11 | 6.89 | 200 | -52 -75 0 |
|  |  |  |  | Ctr-12 | Inf | 634 | -47 -73 -13 |
|  |  | Right | LD > SLD | Ctr-01 | 6.83 | 613 | 49 -65 -5 |
|  |  |  |  | Ctr-02 | Inf | 2391 | 44 -65 -10 |
|  |  |  |  | Ctr-03 | Inf | 1097 | 44 -70 -10 |
|  |  |  |  | Ctr-04 | 6.55 | 81 | 44 -73 -7 |
|  |  |  |  | Ctr-05 | Inf | 327 | 44 -65 -13 |
|  |  |  |  | Ctr-06 | 6.73 | 539 | 49 -68 -7 |
|  |  |  |  | Ctr-07 | 4.90 | 36 | 49 -68 -10 |
|  |  |  |  | Ctr-08 | 5.20 | 71 | 39 -68 -5 |
|  |  |  |  | Ctr-09 | NA | NA | NA |
|  |  |  |  | Ctr-10 | 5.58 | 347 | 44 -68 -13 |
|  |  |  |  | Ctr-11 | 6.96 | 68 | 39 -76 -13 |
|  |  |  |  | Ctr-12 | Inf | 974 | 39 -73 -10 |

**Table 3-2.**
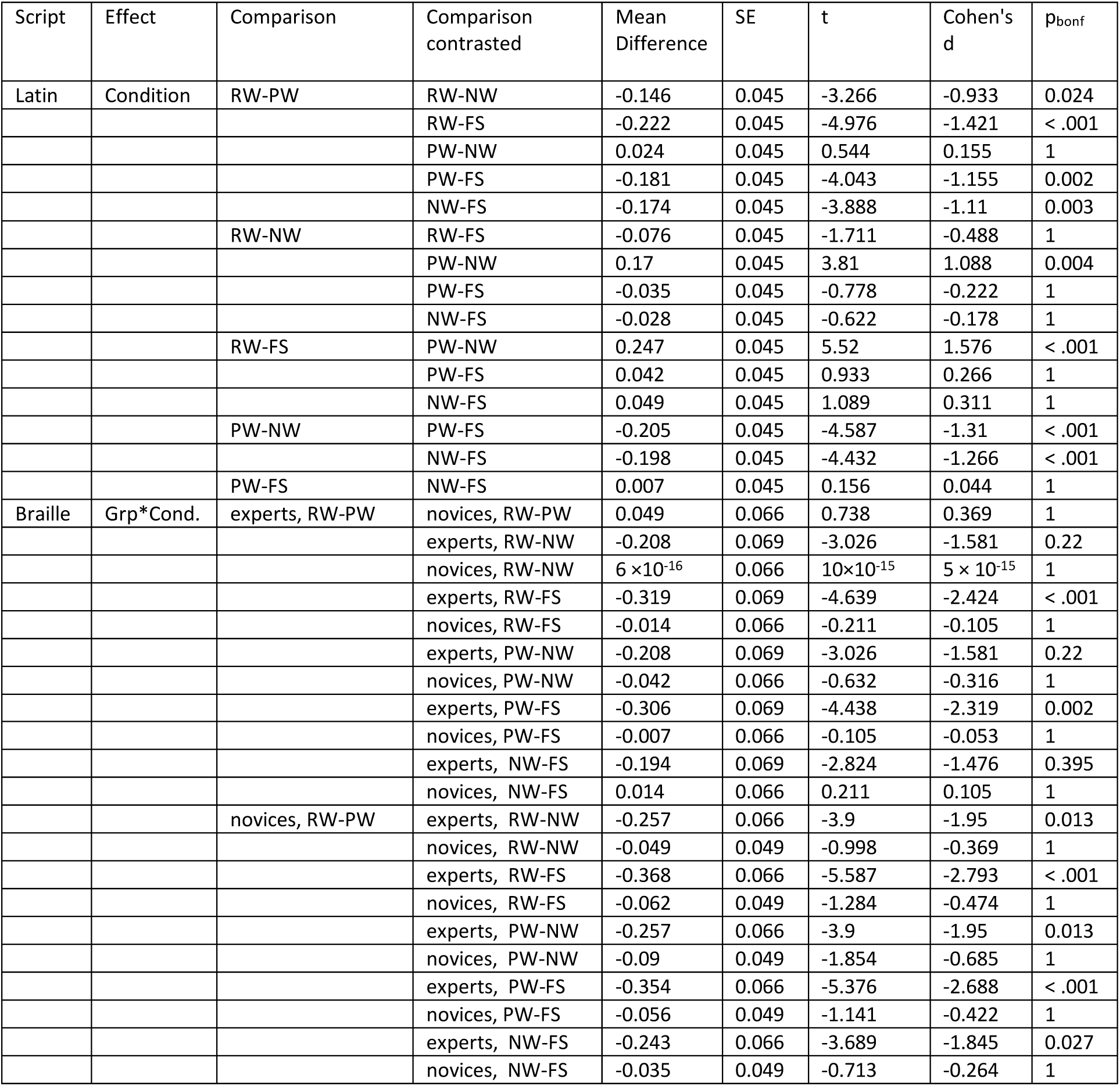
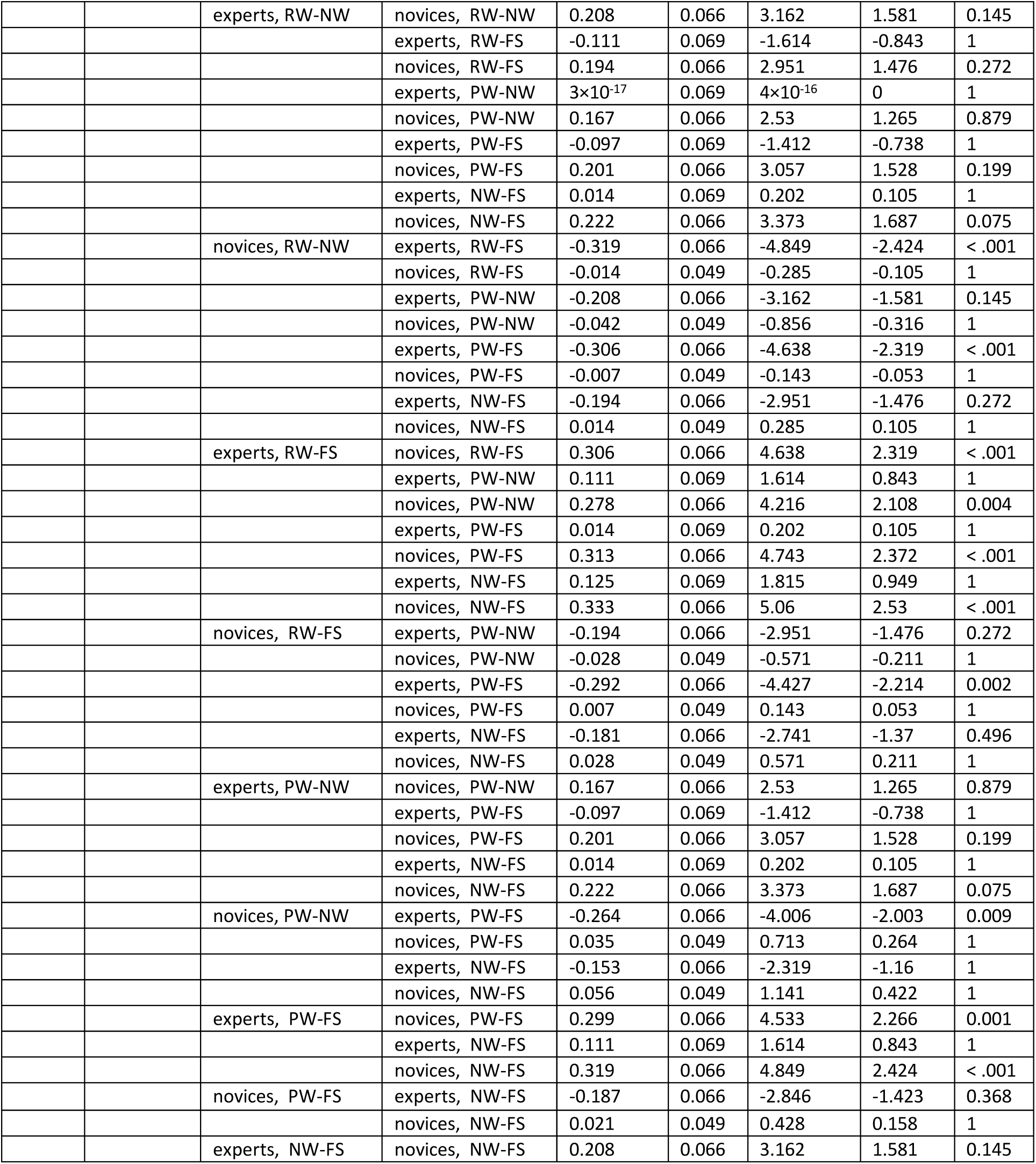
Post-hoc t-tests in Visual Word Form Area (VWFA). For each effect in the repeated measures ANOVA (rmANOVA), we contrasted the different comparisons to identify the causes of the interaction effects. In Latin script, results are averaged over the levels of ‘group’. P-values are adjusted for comparing a family of 15. In Braille script, p-values are adjusted for comparing a family of 66.

**Table 4-1.** Statistical tests in left-LO. Full list of statistical tests performed over the multivariate results of l-PosTemp. Each analysis is detailed with the conditions tested and the statistical test. Statistical tests detail the result, the degrees of freedom (DF), the p-value, and a measure of the effect size. For t-test, effect size was measured through Cohen’s D. For repeated measures ANOVA (rmANOVA), effect size was measured through partial eta-squared. In the case of Pearson’s correlation (R), the value of the correlation is to be taken as the measure of the effect size

| Analysis | Conditions | Statistical test | Result | DF | p-value | Effect size |
| --- | --- | --- | --- | --- | --- | --- |
| Pairwise decoding (average) | Latin, experts | t-test | 8.58 | 5 | < 0.001 | D = 3.5 |
|  | Latin, controls | t-test | 8.57 | 9 | < 0.001 | D = 2.7 |
|  | Braille, experts | t-test | 4.48 | 5 | 0.006 | D = 1.8 |
|  | Braille, controls | t-test | 1.45 | 9 | 0.1 | D = 0.5 |
|  | Latin, experts – Latin, controls | t-test | 1.05 | 11 | 0.31 | D = 0.5 |
|  | Latin, experts – Braille, experts | Paired t-test | 3.26 | 5 | 0.03 | D = 1.9 |
|  | Latin, experts – Braille, controls | t-test | 7.39 | 11 | < 0.001 | D = 4.4 |
|  | Latin, controls – Braille, experts | t-test | -2.51 | 11 | 0.034 | D = 1.2 |
|  | Latin, controls – Braille, controls | Paired t-test | 8.83 | 10 | < 0.001 | D = 2.9 |
|  | Braille, experts – Braille, controls | t-test | 3.50 | 7 | 0.017 | D = 2 |
| Within Braille ANOVA | Effect of group | rmANOVA | 16.01 | 1 | 0.001 | $\eta^2_p = 0.53$ |
| | Effect of conditions | rmANOVA | 1.72 | 5 | 0.14 | $\eta^2_p = 0.11$ |
| | Interaction group and condition | rmANOVA | 0.72 | 5 | 0.61 | $\eta^2_p = 0.05$ |
| Within Latin ANOVA | Effect of group | rmANOVA | 1.06 | 1 | 0.32 | $\eta^2_p = 0.07$ |
| | Effect of conditions | rmANOVA | 35.61 | 5 | < 0.001 | $\eta^2_p = 0.72$ |
| | Interaction group and condition | rmANOVA | 2.94 | 5 | 0.02 | $\eta^2_p = 0.17$ |
| RSA | Latin, experts – Latin, controls | Pearson's R | 0.75 | NA | < 0.001 | NA |
|  | Latin, experts – Braille, experts | Pearson's R | 0.56 | NA | < 0.001 | NA |
|  | Latin, experts – Braille, controls | Pearson's R | 0.19 | NA | 0.07 | NA |
|  | Latin, controls – Braille, experts | Pearson's R | 0.55 | NA | < 0.001 | NA |
|  | Latin, controls – Braille, controls | Pearson's R | 0.14 | NA | 0.13 | NA |
|  | Braille, experts – Braille, controls | Pearson's R | 0.14 | NA | 0.15 | NA |
|  | Latin, experts – Model | Pearson's R | 0.51 | NA | 0.002 | NA |
|  | Braille, experts – Model | Pearson's R | 0.34 | NA | 0.052 | NA |
|  | Latin, controls – Model | Pearson's R | 0.61 | NA | < 0.001 | NA |
|  | Braille, controls – Model | Pearson's R | -0.06 | NA | 0.68 | NA |
| Cross-script decoding | Average | t-test | 1.56 | 5 | 0.16 | D = 0.6 |
|  | Real words – Pseudo-words | t-test | 4.39 | 5 | 0.025 | D = 1.8 |
|  | Real words – non-words | t-test | 0.37 | 5 | 0.42 | D = 0.2 |
|  | Real words – fake script | t-test | 0.68 | 5 | 0.16 | D = 0.7 |
|  | Pseudo-words – non-words | t-test | -0.75 | 5 | 0.76 | D = 0.3 |
|  | Pseudo-words – fake script | t-test | 1.38 | 5 | 0.16 | D = 0.6 |
|  | Non-words – fake script | t-test | 1.57 | 5 | 0.16 | D = 0.6 |

**Table 4-2.** Post-hoc t-tests in left-LO. For each effect in the repeated measures ANOVA (rmANOVA), we contrasted the different comparisons to identify the causes of the interaction effects. In Latin script, results are averaged over the levels of ‘group’. P-values are adjusted for comparing a family of 15. In Braille script, p-values are adjusted for comparing a family of 66.

| Script | Effect | Comparison | Comparison contrasted | Mean Difference | SE | t | Cohen's d | p<sub>bonf</sub> |
| --- | --- | --- | --- | --- | --- | --- | --- | --- |
| Latin | Condition | RW-PW | RW-NW | -0.253 | 0.04 | -6.349 | -1.877 | < .001 |
|  |  |  | RW-FS | -0.394 | 0.04 | -9.907 | -2.929 | < .001 |
|  |  |  | PW-NW | -0.078 | 0.04 | -1.953 | -0.578 | 0.821 |
|  |  |  | PW-FS | -0.383 | 0.04 | -9.628 | -2.847 | < .001 |
|  |  |  | NW-FS | -0.353 | 0.04 | -8.86 | -2.62 | < .001 |
|  |  | RW-NW | RW-FS | -0.142 | 0.04 | -3.558 | -1.052 | 0.01 |
|  |  |  | PW-NW | 0.175 | 0.04 | 4.395 | 1.3 | < .001 |
|  |  |  | PW-FS | -0.131 | 0.04 | -3.279 | -0.97 | 0.024 |
|  |  |  | NW-FS | -0.1 | 0.04 | -2.512 | -0.743 | 0.215 |
|  |  | RW-FS | PW-NW | 0.317 | 0.04 | 7.953 | 2.352 | < .001 |
|  |  |  | PW-FS | 0.011 | 0.04 | 0.279 | 0.083 | 1 |
|  |  |  | NW-FS | 0.042 | 0.04 | 1.047 | 0.309 | 1 |
|  |  | PW-NW | PW-FS | -0.306 | 0.04 | -7.674 | -2.269 | < .001 |
|  |  |  | NW-FS | -0.275 | 0.04 | -6.907 | -2.042 | < .001 |
|  |  | PW-FS | NW-FS | 0.031 | 0.04 | 0.767 | 0.227 | 1 |
| Braille | Grp*Cond. | experts, RW-PW | novices, RW-PW | 0.042 | 0.073 | 0.57 | 0.295 | 1 |
|  |  |  | experts, RW-NW | -0.125 | 0.081 | -1.54 | -0.884 | 1 |
|  |  |  | novices, RW-NW | 0.058 | 0.073 | 0.799 | 0.412 | 1 |
|  |  |  | experts, RW-FS | -0.139 | 0.081 | -1.711 | -0.982 | 1 |
|  |  |  | novices, RW-FS | 0.025 | 0.073 | 0.342 | 0.177 | 1 |
|  |  |  | experts, PW-NW | -0.042 | 0.081 | -0.513 | -0.295 | 1 |
|  |  |  | novices, PW-NW | 0.017 | 0.073 | 0.228 | 0.118 | 1 |
|  |  |  | experts, PW-FS | -0.208 | 0.081 | -2.566 | -1.473 | 0.82 |
|  |  |  | novices, PW-FS | -0.033 | 0.073 | -0.456 | -0.236 | 1 |
|  |  |  | experts, NW-FS | -0.097 | 0.081 | -1.197 | -0.687 | 1 |
|  |  |  | novices, NW-FS | 0.017 | 0.073 | 0.228 | 0.118 | 1 |
|  |  | novices, RW-PW | experts, RW-NW | -0.167 | 0.073 | -2.282 | -1.178 | 1 |
|  |  |  | novices, RW-NW | 0.017 | 0.063 | 0.265 | 0.118 | 1 |
|  |  |  | experts, RW-FS | -0.181 | 0.073 | -2.472 | -1.277 | 1 |
|  |  |  | novices, RW-FS | -0.017 | 0.063 | -0.265 | -0.118 | 1 |
|  |  |  | experts, PW-NW | -0.083 | 0.073 | -1.141 | -0.589 | 1 |
|  |  |  | novices, PW-NW | -0.025 | 0.063 | -0.398 | -0.177 | 1 |
|  |  |  | experts, PW-FS | -0.25 | 0.073 | -3.423 | -1.768 | 0.063 |
|  |  |  | novices, PW-FS | -0.075 | 0.063 | -1.193 | -0.53 | 1 |
|  |  |  | experts, NW-FS | -0.139 | 0.073 | -1.902 | -0.982 | 1 |
|  |  |  | novices, NW-FS | -0.025 | 0.063 | -0.398 | -0.177 | 1 |
|  |  | experts, RW-NW | novices, RW-NW | 0.183 | 0.073 | 2.51 | 1.296 | 0.923 |
|  |  |  | experts, RW-FS | -0.014 | 0.081 | -0.171 | -0.098 | 1 |
|  |  |  | novices, RW-FS | 0.15 | 0.073 | 2.054 | 1.061 | 1 |
|  |  |  | experts, PW-NW | 0.083 | 0.081 | 1.026 | 0.589 | 1 |
|  |  |  | novices, PW-NW | 0.142 | 0.073 | 1.94 | 1.002 | 1 |
|  |  |  | experts, PW-FS | -0.083 | 0.081 | -1.026 | -0.589 | 1 |
|  |  |  | novices, PW-FS | 0.092 | 0.073 | 1.255 | 0.648 | 1 |
|  |  |  | experts, NW-FS | 0.028 | 0.081 | 0.342 | 0.196 | 1 |
|  |  |  | novices, NW-FS | 0.142 | 0.073 | 1.94 | 1.002 | 1 |
|  |  | novices, RW-NW | experts, RW-FS | -0.197 | 0.073 | -2.7 | -1.394 | 0.553 |
|  |  |  | novices, RW-FS | -0.033 | 0.063 | -0.53 | -0.236 | 1 |
|  |  |  | experts, PW-NW | -0.1 | 0.073 | -1.369 | -0.707 | 1 |
|  |  |  | novices, PW-NW | -0.042 | 0.063 | -0.663 | -0.295 | 1 |
|  |  |  | experts, PW-FS | -0.267 | 0.073 | -3.651 | -1.885 | 0.03 |
|  |  |  | novices, PW-FS | -0.092 | 0.063 | -1.458 | -0.648 | 1 |
|  |  |  | experts, NW-FS | -0.156 | 0.073 | -2.13 | -1.1 | 1 |
|  |  |  | novices, NW-FS | -0.042 | 0.063 | -0.663 | -0.295 | 1 |
|  |  | experts, RW-FS | novices, RW-FS | 0.164 | 0.073 | 2.244 | 1.159 | 1 |
|  |  |  | experts, PW-NW | 0.097 | 0.081 | 1.197 | 0.687 | 1 |
|  |  |  | novices, PW-NW | 0.156 | 0.073 | 2.13 | 1.1 | 1 |
|  |  |  | experts, PW-FS | -0.069 | 0.081 | -0.855 | -0.491 | 1 |
|  |  |  | novices, PW-FS | 0.106 | 0.073 | 1.445 | 0.746 | 1 |
|  |  |  | experts, NW-FS | 0.042 | 0.081 | 0.513 | 0.295 | 1 |
|  |  |  | novices, NW-FS | 0.156 | 0.073 | 2.13 | 1.1 | 1 |
|  |  | novices, RW-FS | experts, PW-NW | -0.067 | 0.073 | -0.913 | -0.471 | 1 |
|  |  |  | novices, PW-NW | -0.008 | 0.063 | -0.133 | -0.059 | 1 |
|  |  |  | experts, PW-FS | -0.233 | 0.073 | -3.195 | -1.65 | 0.13 |
|  |  |  | novices, PW-FS | -0.058 | 0.063 | -0.928 | -0.412 | 1 |
|  |  |  | experts, NW-FS | -0.122 | 0.073 | -1.673 | -0.864 | 1 |
|  |  |  | novices, NW-FS | -0.008 | 0.063 | -0.133 | -0.059 | 1 |
|  |  | experts, PW-NW | novices, PW-NW | 0.058 | 0.073 | 0.799 | 0.412 | 1 |
|  |  |  | experts, PW-FS | -0.167 | 0.081 | -2.053 | -1.178 | 1 |
|  |  |  | novices, PW-FS | 0.008 | 0.073 | 0.114 | 0.059 | 1 |
|  |  |  | experts, NW-FS | -0.056 | 0.081 | -0.684 | -0.393 | 1 |
|  |  |  | novices, NW-FS | 0.058 | 0.073 | 0.799 | 0.412 | 1 |
|  |  | novices, PW-NW | experts, PW-FS | -0.225 | 0.073 | -3.081 | -1.591 | 0.184 |
|  |  |  | novices, PW-FS | -0.05 | 0.063 | -0.795 | -0.354 | 1 |
|  |  |  | experts, NW-FS | -0.114 | 0.073 | -1.559 | -0.805 | 1 |
| | | | novices, NW-FS | $1.1 \times 10^{-16}$ | 0.063 | $2 \times 10^{-15}$ | $2 \times 10^{-15}$ | 1 |
|  |  | experts, PW-FS | novices, PW-FS | 0.175 | 0.073 | 2.396 | 1.237 | 1 |
|  |  |  | experts, NW-FS | 0.111 | 0.081 | 1.369 | 0.786 | 1 |
|  |  |  | novices, NW-FS | 0.225 | 0.073 | 3.081 | 1.591 | 0.184 |
|  |  | novices, PW-FS | experts, NW-FS | -0.064 | 0.073 | -0.875 | -0.452 | 1 |
|  |  |  | novices, NW-FS | 0.05 | 0.063 | 0.795 | 0.354 | 1 |
|  |  | experts, NW-FS | novices, NW-FS | 0.114 | 0.073 | 1.559 | 0.805 | 1 |

**Table 4-3.** Statistical tests in right-LO. Full list of statistical tests performed over the multivariate results of l-PosTemp. Each analysis is detailed with the conditions tested and the statistical test. Statistical tests detail the result, the degrees of freedom (DF), the p-value, and a measure of the effect size. For t-test, effect size was measured through Cohen’s D. For repeated measures ANOVA (rmANOVA), effect size was measured through partial eta-squared. In the case of Pearson’s correlation (R), the value of the correlation is to be taken as the measure of the effect size.

| Analysis | Conditions | Statistical test | Result | DF | p-value | Effect size |
| --- | --- | --- | --- | --- | --- | --- |
| Pairwise decoding (average) | Latin, experts | t-test | 9.63 | 5 | < 0.001 | D = 3.9 |
|  | Latin, controls | t-test | 5.87 | 10 | < 0.001 | D = 1.7 |
|  | Braille, experts | t-test | 3.88 | 5 | 0.012 | D = 1.5 |
|  | Braille, controls | t-test | 1.01 | 10 | 0.21 | D = 0.3 |
|  | Latin, experts – Latin, controls | t-test | 0.45 | 15 | 0.66 | D = 0.2 |
|  | Latin, experts – Braille, experts | Paired t-test | 1.85 | 5 | 0.17 | D = 1.1 |
|  | Latin, experts – Braille, controls | t-test | 6.64 | 12 | < 0.001 | D = 3.2 |
|  | Latin, controls – Braille, experts | t-test | -1.25 | 13 | 0.25 | D = 0.6 |
|  | Latin, controls – Braille, controls | Paired t-test | 6.57 | 10 | < 0.001 | D = 1.8 |
|  | Braille, experts – Braille, controls | t-test | 2.92 | 8 | 0.032 | D = 1.6 |
| Within Braille ANOVA | Effect of group | rmANOVA | 10.23 | 1 | 0.006 | $\eta^2_p = 0.41$ |
| | Effect of conditions | rmANOVA | 1.89 | 5 | 0.11 | $\eta^2_p = 0.11$ |
| | Interaction group and condition | rmANOVA | 1.41 | 5 | 0.23 | $\eta^2_p = 0.09$ |
| Within Latin ANOVA | Effect of group | rmANOVA | 0.14 | 1 | 0.72 | $\eta^2_p = 0.01$ |
| | Effect of conditions | rmANOVA | 35.70 | 5 | < 0.001 | $\eta^2_p = 0.70$ |
| | Interaction group and condition | rmANOVA | 0.90 | 5 | 0.49 | $\eta^2_p = 0.06$ |
| RSA | Latin, experts – Latin, controls | Pearson’s R | 0.83 | NA | < 0.001 | NA |
|  | Latin, experts – Braille, experts | Pearson’s R | 0.65 | NA | < 0.001 | NA |
|  | Latin, experts – Braille, controls | Pearson’s R | 0.06 | NA | 0.36 | NA |
|  | Latin, controls – Braille, experts | Pearson’s R | 0.64 | NA | < 0.001 | NA |
|  | Latin, controls – Braille, controls | Pearson’s R | 0.01 | NA | 0.52 | NA |
|  | Braille, experts – Braille, controls | Pearson’s R | 0.04 | NA | 0.44 | NA |
|  | Latin, experts – Model | Pearson’s R | 0.64 | NA | < 0.001 | NA |
|  | Braille, experts – Model | Pearson’s R | 0.53 | NA | 0.002 | NA |
|  | Latin, controls – Model | Pearson’s R | 0.44 | NA | < 0.001 | NA |
|  | Braille, controls – Model | Pearson’s R | 0.04 | NA | 0.46 | NA |
| Cross-script decoding | Average | t-test | 2.35 | 5 | 0.1 | D = 1 |
|  | Real words – Pseudo-words | t-test | -0.36 | 5 | 0.63 | D = 0.1 |
|  | Real words – non-words | t-test | -0.92 | 5 | 0.1 | D = 0.4 |
|  | Real words – fake script | t-test | 1.83 | 5 | 0.1 | D = 0.7 |
|  | Pseudo-words – non-words | t-test | 1.75 | 5 | 0.1 | D = 0.7 |
|  | Pseudo-words – fake script | t-test | 1.94 | 5 | 0.09 | D = 0.8 |
|  | Non-words – fake script | t-test | 2.28 | 5 | 0.1 | D = 0.9 |

**Table 4-4.**
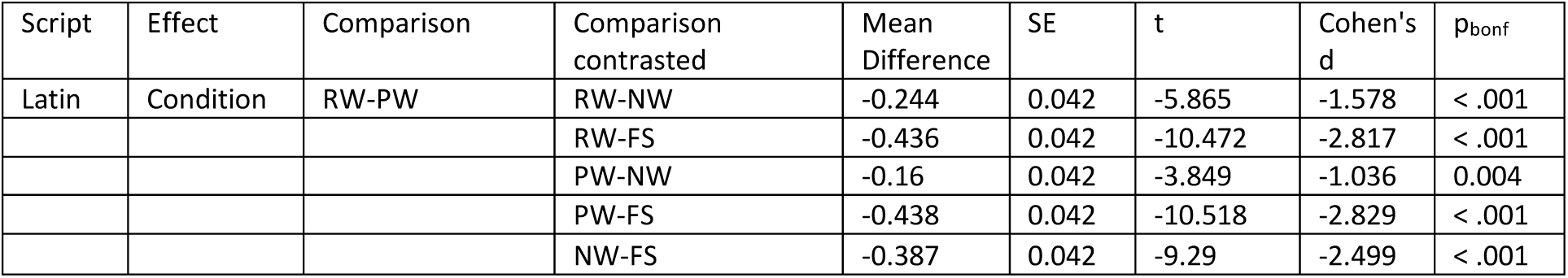
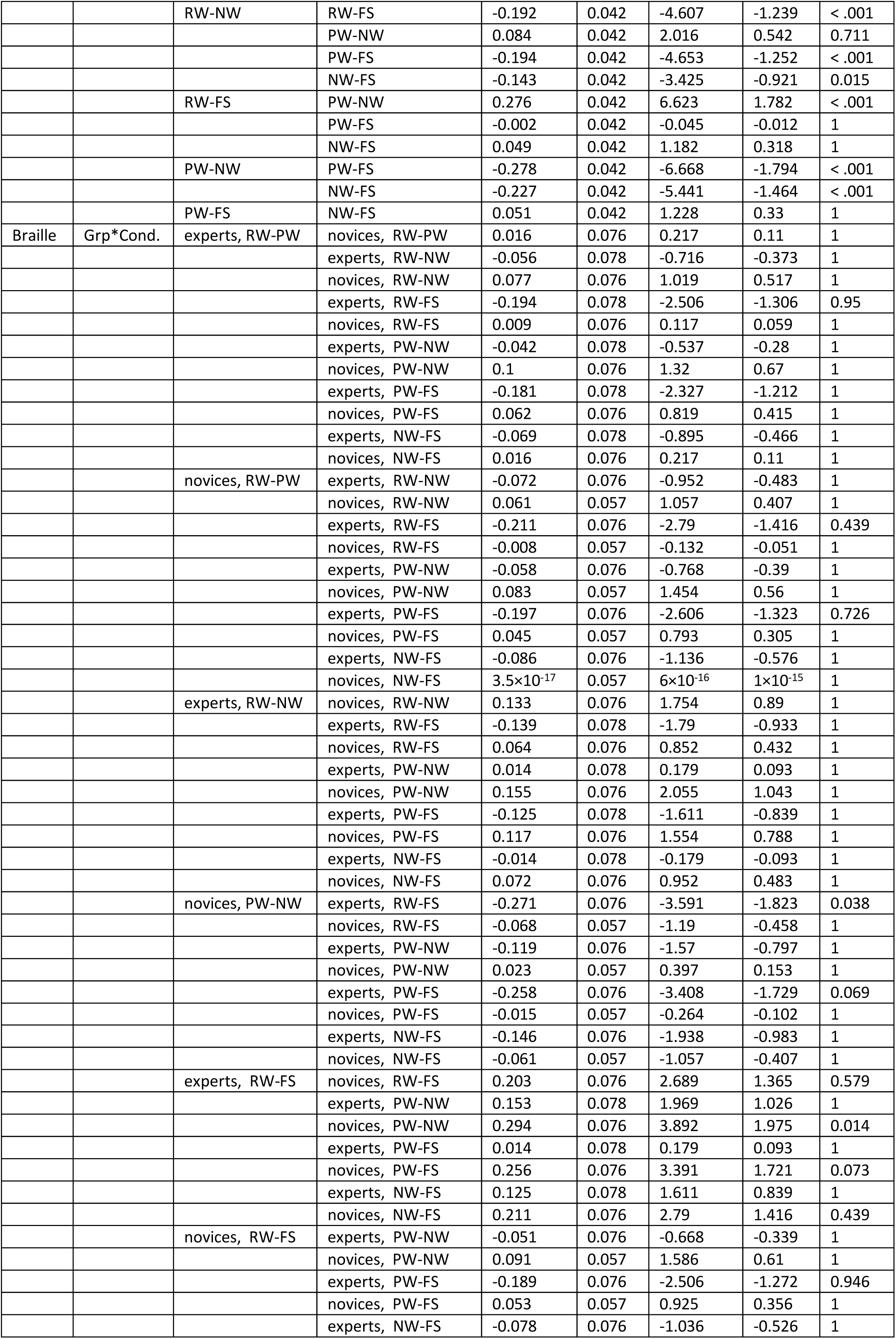
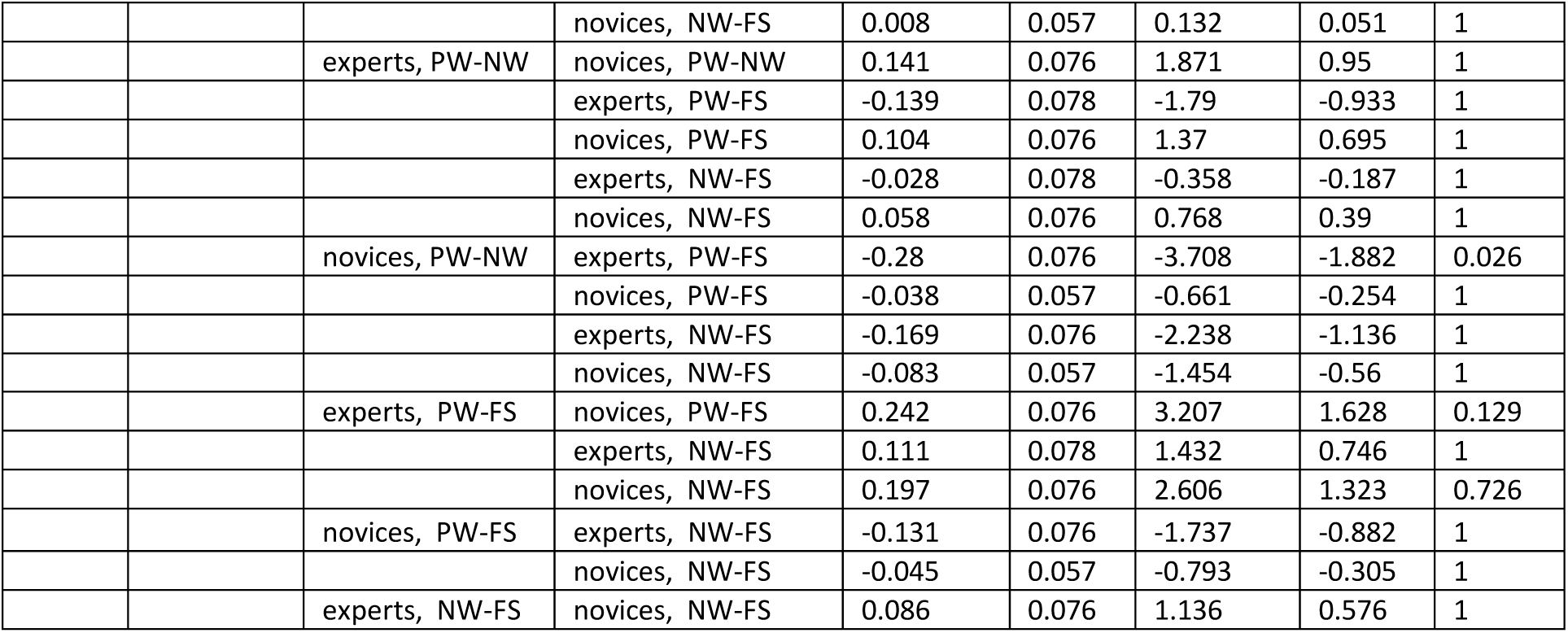
Post-hoc t-tests in right-LO. For each effect in the repeated measures ANOVA (rmANOVA), we contrasted the different comparisons to identify the causes of the interaction effects. In Latin script, results are averaged over the levels of ‘group’. P-values are adjusted for comparing a family of 15. In Braille script, p-values are adjusted for comparing a family of 66.

| Script | Effect | Comparison | Comparison contrasted | Mean Difference | SE | t | Cohen's d | p<sub>bonf</sub> |
| --- | --- | --- | --- | --- | --- | --- | --- | --- |
| Latin | Condition | RW-PW | RW-NW | -0.244 | 0.042 | -5.865 | -1.578 | < .001 |
|  |  |  | RW-FS | -0.436 | 0.042 | -10.472 | -2.817 | < .001 |
|  |  |  | PW-NW | -0.16 | 0.042 | -3.849 | -1.036 | 0.004 |
|  |  |  | PW-FS | -0.438 | 0.042 | -10.518 | -2.829 | < .001 |
|  |  |  | NW-FS | -0.387 | 0.042 | -9.29 | -2.499 | < .001 |
|  |  | RW-NW | RW-FS | -0.192 | 0.042 | -4.607 | -1.239 | < .001 |
|  |  |  | PW-NW | 0.084 | 0.042 | 2.016 | 0.542 | 0.711 |
|  |  |  | PW-FS | -0.194 | 0.042 | -4.653 | -1.252 | < .001 |
|  |  |  | NW-FS | -0.143 | 0.042 | -3.425 | -0.921 | 0.015 |
|  |  | RW-FS | PW-NW | 0.276 | 0.042 | 6.623 | 1.782 | < .001 |
|  |  |  | PW-FS | -0.002 | 0.042 | -0.045 | -0.012 | 1 |
|  |  |  | NW-FS | 0.049 | 0.042 | 1.182 | 0.318 | 1 |
|  |  | PW-NW | PW-FS | -0.278 | 0.042 | -6.668 | -1.794 | < .001 |
|  |  |  | NW-FS | -0.227 | 0.042 | -5.441 | -1.464 | < .001 |
|  |  | PW-FS | NW-FS | 0.051 | 0.042 | 1.228 | 0.33 | 1 |
| Braille | Grp*Cond. | experts, RW-PW | novices, RW-PW | 0.016 | 0.076 | 0.217 | 0.11 | 1 |
|  |  |  | experts, RW-NW | -0.056 | 0.078 | -0.716 | -0.373 | 1 |
|  |  |  | novices, RW-NW | 0.077 | 0.076 | 1.019 | 0.517 | 1 |
|  |  |  | experts, RW-FS | -0.194 | 0.078 | -2.506 | -1.306 | 0.95 |
|  |  |  | novices, RW-FS | 0.009 | 0.076 | 0.117 | 0.059 | 1 |
|  |  |  | experts, PW-NW | -0.042 | 0.078 | -0.537 | -0.28 | 1 |
|  |  |  | novices, PW-NW | 0.1 | 0.076 | 1.32 | 0.67 | 1 |
|  |  |  | experts, PW-FS | -0.181 | 0.078 | -2.327 | -1.212 | 1 |
|  |  |  | novices, PW-FS | 0.062 | 0.076 | 0.819 | 0.415 | 1 |
|  |  |  | experts, NW-FS | -0.069 | 0.078 | -0.895 | -0.466 | 1 |
|  |  |  | novices, NW-FS | 0.016 | 0.076 | 0.217 | 0.11 | 1 |
|  |  | novices, RW-PW | experts, RW-NW | -0.072 | 0.076 | -0.952 | -0.483 | 1 |
|  |  |  | novices, RW-NW | 0.061 | 0.057 | 1.057 | 0.407 | 1 |
|  |  |  | experts, RW-FS | -0.211 | 0.076 | -2.79 | -1.416 | 0.439 |
|  |  |  | novices, RW-FS | -0.008 | 0.057 | -0.132 | -0.051 | 1 |
|  |  |  | experts, PW-NW | -0.058 | 0.076 | -0.768 | -0.39 | 1 |
|  |  |  | novices, PW-NW | 0.083 | 0.057 | 1.454 | 0.56 | 1 |
|  |  |  | experts, PW-FS | -0.197 | 0.076 | -2.606 | -1.323 | 0.726 |
|  |  |  | novices, PW-FS | 0.045 | 0.057 | 0.793 | 0.305 | 1 |
|  |  |  | experts, NW-FS | -0.086 | 0.076 | -1.136 | -0.576 | 1 |
| | | | novices, NW-FS | $3.5 \times 10^{-17}$ | 0.057 | $6 \times 10^{-16}$ | $1 \times 10^{-15}$ | 1 |
|  |  | experts, RW-NW | novices, RW-NW | 0.133 | 0.076 | 1.754 | 0.89 | 1 |
|  |  |  | experts, RW-FS | -0.139 | 0.078 | -1.79 | -0.933 | 1 |
|  |  |  | novices, RW-FS | 0.064 | 0.076 | 0.852 | 0.432 | 1 |
|  |  |  | experts, PW-NW | 0.014 | 0.078 | 0.179 | 0.093 | 1 |
|  |  |  | novices, PW-NW | 0.155 | 0.076 | 2.055 | 1.043 | 1 |
|  |  |  | experts, PW-FS | -0.125 | 0.078 | -1.611 | -0.839 | 1 |
|  |  |  | novices, PW-FS | 0.117 | 0.076 | 1.554 | 0.788 | 1 |
|  |  |  | experts, NW-FS | -0.014 | 0.078 | -0.179 | -0.093 | 1 |
|  |  |  | novices, NW-FS | 0.072 | 0.076 | 0.952 | 0.483 | 1 |
|  |  | novices, PW-NW | experts, RW-FS | -0.271 | 0.076 | -3.591 | -1.823 | 0.038 |
|  |  |  | novices, RW-FS | -0.068 | 0.057 | -1.19 | -0.458 | 1 |
|  |  |  | experts, PW-NW | -0.119 | 0.076 | -1.57 | -0.797 | 1 |
|  |  |  | novices, PW-NW | 0.023 | 0.057 | 0.397 | 0.153 | 1 |
|  |  |  | experts, PW-FS | -0.258 | 0.076 | -3.408 | -1.729 | 0.069 |
|  |  |  | novices, PW-FS | -0.015 | 0.057 | -0.264 | -0.102 | 1 |
|  |  |  | experts, NW-FS | -0.146 | 0.076 | -1.938 | -0.983 | 1 |
|  |  |  | novices, NW-FS | -0.061 | 0.057 | -1.057 | -0.407 | 1 |
|  |  | experts, RW-FS | novices, RW-FS | 0.203 | 0.076 | 2.689 | 1.365 | 0.579 |
|  |  |  | experts, PW-NW | 0.153 | 0.078 | 1.969 | 1.026 | 1 |
|  |  |  | novices, PW-NW | 0.294 | 0.076 | 3.892 | 1.975 | 0.014 |
|  |  |  | experts, PW-FS | 0.014 | 0.078 | 0.179 | 0.093 | 1 |
|  |  |  | novices, PW-FS | 0.256 | 0.076 | 3.391 | 1.721 | 0.073 |
|  |  |  | experts, NW-FS | 0.125 | 0.078 | 1.611 | 0.839 | 1 |
|  |  |  | novices, NW-FS | 0.211 | 0.076 | 2.79 | 1.416 | 0.439 |
|  |  | novices, RW-FS | experts, PW-NW | -0.051 | 0.076 | -0.668 | -0.339 | 1 |
|  |  |  | novices, PW-NW | 0.091 | 0.057 | 1.586 | 0.61 | 1 |
|  |  |  | experts, PW-FS | -0.189 | 0.076 | -2.506 | -1.272 | 0.946 |
|  |  |  | novices, PW-FS | 0.053 | 0.057 | 0.925 | 0.356 | 1 |
|  |  |  | experts, NW-FS | -0.078 | 0.076 | -1.036 | -0.526 | 1 |

|  |  |  |  |  |  |  |  |
| --- | --- | --- | --- | --- | --- | --- | --- |
|  |  | novices, NW-FS | 0.008 | 0.057 | 0.132 | 0.051 | 1 |
|  | experts, PW-NW | novices, PW-NW | 0.141 | 0.076 | 1.871 | 0.95 | 1 |
|  |  | experts, PW-FS | -0.139 | 0.078 | -1.79 | -0.933 | 1 |
|  |  | novices, PW-FS | 0.104 | 0.076 | 1.37 | 0.695 | 1 |
|  |  | experts, NW-FS | -0.028 | 0.078 | -0.358 | -0.187 | 1 |
|  |  | novices, NW-FS | 0.058 | 0.076 | 0.768 | 0.39 | 1 |
|  | novices, PW-NW | experts, PW-FS | -0.28 | 0.076 | -3.708 | -1.882 | 0.026 |
|  |  | novices, PW-FS | -0.038 | 0.057 | -0.661 | -0.254 | 1 |
|  |  | experts, NW-FS | -0.169 | 0.076 | -2.238 | -1.136 | 1 |
|  |  | novices, NW-FS | -0.083 | 0.057 | -1.454 | -0.56 | 1 |
|  | experts, PW-FS | novices, PW-FS | 0.242 | 0.076 | 3.207 | 1.628 | 0.129 |
|  |  | experts, NW-FS | 0.111 | 0.078 | 1.432 | 0.746 | 1 |
|  |  | novices, NW-FS | 0.197 | 0.076 | 2.606 | 1.323 | 0.726 |
|  | novices, PW-FS | experts, NW-FS | -0.131 | 0.076 | -1.737 | -0.882 | 1 |
|  |  | novices, NW-FS | -0.045 | 0.057 | -0.793 | -0.305 | 1 |
|  | experts, NW-FS | novices, NW-FS | 0.086 | 0.076 | 1.136 | 0.576 | 1 |

**Table 5-1.** Statistical tests in V1. Full list of statistical tests performed over the multivariate results of l-PosTemp. Each analysis is detailed with the conditions tested and the statistical test. Statistical tests detail the result, the degrees of freedom (DF), the p-value, and a measure of the effect size. For t-test, effect size was measured through Cohen’s D. For repeated measures ANOVA (rmANOVA), effect size was measured through partial eta-squared. In the case of Pearson’s correlation (R), the value of the correlation is to be taken as the measure of the effect size

| Analysis | Conditions | Statistical test | Result | DF | p-value | Effect size |
| --- | --- | --- | --- | --- | --- | --- |
| Pairwise decoding (average) | Latin, experts | t-test | 4.35 | 5 | 0.018 | D = 1.8 |
|  | Latin, controls | t-test | 3.15 | 10 | 0.018 | D = 0.9 |
|  | Braille, experts | t-test | 3.93 | 5 | 0.018 | D = 1.6 |
|  | Braille, controls | t-test | 2.45 | 10 | 0.28 | D = 0.7 |
|  | Latin, experts – Latin, controls | t-test | 0.71 | 13 | 0.53 | D = 0.3 |
|  | Latin, experts – Braille, experts | Paired t-test | 1.60 | 5 | 0.21 | D = 0.6 |
|  | Latin, experts – Braille, controls | t-test | 3.47 | 6 | 0.027 | D = 2.2 |
|  | Latin, controls – Braille, experts | t-test | 0.23 | 15 | 0.82 | D = 0.1 |
|  | Latin, controls – Braille, controls | Paired t-test | 3.01 | 10 | 0.027 | D = 0.7 |
|  | Braille, experts – Braille, controls | t-test | 2.78 | 6 | 0.04 | D = 1.7 |
| Within Braille ANOVA | Effect of group | rmANOVA | 11.12 | 1 | 0.005 | $\eta^2_p = 0.43$ |
| | Effect of conditions | rmANOVA | 4.05 | 5 | 0.003 | $\eta^2_p = 0.21$ |
| | Interaction group and condition | rmANOVA | 2.33 | 5 | 0.051 | $\eta^2_p = 0.13$ |
| Within Latin ANOVA | Effect of group | rmANOVA | 0.41 | 1 | 0.53 | $\eta^2_p = 0.03$ |
| | Effect of conditions | rmANOVA | 11.09 | 5 | < 0.001 | $\eta^2_p = 0.43$ |
| | Interaction group and condition | rmANOVA | 1.32 | 5 | 0.26 | $\eta^2_p = 0.08$ |
| RSA | Latin, experts – Latin, controls | Pearson’s R | 0.56 | NA | < 0.001 | NA |
|  | Latin, experts – Braille, experts | Pearson’s R | 0.50 | NA | < 0.001 | NA |
|  | Latin, experts – Braille, controls | Pearson’s R | 0.20 | NA | 0.06 | NA |
|  | Latin, controls – Braille, experts | Pearson’s R | 0.49 | NA | < 0.001 | NA |
|  | Latin, controls – Braille, controls | Pearson’s R | 0.22 | NA | 0.02 | NA |
|  | Braille, experts – Braille, controls | Pearson’s R | 0.18 | NA | 0.07 | NA |
|  | Latin, experts – Model | Pearson’s R | 0.68 | NA | < 0.001 | NA |
|  | Braille, experts – Model | Pearson’s R | 0.25 | NA | 0.09 | NA |
|  | Latin, controls – Model | Pearson’s R | 0.28 | NA | 0.025 | NA |
|  | Braille, controls – Model | Pearson’s R | 0.14 | NA | 0.15 | NA |
| Cross-script decoding | Average | t-test | -0.36 | 5 | 0.86 | D = 0.1 |
|  | Real words – Pseudo-words | t-test | -0.79 | 5 | 0.86 | D = 0.3 |
|  | Real words – non-words | t-test | -1.21 | 5 | 0.86 | D = 0.5 |
|  | Real words – fake script | t-test | 0.73 | 5 | 0.71 | D = 0.3 |
|  | Pseudo-words – non-words | t-test | -1.03 | 5 | 0.86 | D = 0.4 |
|  | Pseudo-words – fake script | t-test | 0.54 | 5 | 0.71 | D = 0.2 |
|  | Non-words – fake script | t-test | 0.88 | 5 | 0.71 | D = 0.3 |

**Table 5-2.**
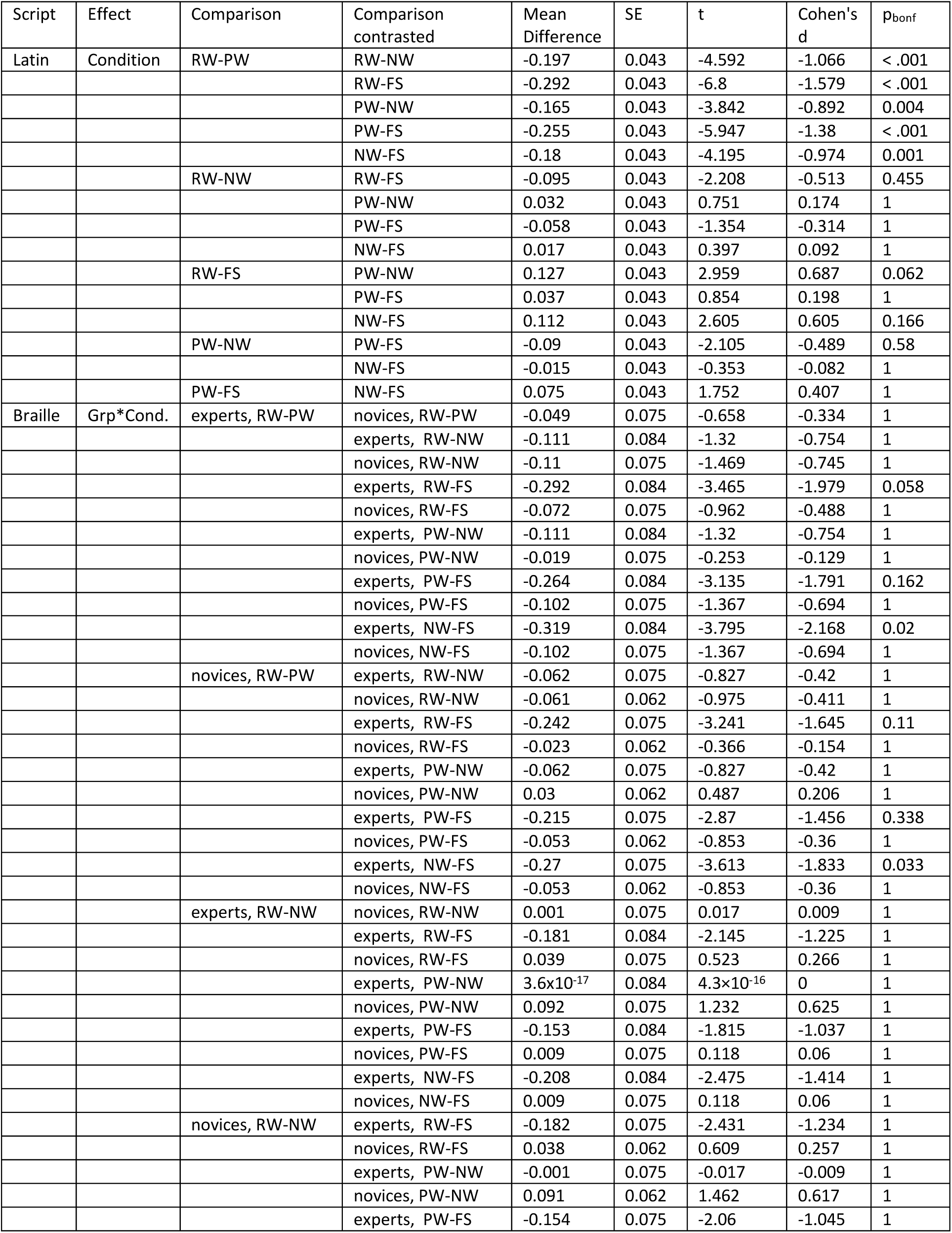
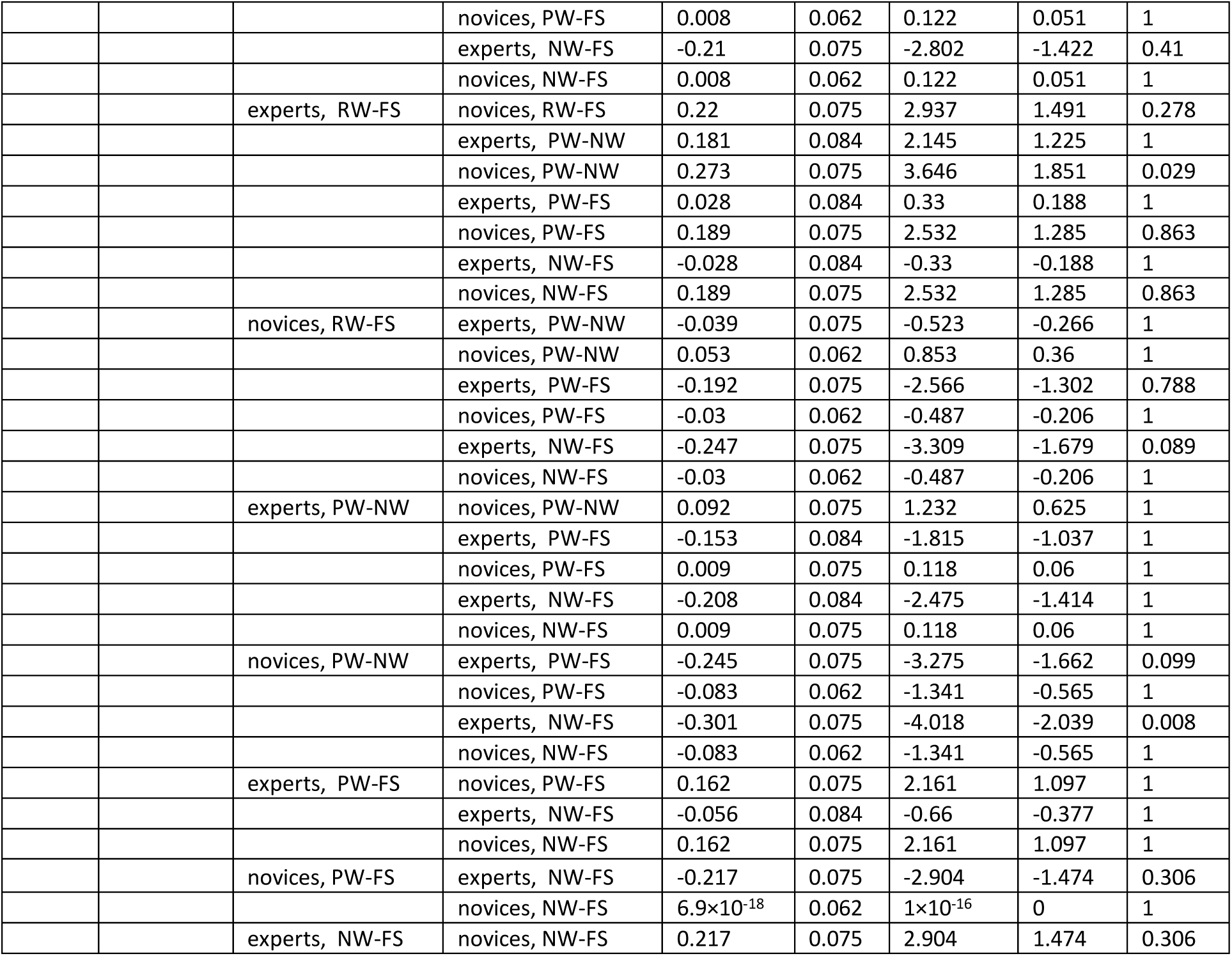
Post-hoc t-tests in V1. For each effect in the repeated measures ANOVA (rmANOVA), we contrasted the different comparisons to identify the causes of the interaction effects. In Latin script, results are averaged over the levels of ‘group’. P-values are adjusted for comparing a family of 15. In Braille script, p-values are adjusted for comparing a family of 66.

**Table 6-1.** Statistical tests in l-PosTemp. Full list of statistical tests performed over the multivariate results of l-PosTemp. Each analysis is detailed with the conditions tested and the statistical test. Statistical tests detail the result, the degrees of freedom (DF), the p-value, and a measure of the effect size. For t-test, effect size was measured through Cohen’s D. For repeated measures ANOVA (rmANOVA), effect size was measured through partial eta-squared. In the case of Pearson’s correlation (R), the value of the correlation is to be taken as the measure of the effect size.

| Analysis | Conditions | Statistical test | Result | DF | p-value | Effect size |
| --- | --- | --- | --- | --- | --- | --- |
| Pairwise decoding (average) | Latin, experts | t-test | 10.12 | 4 | < 0.001 | D = 4.5 |
|  | Latin, controls | t-test | 5.35 | 9 | < 0.001 | D = 1.7 |
|  | Braille, experts | t-test | 16.66 | 4 | < 0.001 | D = 7.4 |
|  | Braille, controls | t-test | 0.92 | 9 | 0.24 | D = 0.3 |
|  | Latin, experts – Latin, controls | t-test | 1.14 | 12 | 0.30 | D = 0.5 |
|  | Latin, experts – Braille, experts | Paired t-test | -1.08 | 4 | 0.34 | D = 0.4 |
|  | Latin, experts – Braille, controls | t-test | 6.96 | 10 | < 0.001 | D = 3.5 |
|  | Latin, controls – Braille, experts | t-test | 1.90 | 12 | 0.11 | D = 0.7 |
|  | Latin, controls – Braille, controls | Paired t-test | 7.88 | 9 | < 0.001 | D = 1.5 |
|  | Braille, experts – Braille, controls | t-test | 9.32 | 12 | < 0.001 | D = 4.2 |
| Within Braille ANOVA | Effect of group | rmANOVA | 58.42 | 1 | < 0.001 | $\eta^2_p = 0.82$ |
| | Effect of conditions | rmANOVA | 4.30 | 5 | 0.002 | $\eta^2_p = 0.25$ |
| | Interaction group and condition | rmANOVA | 2.27 | 5 | 0.06 | $\eta^2_p = 0.15$ |
| Within Latin ANOVA | Effect of group | rmANOVA | 0.83 | 1 | 0.38 | $\eta^2_p = 0.06$ |
| | Effect of conditions | rmANOVA | 4.99 | 5 | < 0.001 | $\eta^2_p = 0.28$ |
| | Interaction group and condition | rmANOVA | 1.27 | 5 | 0.31 | $\eta^2_p = 0.09$ |
| RSA | Latin, experts – Latin, controls | Pearson’s R | 0.43 | NA | < 0.001 | NA |
|  | Latin, experts – Braille, experts | Pearson’s R | 0.52 | NA | < 0.001 | NA |
|  | Latin, experts – Braille, controls | Pearson’s R | 0.15 | NA | 0.13 | NA |
|  | Latin, controls – Braille, experts | Pearson's R | 0.44 | NA | < 0.001 | NA |
|  | Latin, controls – Braille, controls | Pearson's R | 0.18 | NA | 0.06 | NA |
|  | Braille, experts – Braille, controls | Pearson's R | 0.15 | NA | 0.13 | NA |
|  | Latin, experts – Model | Pearson's R | 0.18 | NA | 0.21 | NA |
|  | Braille, experts – Model | Pearson's R | 0.27 | NA | 0.13 | NA |
|  | Latin, controls – Model | Pearson's R | 0.26 | NA | 0.06 | NA |
|  | Braille, controls – Model | Pearson's R | -0.06 | NA | 0.65 | NA |
| Cross-script decoding | Average | t-test | 4.08 | 4 | 0.025 | D = 1.8 |
|  | Real words – Pseudo-words | t-test | 2.24 | 4 | 0.052 | D = 1 |
|  | Real words – non-words | t-test | 4.49 | 4 | 0.025 | D = 2 |
|  | Real words – fake script | t-test | 0.29 | 4 | 0.39 | D = 0.1 |
|  | Pseudo-words – non-words | t-test | 3.64 | 4 | 0.025 | D = 1.6 |
|  | Pseudo-words – fake script | t-test | 2.69 | 4 | 0.04 | D = 1.2 |
|  | Non-words – fake script | t-test | 3.35 | 4 | 0.025 | D = 1.5 |

**Table 6-2.**
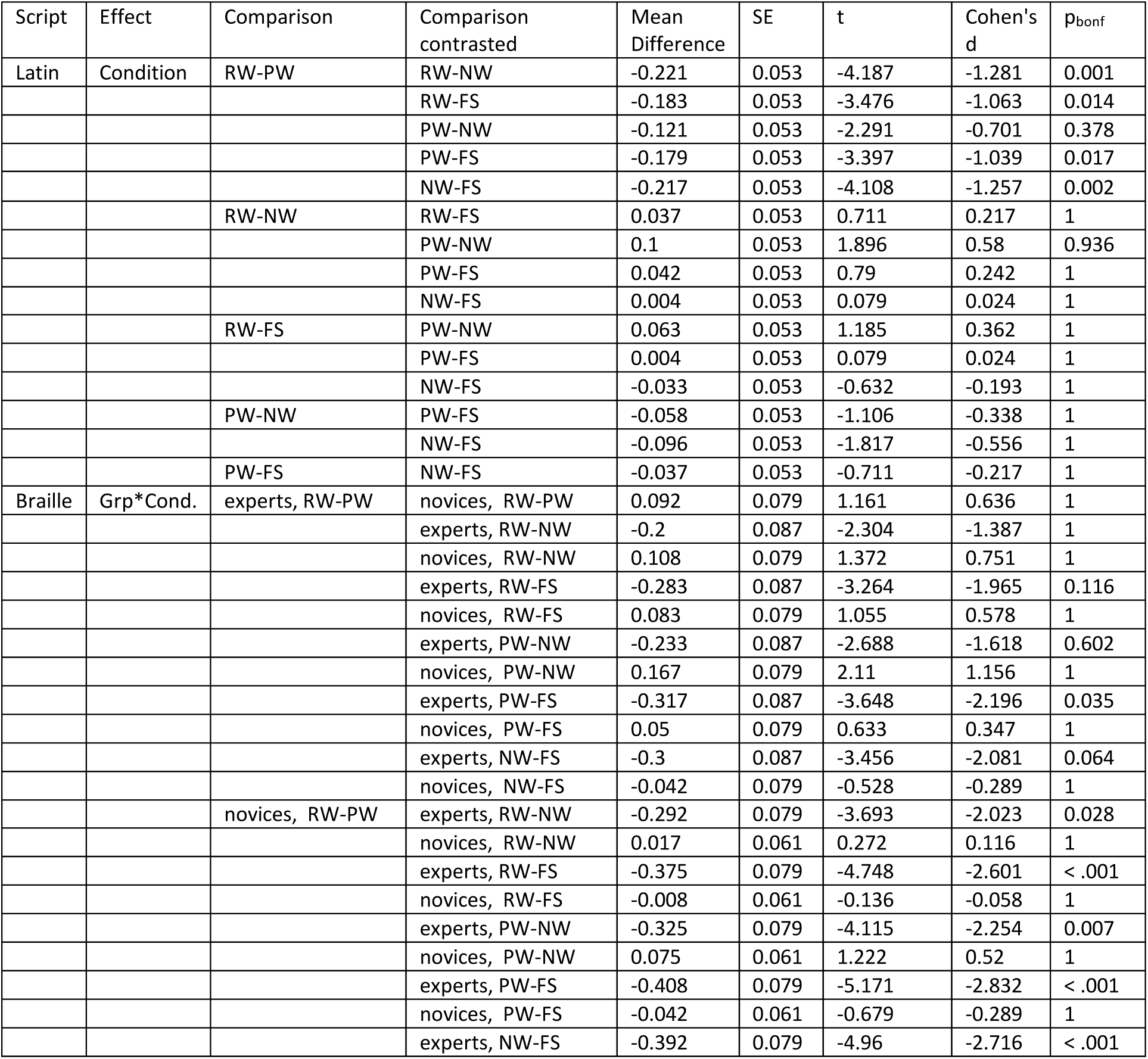
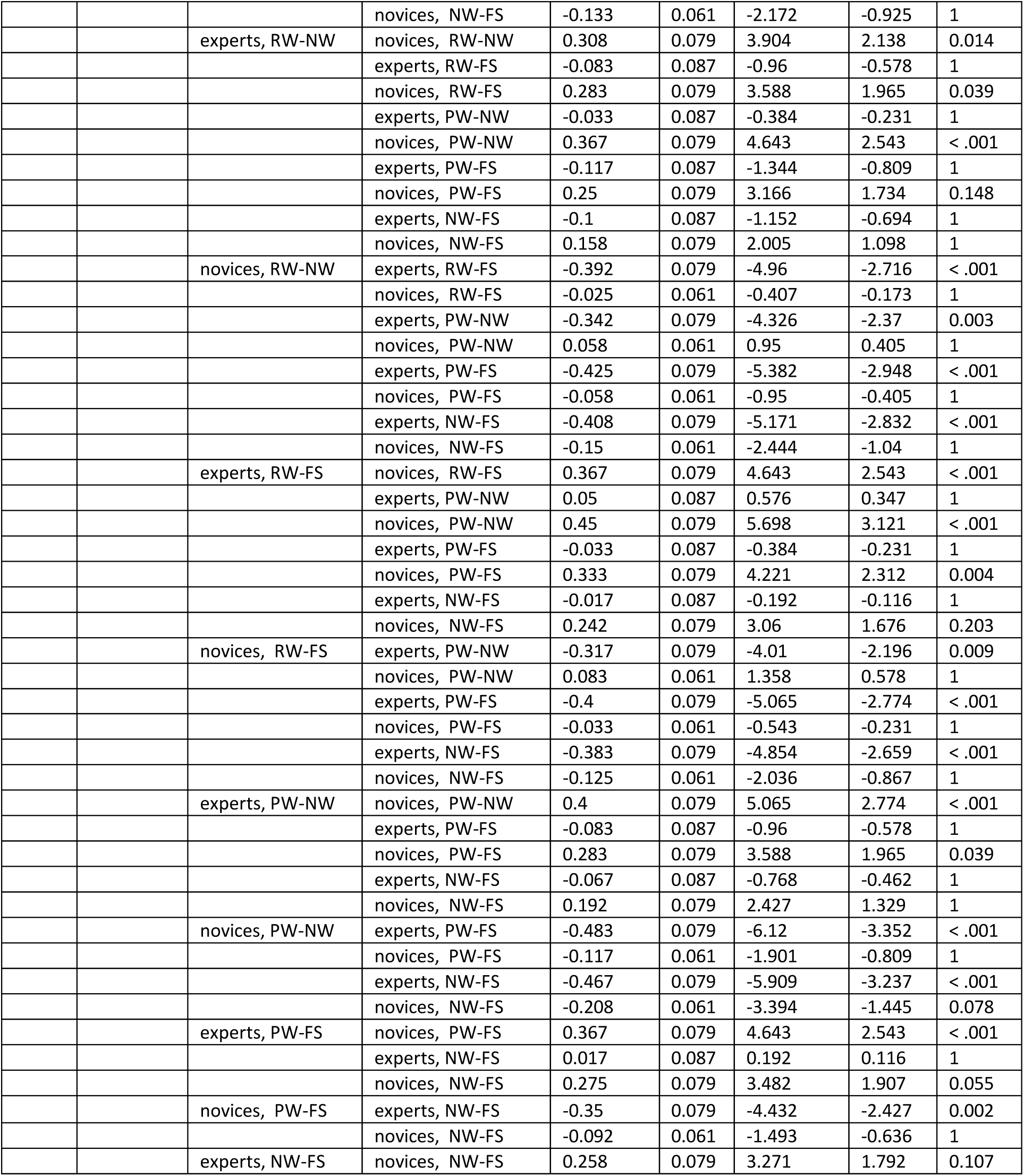
Post-hoc t-tests in l-PosTemp. For each effect in the repeated measures ANOVA (rmANOVA), we contrasted the different comparisons to identify the causes of the interaction effects. In Latin script, results are averaged over the levels of ‘group’. P-values are adjusted for comparing a family of 15. In Braille script, p-values are adjusted for comparing a family of 66.

| Script | Effect | Comparison | Comparison contrasted | Mean Difference | SE | t | Cohen's d | p<sub>bonf</sub> |
| --- | --- | --- | --- | --- | --- | --- | --- | --- |
| Latin | Condition | RW-PW | RW-NW | -0.221 | 0.053 | -4.187 | -1.281 | 0.001 |
|  |  |  | RW-FS | -0.183 | 0.053 | -3.476 | -1.063 | 0.014 |
|  |  |  | PW-NW | -0.121 | 0.053 | -2.291 | -0.701 | 0.378 |
|  |  |  | PW-FS | -0.179 | 0.053 | -3.397 | -1.039 | 0.017 |
|  |  |  | NW-FS | -0.217 | 0.053 | -4.108 | -1.257 | 0.002 |
|  |  | RW-NW | RW-FS | 0.037 | 0.053 | 0.711 | 0.217 | 1 |
|  |  |  | PW-NW | 0.1 | 0.053 | 1.896 | 0.58 | 0.936 |
|  |  |  | PW-FS | 0.042 | 0.053 | 0.79 | 0.242 | 1 |
|  |  |  | NW-FS | 0.004 | 0.053 | 0.079 | 0.024 | 1 |
|  |  | RW-FS | PW-NW | 0.063 | 0.053 | 1.185 | 0.362 | 1 |
|  |  |  | PW-FS | 0.004 | 0.053 | 0.079 | 0.024 | 1 |
|  |  |  | NW-FS | -0.033 | 0.053 | -0.632 | -0.193 | 1 |
|  |  | PW-NW | PW-FS | -0.058 | 0.053 | -1.106 | -0.338 | 1 |
|  |  |  | NW-FS | -0.096 | 0.053 | -1.817 | -0.556 | 1 |
|  |  | PW-FS | NW-FS | -0.037 | 0.053 | -0.711 | -0.217 | 1 |
| Braille | Grp*Cond. | experts, RW-PW | novices, RW-PW | 0.092 | 0.079 | 1.161 | 0.636 | 1 |
|  |  |  | experts, RW-NW | -0.2 | 0.087 | -2.304 | -1.387 | 1 |
|  |  |  | novices, RW-NW | 0.108 | 0.079 | 1.372 | 0.751 | 1 |
|  |  |  | experts, RW-FS | -0.283 | 0.087 | -3.264 | -1.965 | 0.116 |
|  |  |  | novices, RW-FS | 0.083 | 0.079 | 1.055 | 0.578 | 1 |
|  |  |  | experts, PW-NW | -0.233 | 0.087 | -2.688 | -1.618 | 0.602 |
|  |  |  | novices, PW-NW | 0.167 | 0.079 | 2.11 | 1.156 | 1 |
|  |  |  | experts, PW-FS | -0.317 | 0.087 | -3.648 | -2.196 | 0.035 |
|  |  |  | novices, PW-FS | 0.05 | 0.079 | 0.633 | 0.347 | 1 |
|  |  |  | experts, NW-FS | -0.3 | 0.087 | -3.456 | -2.081 | 0.064 |
|  |  |  | novices, NW-FS | -0.042 | 0.079 | -0.528 | -0.289 | 1 |
|  |  | novices, RW-PW | experts, RW-NW | -0.292 | 0.079 | -3.693 | -2.023 | 0.028 |
|  |  |  | novices, RW-NW | 0.017 | 0.061 | 0.272 | 0.116 | 1 |
|  |  |  | experts, RW-FS | -0.375 | 0.079 | -4.748 | -2.601 | < .001 |
|  |  |  | novices, RW-FS | -0.008 | 0.061 | -0.136 | -0.058 | 1 |
|  |  |  | experts, PW-NW | -0.325 | 0.079 | -4.115 | -2.254 | 0.007 |
|  |  |  | novices, PW-NW | 0.075 | 0.061 | 1.222 | 0.52 | 1 |
|  |  |  | experts, PW-FS | -0.408 | 0.079 | -5.171 | -2.832 | < .001 |
|  |  |  | novices, PW-FS | -0.042 | 0.061 | -0.679 | -0.289 | 1 |
|  |  |  | experts, NW-FS | -0.392 | 0.079 | -4.96 | -2.716 | < .001 |
|  |  |  | novices, NW-FS | -0.133 | 0.061 | -2.172 | -0.925 | 1 |
|  |  | experts, RW-NW | novices, RW-NW | 0.308 | 0.079 | 3.904 | 2.138 | 0.014 |
|  |  |  | experts, RW-FS | -0.083 | 0.087 | -0.96 | -0.578 | 1 |
|  |  |  | novices, RW-FS | 0.283 | 0.079 | 3.588 | 1.965 | 0.039 |
|  |  |  | experts, PW-NW | -0.033 | 0.087 | -0.384 | -0.231 | 1 |
|  |  |  | novices, PW-NW | 0.367 | 0.079 | 4.643 | 2.543 | < .001 |
|  |  |  | experts, PW-FS | -0.117 | 0.087 | -1.344 | -0.809 | 1 |
|  |  |  | novices, PW-FS | 0.25 | 0.079 | 3.166 | 1.734 | 0.148 |
|  |  |  | experts, NW-FS | -0.1 | 0.087 | -1.152 | -0.694 | 1 |
|  |  |  | novices, NW-FS | 0.158 | 0.079 | 2.005 | 1.098 | 1 |
|  |  | novices, RW-NW | experts, RW-FS | -0.392 | 0.079 | -4.96 | -2.716 | < .001 |
|  |  |  | novices, RW-FS | -0.025 | 0.061 | -0.407 | -0.173 | 1 |
|  |  |  | experts, PW-NW | -0.342 | 0.079 | -4.326 | -2.37 | 0.003 |
|  |  |  | novices, PW-NW | 0.058 | 0.061 | 0.95 | 0.405 | 1 |
|  |  |  | experts, PW-FS | -0.425 | 0.079 | -5.382 | -2.948 | < .001 |
|  |  |  | novices, PW-FS | -0.058 | 0.061 | -0.95 | -0.405 | 1 |
|  |  |  | experts, NW-FS | -0.408 | 0.079 | -5.171 | -2.832 | < .001 |
|  |  |  | novices, NW-FS | -0.15 | 0.061 | -2.444 | -1.04 | 1 |
|  |  | experts, RW-FS | novices, RW-FS | 0.367 | 0.079 | 4.643 | 2.543 | < .001 |
|  |  |  | experts, PW-NW | 0.05 | 0.087 | 0.576 | 0.347 | 1 |
|  |  |  | novices, PW-NW | 0.45 | 0.079 | 5.698 | 3.121 | < .001 |
|  |  |  | experts, PW-FS | -0.033 | 0.087 | -0.384 | -0.231 | 1 |
|  |  |  | novices, PW-FS | 0.333 | 0.079 | 4.221 | 2.312 | 0.004 |
|  |  |  | experts, NW-FS | -0.017 | 0.087 | -0.192 | -0.116 | 1 |
|  |  |  | novices, NW-FS | 0.242 | 0.079 | 3.06 | 1.676 | 0.203 |
|  |  | novices, RW-FS | experts, PW-NW | -0.317 | 0.079 | -4.01 | -2.196 | 0.009 |
|  |  |  | novices, PW-NW | 0.083 | 0.061 | 1.358 | 0.578 | 1 |
|  |  |  | experts, PW-FS | -0.4 | 0.079 | -5.065 | -2.774 | < .001 |
|  |  |  | novices, PW-FS | -0.033 | 0.061 | -0.543 | -0.231 | 1 |
|  |  |  | experts, NW-FS | -0.383 | 0.079 | -4.854 | -2.659 | < .001 |
|  |  |  | novices, NW-FS | -0.125 | 0.061 | -2.036 | -0.867 | 1 |
|  |  | experts, PW-NW | novices, PW-NW | 0.4 | 0.079 | 5.065 | 2.774 | < .001 |
|  |  |  | experts, PW-FS | -0.083 | 0.087 | -0.96 | -0.578 | 1 |
|  |  |  | novices, PW-FS | 0.283 | 0.079 | 3.588 | 1.965 | 0.039 |
|  |  |  | experts, NW-FS | -0.067 | 0.087 | -0.768 | -0.462 | 1 |
|  |  |  | novices, NW-FS | 0.192 | 0.079 | 2.427 | 1.329 | 1 |
|  |  | novices, PW-NW | experts, PW-FS | -0.483 | 0.079 | -6.12 | -3.352 | < .001 |
|  |  |  | novices, PW-FS | -0.117 | 0.061 | -1.901 | -0.809 | 1 |
|  |  |  | experts, NW-FS | -0.467 | 0.079 | -5.909 | -3.237 | < .001 |
|  |  |  | novices, NW-FS | -0.208 | 0.061 | -3.394 | -1.445 | 0.078 |
|  |  | experts, PW-FS | novices, PW-FS | 0.367 | 0.079 | 4.643 | 2.543 | < .001 |
|  |  |  | experts, NW-FS | 0.017 | 0.087 | 0.192 | 0.116 | 1 |
|  |  |  | novices, NW-FS | 0.275 | 0.079 | 3.482 | 1.907 | 0.055 |
|  |  | novices, PW-FS | experts, NW-FS | -0.35 | 0.079 | -4.432 | -2.427 | 0.002 |
|  |  |  | novices, NW-FS | -0.092 | 0.061 | -1.493 | -0.636 | 1 |
|  |  | experts, NW-FS | novices, NW-FS | 0.258 | 0.079 | 3.271 | 1.792 | 0.107 |

## References

Arcaro, M. J., & Livingstone, M. S. (2017). A hierarchical, retinotopic proto-organization of the primate visual system at birth. eLife, 6, e26196. 10.7554/eLife.26196

Baker, C. I., Liu, J., Wald, L. L., Kwong, K. K., Benner, T., & Kanwisher, N. (2007). Visual word processing and experiential origins of functional selectivity in human extrastriate cortex. Proceedings of the National Academy of Sciences, 104(21), 9087–9092. 10.1073/pnas.0703300104

Ben-Shachar, M., Dougherty, R. F., Deutsch, G. K., & Wandell, B. A. (2007). Differential Sensitivity to Words and Shapes in Ventral Occipito-Temporal Cortex. Cerebral Cortex, 17(7), 1604–1611. 10.1093/cercor/bhl071

Bhaya-Grossman, I., & Chang, E. F. (2022). Speech Computations of the Human Superior Temporal Gyrus. Annual Review of Psychology, 73(1), 79–102. 10.1146/annurev-psych-022321-035256

Biederman, I. (1987). Recognition-by-Components: A Theory of Human Image Understanding.

Bilalic, M., Langner, R., Ulrich, R., & Grodd, W. (2011). Many Faces of Expertise: Fusiform Face Area in Chess Experts and Novices. Journal of Neuroscience, 31(28), 10206–10214. 10.1523/JNEUROSCI.5727-10.2011

Bola, Ł., Radziun, D., Siuda-Krzywicka, K., Sowa, J. E., Paplińska, M., Sumera, E., & Szwed, M. (2017). Universal Visual Features Might Be Necessary for Fluent Reading. A Longitudinal Study of Visual Reading in Braille and Cyrillic Alphabets. Frontiers in Psychology, 8. 10.3389/fpsyg.2017.00514

Brodeur, M. B., Dionne-Dostie, E., Montreuil, T., & Lepage, M. (2010). The Bank of Standardized Stimuli (BOSS), a New Set of 480 Normative Photos of Objects to Be Used as Visual Stimuli in Cognitive Research. PLoS ONE, 5(5), e10773. 10.1371/journal.pone.0010773

Büchel, C., Price, C., & Friston, K. (1998). A multimodal language region in the ventral visual pathway. Nature, 394(6690), 274–277. 10.1038/28389

Carling, K. (2000). Resistant outlier rules and the non-Gaussian case. Computational Statistics & Data Analysis, 33(3), 249–258. 10.1016/S0167-9473(99)00057-2

Chang, E. F., Rieger, J. W., Johnson, K., Berger, M. S., Barbaro, N. M., & Knight, R. T. (2010). Categorical speech representation in human superior temporal gyrus. Nature Neuroscience, 13(11), 1428–1432. 10.1038/nn.2641

Changizi, M. A., Zhang, Q., Ye, H., & Shimojo, S. (2006). The Structures of Letters and Symbols throughout Human History Are Selected to Match Those Found in Objects in Natural Scenes. 23.

Code braille français uniformisé pour la transcription des textes imprimés (CBFU) Réalisé. (2008). 76(3), 61–64.

Cohen, L., Dehaene, S., Naccache, L., Lehéricy, S., Dehaene-Lambertz, G., Hénaff, M.-A., & Michel, F. (2000). The visual word form area: Spatial and temporal characterization of an initial stage of reading in normal subjects and posterior split-brain patients. Brain: A Journal of Neurology, 123, 291–307.

Cohen, L., Lehéricy, S., Chochon, F., Lemer, C., Rivaud, S., & Dehaene, S. (2002). Language-specific tuning of visual cortex? Functional properties of the Visual Word Form Area. Brain, 125(5), 1054–1069. 10.1093/brain/awf094

Coquillart, N., Collignon, O., Cerpelloni, F., & Van Audenhaege, A. (2022). Lecture visuelle du braille et plasticité cérébrale. Zoom sur l’aire visuelle de la forme des mots [UCLouvain]. https://dial.uclouvain.be/memoire/ucl/object/thesis:37423

Dȩbska, A., Wójcik, M., Chyl, K., Dziȩgiel-Fivet, G., & Jednoróg, K. (2023). Beyond the Visual Word Form Area – a cognitive characterization of the left ventral occipitotemporal cortex. Frontiers in Human Neuroscience, 17, 1199366. 10.3389/fnhum.2023.1199366

Dehaene, S., & Cohen, L. (2011). The unique role of the visual word form area in reading. Trends in Cognitive Sciences, 15(6), 254–262. 10.1016/j.tics.2011.04.003

Dehaene, S., Cohen, L., Morais, J., & Kolinsky, R. (2015). Illiterate to literate: Behavioural and cerebral changes induced by reading acquisition. Nature Reviews Neuroscience, 16(4), 234–244. 10.1038/nrn3924

Dehaene, S., Cohen, L., Sigman, M., & Vinckier, F. (2005). The neural code for written words: A proposal. Trends in Cognitive Sciences, 9(7), 335–341. 10.1016/j.tics.2005.05.004

Dehaene-Lambertz, G., Monzalvo, K., & Dehaene, S. (2018). The emergence of the visual word form: Longitudinal evolution of category-specific ventral visual areas during reading acquisition. PLOS Biology, 16(3), e2004103. 10.1371/journal.pbio.2004103

Eickhoff, S. B., Stephan, K. E., Mohlberg, H., Grefkes, C., Fink, G. R., Amunts, K., & Zilles, K. (2005). A new SPM toolbox for combining probabilistic cytoarchitectonic maps and functional imaging data. NeuroImage, 25(4), 1325–1335. 10.1016/j.neuroimage.2004.12.034

Fedorenko, E., Behr, M. K., & Kanwisher, N. (2011). Functional specificity for high-level linguistic processing in the human brain. Proceedings of the National Academy of Sciences, 108(39), 16428–16433. 10.1073/pnas.1112937108

Fedorenko, E., Hsieh, P.-J., Nieto-Castañón, A., Whitfield-Gabrieli, S., & Kanwisher, N. (2010). New Method for fMRI Investigations of Language: Defining ROIs Functionally in Individual Subjects. Journal of Neurophysiology, 104(2), 1177–1194. 10.1152/jn.00032.2010

Ferrand, L., New, B., Brysbaert, M., Keuleers, E., Bonin, P., Méot, A., Augustinova, M., & Pallier, C. (2010). The French Lexicon Project: Lexical decision data for 38,840 French words and 38,840 pseudowords. Behavior Research Methods, 42(2), 488–496. 10.3758/BRM.42.2.488

Frey, M., Nau, M., & Doeller, C. F. (2021). Magnetic resonance-based eye tracking using deep neural networks. Nature Neuroscience, 24(12), 1772–1779. 10.1038/s41593-021-00947-w

Gau, R., Barilari, M., Battal, C., Caron-Guyon, J., Cerpelloni, F., Falagiarda, F., MacLean, M., Mattioni, S., Rezk, M., Shahzad, I., Yang, Y., & Collignon, O. (2023). *Bidspm: An spm-centric bids app for flexible statistical analysis*. OHBM. https://dial.uclouvain.be/pr/boreal/en/object/boreal%3A279516

Gau, R., & Cabee, P. (2023). *bidsMReye* (Version 0.3.0) [Computer software]. Zenodo. 10.5281/zenodo.7574501

Gauthier, I., & Tarr, M. J. (1997). Becoming a “Greeble” Expert: Exploring Mechanisms for Face Recognition. Vision Research, 37(12), 1673–1682. 10.1016/S0042-6989(96)00286-6

Gauthier, I., Williams, P., Tarr, M. J., & Tanaka, J. (1998). Training ‘greeble’ experts: A framework for studying expert object recognition processes. Vision Research, 38(15–16), 2401–2428. 10.1016/S0042-6989(97)00442-2

Gimenes, M., Perret, C., & New, B. (2020). Lexique-Infra: Grapheme-phoneme, phoneme-grapheme regularity, consistency, and other sublexical statistics for 137,717 polysyllabic French words. Behavior Research Methods, 52(6), 2480–2488. 10.3758/s13428-020-01396-2

Gorgolewski, K. J., Auer, T., Calhoun, V. D., Craddock, R. C., Das, S., Duff, E. P., Flandin, G., Ghosh, S. S., Glatard, T., Halchenko, Y. O., Handwerker, D. A., Hanke, M., Keator, D., Li, X., Michael, Z., Maumet, C., Nichols, B. N., Nichols, T. E., Pellman, J., … Poldrack, R. A. (2016). The brain imaging data structure, a format for organizing and describing outputs of neuroimaging experiments. Scientific Data, 3(1), 160044. 10.1038/sdata.2016.44

Grill-Spector, K., Kourtzi, Z., & Kanwisher, N. (2001). The lateral occipital complex and its role in object recognition. Vision Research, 41(10–11), 1409–1422. 10.1016/S0042-6989(01)00073-6

Grill-Spector, K., & Malach, R. (2004). The human visual cortex. Annual Review of Neuroscience, 27(1), 649–677. 10.1146/annurev.neuro.27.070203.144220

Kanwisher, N., Chun, M. M., McDermott, J., & Ledden, P. J. (1996). Functional imaging of human visual recognition. Cognitive Brain Research, 5(1–2), 55–67. 10.1016/S0926-6410(96)00041-9

Kriegeskorte, N., Mur, M., & Bandettini, P. (2008). Representational similarity analysis—Connecting the branches of systems neuroscience. Frontiers in Systems Neuroscience, 2(NOV), 1–28. 10.3389/neuro.06.004.2008

Krizhevsky, A., Sutskever, I., & Hinton, G. E. (2017). ImageNet classification with deep convolutional neural networks. Communications of the ACM, 60(6), 84–90. 10.1145/3065386

Landau, B., Smith, L. B., & Jones, S. S. (1988). The importance of shape in early lexical learning. Cognitive Development, 3(3), 299–321. 10.1016/0885-2014(88)90014-7

Li, J., Osher, D. E., Hansen, H. A., & Saygin, Z. M. (2020). Innate connectivity patterns drive the development of the visual word form area. Scientific Reports, 10(1), 18039. 10.1038/s41598-020-75015-7

Malach, R., Reppas, J. B., Benson, R. R., Kwong, K. K., Jiang, H., Kennedy, W. A., Ledden, P. J., Brady, T. J., Rosen, B. R., & Tootell, R. B. (1995). Object-related activity revealed by functional magnetic resonance imaging in human occipital cortex. Proceedings of the National Academy of Sciences, 92(18), 8135–8139. 10.1073/pnas.92.18.8135

Marr, D., & Hildreth, E. (1980). Theory of edge detection.

Martin, A., Schurz, M., Kronbichler, M., & Richlan, F. (2015). Reading in the brain of children and adults: A meta-analysis of 40 functional magnetic resonance imaging studies. Human Brain Mapping, 36(5), 1963–1981. 10.1002/hbm.22749

Martin, L., Durisko, C., Moore, M. W., Coutanche, M. N., Chen, D., & Fiez, J. A. (2019). The VWFA Is the Home of Orthographic Learning When Houses Are Used as Letters. Eneuro, 6(1), ENEURO.0425-17.2019. 10.1523/ENEURO.0425-17.2019

Mattioni, S., Rezk, M., Battal, C., Bottini, R., Cuculiza Mendoza, K. E., Oosterhof, N. N., & Collignon, O. (2020). Categorical representation from sound and sight in the ventral occipito-temporal cortex of sighted and blind. eLife, 9, e50732. 10.7554/eLife.50732

Moore, M. W., Durisko, C., Perfetti, C. A., & Fiez, J. A. (2014). Learning to Read an Alphabet of Human Faces Produces Left-lateralized Training Effects in the Fusiform Gyrus. Journal of Cognitive Neuroscience, 26(4), 896–913. 10.1162/jocn_a_00506

New, B., Pallier, C., Ferrand, L., & Matos, R. (2005). Manuel de Lexique 3. Behavior Research Methods, Instruments, & Computers, 36(3), 516–524.

Oosterhof, N. N., Connolly, A. C., & Haxby, J. V. (2016). CoSMoMVPA: Multi-Modal Multivariate Pattern Analysis of Neuroimaging Data in Matlab/GNU Octave. Frontiers in Neuroinformatics, 10. 10.3389/fninf.2016.00027

Pattamadilok, C., Planton, S., & Bonnard, M. (2019). Spoken language coding neurons in the Visual Word Form Area: Evidence from a TMS adaptation paradigm. NeuroImage, 186, 278–285. 10.1016/j.neuroimage.2018.11.014

Planton, S., Chanoine, V., Sein, J., Anton, J.-L., Nazarian, B., Pallier, C., & Pattamadilok, C. (2019). Top-down activation of the visuo-orthographic system during spoken sentence processing. NeuroImage, 202, 116135. 10.1016/j.neuroimage.2019.116135

Price, C. J., & Devlin, J. T. (2003). The myth of the visual word form area. NeuroImage, 19(3), 473–481. 10.1016/S1053-8119(03)00084-3

Price, C. J., & Devlin, J. T. (2004). The pro and cons of labelling a left occipitotemporal region: “The visual word form area”. NeuroImage, 22(1), 477–479. 10.1016/j.neuroimage.2004.01.018

Price, C. J., & Devlin, J. T. (2011). The Interactive Account of ventral occipitotemporal contributions to reading. Trends in Cognitive Sciences, 15(6), 246–253. 10.1016/j.tics.2011.04.001

Rajalingham, R., Kar, K., Sanghavi, S., Dehaene, S., & DiCarlo, J. J. (2020). The inferior temporal cortex is a potential cortical precursor of orthographic processing in untrained monkeys. Nature Communications, 11(1), 3886. 10.1038/s41467-020-17714-3

Reich, L., Szwed, M., Cohen, L., & Amedi, A. (2011). A Ventral Visual Stream Reading Center Independent of Visual Experience. Current Biology, 21(5), 363–368. 10.1016/j.cub.2011.01.040

Riesenhuber, M., & Poggio, T. (1999). Hierarchical models of object recognition in cortex. Nature Neuroscience, 2(11), 1019–1025. 10.1038/14819

Roberts, D. J., Woollams, A. M., Kim, E., Beeson, P. M., Rapcsak, S. Z., & Lambon Ralph, M. A. (2013). Efficient Visual Object and Word Recognition Relies on High Spatial Frequency Coding in the Left Posterior Fusiform Gyrus: Evidence from a Case-Series of Patients with Ventral Occipito-Temporal Cortex Damage. Cerebral Cortex, 23(11), 2568–2580. 10.1093/cercor/bhs224

Rossion, B., & Pourtois, G. (2004). Revisiting Snodgrass and Vanderwart’s Object Pictorial Set: The Role of Surface Detail in Basic-Level Object Recognition. Perception, 33(2), 217–236. 10.1068/p5117

Saygin, Z. M., Osher, D. E., Norton, E. S., Youssoufian, D. A., Beach, S. D., Feather, J., Gaab, N., Gabrieli, J. D. E., & Kanwisher, N. (2016). Connectivity precedes function in the development of the visual word form area. Nature Neuroscience, 19(9), 1250–1255. 10.1038/nn.4354

Siuda-Krzywicka, K., Bola, Ł., Paplińska, M., Sumera, E., Jednoróg, K., Marchewka, A., Śliwińska, M. W., Amedi, A., & Szwed, M. (2016). Massive cortical reorganization in sighted Braille readers. eLife, 5, e10762. 10.7554/eLife.10762

Stelzer, J., Chen, Y., & Turner, R. (2013). Statistical inference and multiple testing correction in classification-based multi-voxel pattern analysis (MVPA): Random permutations and cluster size control. NeuroImage, 65, 69–82. 10.1016/j.neuroimage.2012.09.063

Stevens, W. D., Kravitz, D. J., Peng, C. S., Tessler, M. H., & Martin, A. (2017). Privileged Functional Connectivity between the Visual Word Form Area and the Language System. The Journal of Neuroscience, 37(21), 5288–5297. 10.1523/JNEUROSCI.0138-17.2017

Szwed, M., Dehaene, S., Kleinschmidt, A., Eger, E., Valabrègue, R., Amadon, A., & Cohen, L. (2011). Specialization for written words over objects in the visual cortex. NeuroImage, 56(1), 330–344. 10.1016/j.neuroimage.2011.01.073

Szwed, M., Qiao, E., Jobert, A., Dehaene, S., & Cohen, L. (2014). Effects of Literacy in Early Visual and Occipitotemporal Areas of Chinese and French Readers. Journal of Cognitive Neuroscience, 26(3), 459–475. 10.1162/jocn_a_00499

Tarr, M. J., & Bulthoff, H. H. (1998). Image-based object recognition in man, monkey and machine.

Vinckier, F., Dehaene, S., Jobert, A., Dubus, J. P., Sigman, M., & Cohen, L. (2007). Hierarchical Coding of Letter Strings in the Ventral Stream: Dissecting the Inner Organization of the Visual Word-Form System. Neuron, 55(1), 143–156. 10.1016/j.neuron.2007.05.031

Vogel, A. C., Petersen, S. E., & Schlaggar, B. L. (2012). The Left Occipitotemporal Cortex Does Not Show Preferential Activity for Words. Cerebral Cortex, 22(12), 2715–2732. 10.1093/cercor/bhr295

Xu, Y., Vignali, L., Sigismondi, F., Crepaldi, D., Bottini, R., & Collignon, O. (2023). Similar object shape representation encoded in the inferolateral occipitotemporal cortex of sighted and early blind people. PLOS Biology, 21(7), e3001930. 10.1371/journal.pbio.3001930

Xue, G., & Poldrack, R. A. (2007). The Neural Substrates of Visual Perceptual Learning of Words: Implications for the Visual Word Form Area Hypothesis. Journal of Cognitive Neuroscience, 19(10), 1643–1655. 10.1162/jocn.2007.19.10.1643

Yarkoni, T., Poldrack, R. A., Nichols, T. E., Van Essen, D. C., & Wager, T. D. (2011). Large-scale automated synthesis of human functional neuroimaging data. Nature Methods, 8(8), 665–670. 10.1038/nmeth.1635

Zhan, M., Pallier, C., Agrawal, A., Dehaene, S., & Cohen, L. (2023). Does the visual word form area split in bilingual readers? A millimeter-scale 7-T fMRI study. Science Advances.

Zhang, M., Weisser, V. D., Stilla, R., Prather, S. C., & Sathian, K. (2004). Multisensory cortical processing of object shape and its relation to mental imagery. *Cognitive, Affective*, & Behavioral Neuroscience, 4(2), 251–259. 10.3758/CABN.4.2.251

